# EEG Signal Processing to Control a Finger Rehabilitation System

**DOI:** 10.1101/2023.07.02.547366

**Authors:** Mahdi FallahTaherpazir, Mohammadbagher Menhaj, Atena Sajedin

## Abstract

This study aims to provide a comprehensive comparison for classification of Electroencephalography (EEG) signal based motor imagery, in time domain and time-frequency domain with different classifiers. We used EEG signals recorded while the subjects were imagining the movement of individual fingers, and analyzed the signals in time domain as well as using wavelet transform and Wigner transform. Our main goal is to compare different methods of feature extraction and classification as the important steps in the process of EEG signals for the Brain-Computer Interface (BCI) system. The experimental results indicate that the Support Vector Machine (SVM) method provides a better classification performance compared with other classification methods. Also, Linear Discriminative Analysis (LDA) performs as well as the SVM, after applying PCA for dimension reduction. The proposed scheme can be applied successfully to BCI systems where the amount of large data.

## Introduction

Brain–Computer Interfaces (BCIs) are systems that decode brain signals and translate them to control commands to control an external device like prosthetic and bionic devices^1,2^. This technology can make a new way of communication especially for people with motor disabilities, who cannot communicate with the external world through normal pathways like Amyotrophic Lateral Sclerosis (ALS) patients ^3^. To record brain activities for a BCI system, there are variety of approaches, including electroencephalography (EEG) ^4–7^, Electrocorticography (ECoG) ^8–10^, Electromyography (EMG) ^11–13^, Functional Magnetic Resonance Imaging (fMRI) ^14–16^, Magnetoencephalography (MEG) ^17–19^, and Near-Infrared Spectroscopy (NIRS) ^20–22^. EEG signals are widely used and very popular, because of its non-invasive nature, low cost, good temporal resolution and portability^23–28^. Although EEG signals have good temporal resolution, they suffer from low spatial resolution ^29–31^, which makes decoding these types of brain signals a challenging issue. So, processing collected EEG signals to identify brain commands is an important step in BCI systems ^32^. Motor Imagery signal is a type of EEG signal that records when the subject is imagining the movement of a limb. Most studies in motor imagery EEG based BCIs, are focused on decoding movements of large parts of the body like hands, foots and tongue ^33–36^. There are a limited number of studies that considered decoding the movement of finer parts of the body like individual fingers.

Experimental studies evaluated multiple movement related features for discrimination of individual fingers from one hand using EEG signal ^2^, and indicated the existence of features in EEG signals for individual fingers discrimination. They also used spectral features obtained from spectral Principal Component Analysis (PCA), to classify EEG signals, which yield 45.2 % accuracy. In addition, some researchers investigated the discrimination of individual fingers from one hand ^24^. They detected broadband power increase and low-frequency-band power decrease in EEG, when EEG power spectra were decomposed by PCA. They used these movement-related spectral structures and achieved the accuracy of 77.11 % over all subjects using Support Vector Machine (SVM) classifier. Some studies used Fisher’s linear discriminant criteria and power spectral density analysis to analyze the Event-Related Desynchronization (ERD) patterns and extract both band power and approximate entropy as features and reported higher accuracy of 81.32% for five subjects using a modified version of Sparse Representation-based Classification method. ^37^Another study indicated that the EEG pattern recognition method with Random Forest achieved the best accuracy of about 54%^38^. Another researchers introduced Autonomous Deep Learning (ADL) to classify five fingers to overcome the issue of subject dependency of EEG signals^39^. They used Common Spatial Pattern (CSP) extracted from the EEG signal of healthy subjects, as the input of ADL. The results on the subject-dependence classification across four subjects using 10-fold cross-validation showed that their method achieved the classification accuracy of around 77%. On the subject-independent classification, ADL outperforms CNN by resulting stable accuracies for both training and testing, different from CNN that experience accuracy degradation to approximately 50%. EEG signals have non-stationary nature and time-frequency representation of these signals can better reveal their properties^40^. Many researchers considered classification problems in EEG signals in both time domain^41,42^ and time-frequency domain^43,44^. As EEG signals are subject oriented^45^, most of the studies focused on subject-dependent classifications. But for practical purposes, like providing a reliable input signal for BCI systems, investigating subject-independent classifications are important.

This study aims to provide a comprehensive comparison for classification of EEG based motor imagery in time domain and time-frequency domain. We aimed to provide a framework to design a reliable and accurate BCI system based on EEG signals. We discussed the pros and cons of different EEG processing methods in different domains, and considered the feasibility of classification of subject-independent data. We used EEG signals of eight healthy subjects while they were performing motor imagery tasks of flexing individual fingers. We analyzed the signals in time domain and time-frequency domain, with different classifiers like SVM with polynomial and RBF kernels, Random Forest, LDA and KNN. We used different methods to represent our data in time-frequency domain like wavelet transform and Wigner-Ville time-frequency distribution. Our results indicated the feasibility of classification of individual finger movement imagery through EEG signals. Also, we revealed that classification accuracies in time domain are significantly higher than time frequency domain, while spending less time for training. SVM performed significantly higher accuracy results than the other classifiers in both domains. But, after dimension reduction of the data with PCA, accuracies of LDA classifier improved and it performed similar to SVM.

## Experimental Procedure

### A. Subjects

Participants were thirteen healthy students studying engineering and science programs in Toros University and Mersin University. They were between the age of 20 and 35, and all had been screened for the absence of psychiatric conditions, any taken medications, and contraindications to EEG. Also, participants briefed about the goals and procedures of the experiment. They are known by their aliases “Subject A” through “Subject M” ^46^.

### B. Paradigm

Each recording session lasted for 45 minutes, which was separated into three 15 minutes interaction parts with a 2 minutes rest in between, to prevent participants’ fatigue. In the rest period, participants were free to move and talk, and EEG signals were continuously recorded during resting times. Participants sat in front of a computer screen, remained motionless and stared at the center of the screen, during the interaction period. Each trial began with the presentation of a stimulus action-signal on the computer screen, which showed the motor imagery task that participants should perform which imagination of flexion of the corresponding finger up or was down. The action signal remained on screen for 1 seconds, during which time the participants implemented the indicated mental imagery once. A pause of variable duration of 1.5-2.5 seconds followed, concluding the trial. After performing each task, participants remained passive till the next action signal. So, in each interaction period of 15 minutes, approximately 300 trials could be performed ^46^.

### C. EEG Setup

EEG data were recorded via “EEG-1200 JE-921A” EEG system, including 19 EEG leads. Recording were performed by a standard 10/20 EEG cap (Electro-Cap International, USA) with nineteen bridge electrodes. Electrodes’ impedances were monitored using the impedance-check mode of the EEG-1200 system to be ≤10 kΩ with the impedance imbalance ≤5 kΩ. Modified 10/20 standard montage was used for these recordings which include nineteen 10/20 standard electrodes and two ground electrodes, called A1 and A2, on the earbuds. In addition, one bipolar lead X3 was also used for data synchronization. Reference point or “System 0 volts”, for all the recordings, as mentioned in EEG-1200’ technical manual was at 0.55*(C3 + C4) V. Sampling frequency for all recordings was 1000 Hz and no custom filtering was applied to the recorded EEG signal. A 0.53–100Hz band-pass filter (the widest choice possible in Neurofax software) was applied to the EEG recordings. Additionally, a 50 Hz notch filter was applied in the EEG-1200 hardware to reduce electrical grid interference ^46^.

## Methods

### A. Signal processing

This section describes the used and suggested methods for EEG signal processing, including preprocessing, feature extraction, and classification techniques. An overview of the stages of analysis of EEG signals is shown in **Figure 1**. First of all, in order to preprocess EEG data and remove artifacts and de-noise signals, Independent Component Analysis (ICA) technique is used. Then the signals are applied to a band-pass filter to locate the desired frequency region. The features are then extracted from the signals using a range of metrics, including Morlet Wavelet, Bump Wavelet and Winger distribution. Finally, various classifiers, including support vector machine (SVM), and K-Nearest Neighbors (KNN), Linear Discriminant Analysis (LDA), and Random Forest (RF) are employed to discriminate the features. The following subsections provide more details on each stage of the block diagram.

**Figure 1.**
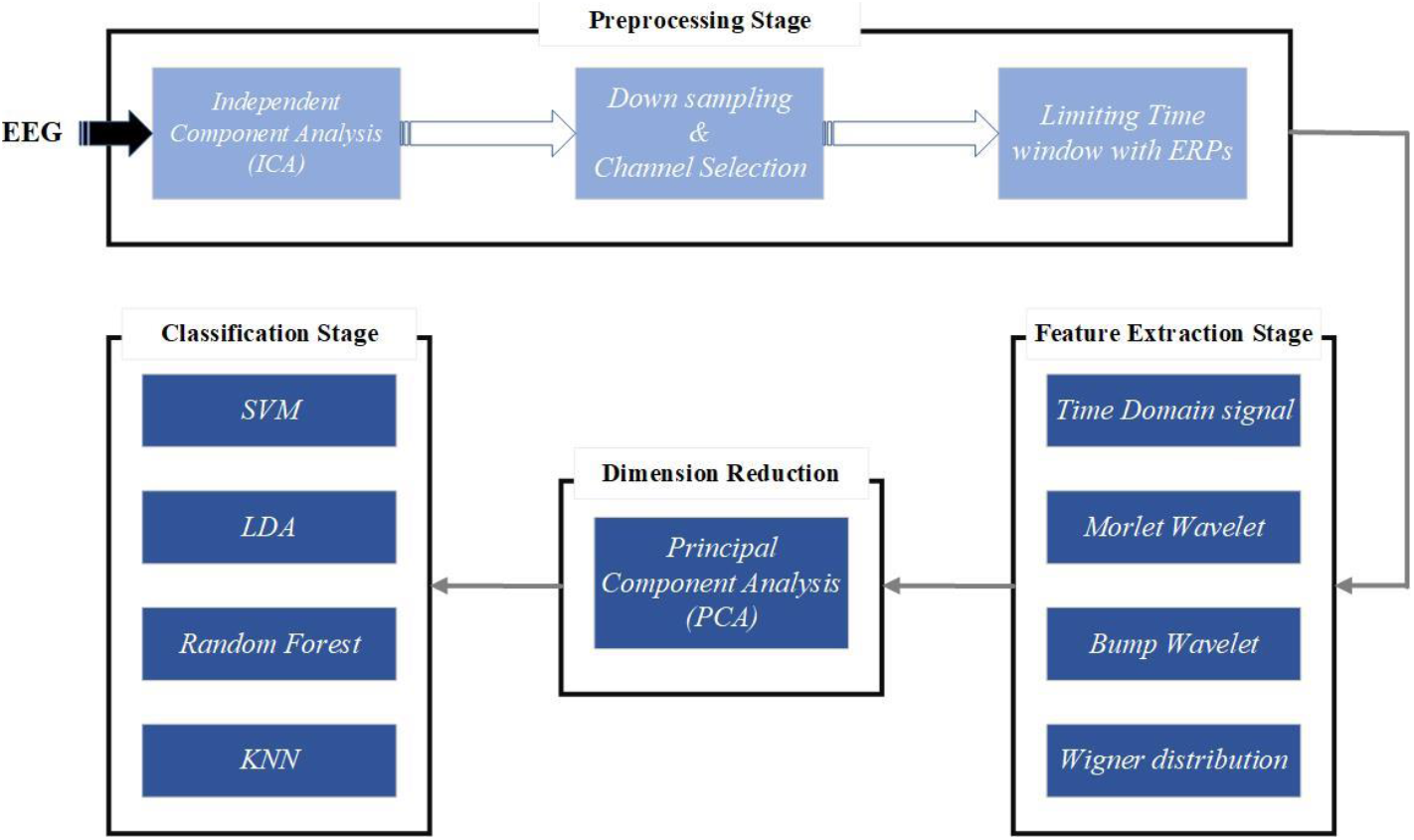
Signal Processing method. Signal processing method consists of four main stages, i) Preprocessing, ii) Feature extraction, iii) Dimension reduction and iv) Classification. The Preprocessing stage includes Independent Component Analysis as denoising step, down sampling and channel selection, in order to reduce size of the dataset, and taking advantage of ERP method to select the most relevant part of the signal. In the Feature extraction stage, the preprocessed signal is considered in two different domains, Time domain and joint Time-Frequency domain. Principal Component Analysis (PCA) is used for dimension reduction. At the final stage, the processed data are classified. After computing independent components, they should be labeled and those components that are labeled as an artifact should be removed. Independent component topographic maps and their labels are shown in Figure **2**.

- **Preprocessing Stage**: In this study, an open-source EEG datasets were used to test the proposed approaches^46^. In this stage, EEG signals are denoised using ICA algorithm. EEG signals are first applied to a high-pass filter above 2 Hz in which ICA performs notably better Signals were recorded with 1000 Hz sampling rate and through 19 EEG channels, which leads to a high dimensional data. So, in the next step, signals are down sampled to 200 Hz. In addition, only EEG signals that were recorded by the electrodes corresponding to the motor cortex are selected for further processing and analysis. The selected electrodes are shown in Figure1S. Subjects performed imagery tasks once in 1 second period after stimulus onset. The period of 100 ms before stimulus is considered as a baseline period. Event-Related Potentials (ERPs) of subject C for P3, P4, C3, C4 channels are shown in **Figure *3*** as an example. According to **Figure *3*** and Event-Related Potential (ERP) plots for other subjects and channels, it can be interpreted that the subjects started to perform the motor imageries roughly 400 ms after stimulus onset. So, the response period is considered between 400 – 900 ms after stimulus onset.

**Figure 2.**
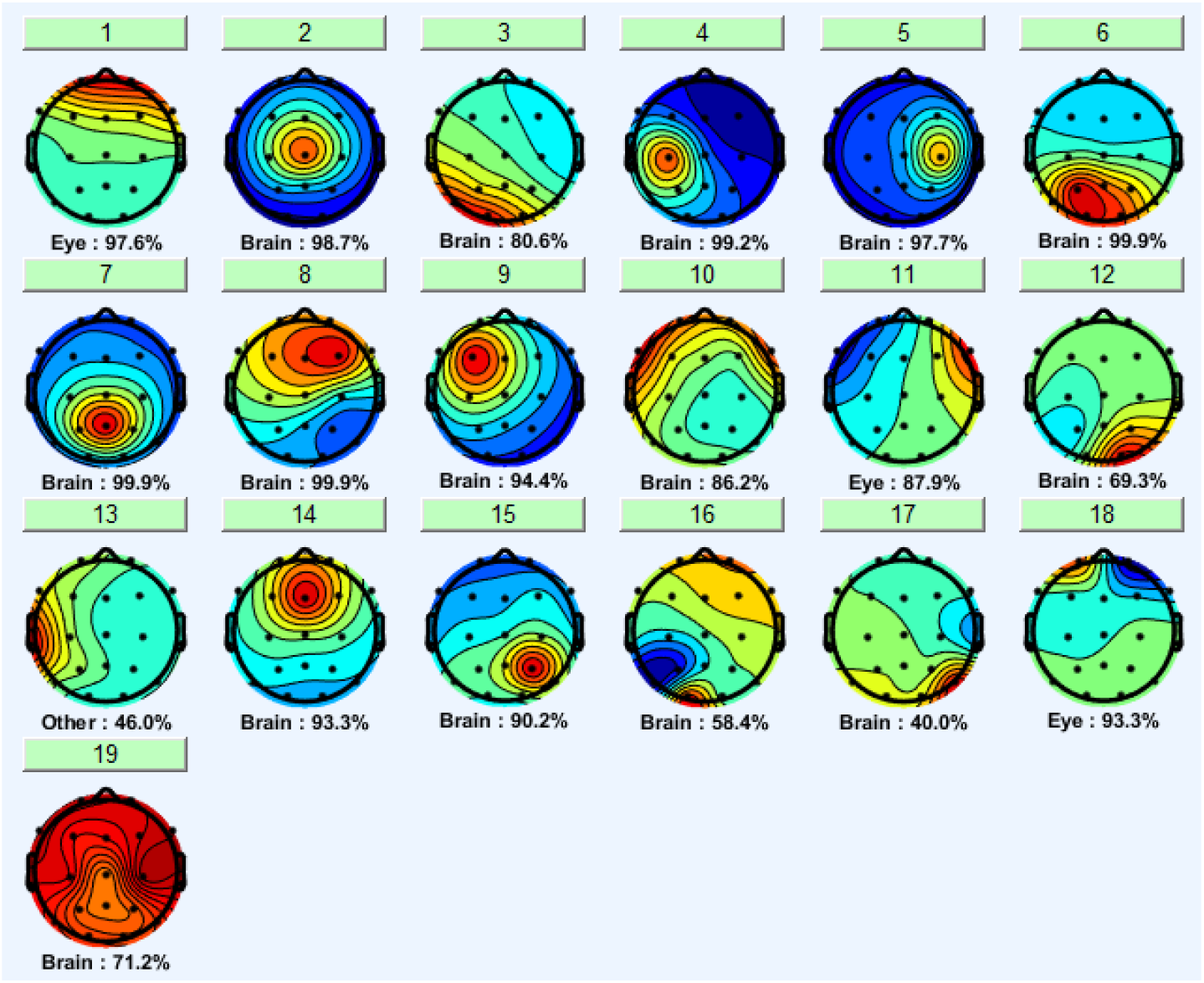
A representative Independent Components’ Topographic map for Subject C. Signals are decomposed into nineteen independent components through the ICA algorithm. Under each map, labels are shown with their percentage. These components are labeled using EEGLAB toolbox.

**Figure 3.**
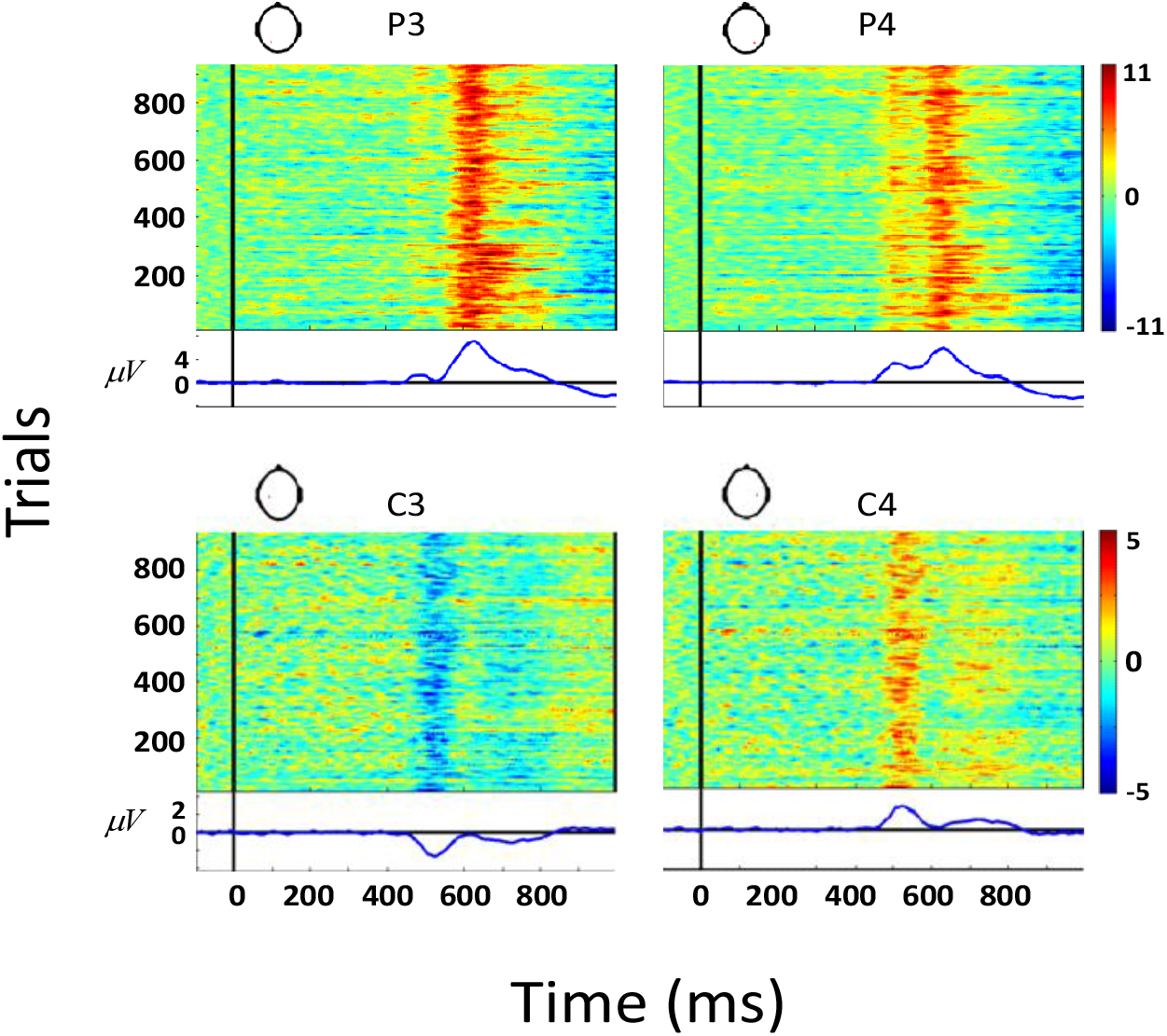
ERPs for Subject C. top: the amplitude of each trial 100 ms before the stimulus onset to 1s after stimulus onset, for each trial the baseline average amplitude is subtracted from the signal (Baseline is considered from 100 ms before stimulus onset). Event related potentials (ERPs) are shown as trial average of baseline corrected signals (down). According to these figures, the motor imagery task is performed in 400 -900 ms period.
- **Feature Extraction and Dimension Reduction Stage**: In this stage, first, time domain representation of the signals is considered as the feature vector. In addition, as EEG signals are non-stationary, and its frequency content varies over time, it is hypothesized that time-frequency representation of EEG signals can perform better^48^. Therefore, two different types of time-frequency representation of the signal, wavelet transform and Wigner-Ville distribution, are also considered as feature vectors. In wavelet transform representation two different mother wavelets are used, the Morlet wavelet which is used frequently in EEG signal processing ^49–51^, and Bump wavelet. In the preprocessing stage, dataset dimensions are reduced by selecting only six electrodes out of nineteen, and by limiting task performing period to 400 – 900 ms.

In addition to the preprocessing stage, in this stage, Principal Component Analysis (PCA) algorithm is used to reduce the dimension of the data. Classification results and training times compared after and before using PCA.

- **Classification Stage:** Four different classifiers are used in this stage: Support Vector machine (SVM) with polynomial and Radial Basis Function (RBF) kernels, Linear Discriminative Analysis (LDA), Random Forest and K-nearest neighbors (KNN).
- **Principal Component Analysis (PCA)**: Principal Component Analysis (PCA) was introduced by Pearson to find lines and planes of the closest fit to a system of points in space ^52^. This statistical technique is widely used to reduce the dimension of the datasets. The main purpose of PCA is to find the most relevant basis to transform the dataset to a new basis, in which diminishes the noise and exposes the hidden structure of dataset ^53^. PCA transforms data into a new basis in which most of the variation in data can be represented with less dimensions. In order to use PCA, following steps should be followed: a) standardize dataset, b) compute the covariance matrix of dataset, c) compute eigenvalues and eigenvectors of covariance matrix, d) select k first eigen vectors based on biggest eigenvalues, e) construct transformation matrix based on selected eigenvectors, f) transform dataset with transformation matrix.
- **Independent Component Analysis (ICA)**: Herault and Jutten introduced the Independent Component Analysis (ICA) method in order to find a linear transformation that minimizes the dependence between its components. ICA can be interpreted as an extension of PCA, which only imposes independence up to second order ^54^. ICA is one of the methods for Blind Source Separation methods (BSS) for separating data to its underlying informational components. ICA uses an assumption which is: if statistically independent signals can be extracted from signal mixture, then these extracted signals must be from different physical processes. According to this assumption, ICA separates signal mixtures into statistically independent signals ^55^. Any signal could be written as *s*_*i*_ = {*s*_*i*1_, *s*_*i*2_, … *s*_*i*N }_ where N is the number of time steps. Suppose we had two sources of signals, *s*_1_ and *s*_2_, so, we can write them as following:

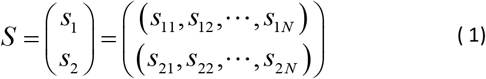

These source signals could be mixed as *x*_1_ = *as*_1_ + *bs*_2_ and *x*_2_ = *cs*_1_ + *ds*_2_ which could be rewrite as the following:

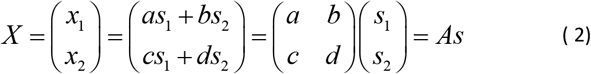

If we had the mixing matrix A, we could find the Independent Components or signal sources by inverting the above transformation. But in a source separation problem we do not have access to the mixing matrix and we only have mixed signals. For simplicity we suppose that the number of sources and the number of mixed signals is the same so the mixing matrix is a square matrix. Signal sources could be find as following:

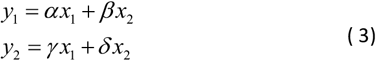

Where *α, β, γ, δ*, are called separation coefficient. The above equation could be written in the matrix form *Y* =*WX* where W is the separation matrix and it represents the inverse of the mixing matrix. Finding W is done with rotating rows of the W, so that each row has the orientation which is perpendicular to the others. In Independent Component Analysis W is first chosen randomly and then it rotated and updated iteratively. There are a variety of algorithms to find W, Infomax ICA, Jader algorithm and SOBI algorithm. In this study ICA is done with the EEGLAB toolbox and Infomax ICA algorithm.
- **Wavelet Transforms:** Wavelet transforms provide a powerful tool for the analysis of non-stationary signals like EEGs, compared to Short-Time Fourier Transform (STFT) or Gabor transform. Wavelet transform adjusts its analysis window length according to frequency, instead of using a fixed length analysis window like STFT ^56^. Wavelet transform of signal *s* (*t*) calculated from following equation:

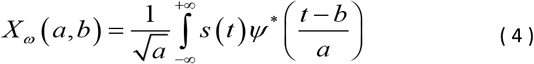

Where *ψ* is mother Wavelet. In this study two kinds of wavelet transforms i.e., Morlet and Bump, are used to transform time domain data to time-frequency domain to see how these representations affect decoding accuracies. Mother wavelets for these two transforms are displayed in **Figure *4***.

**Figure 4.**
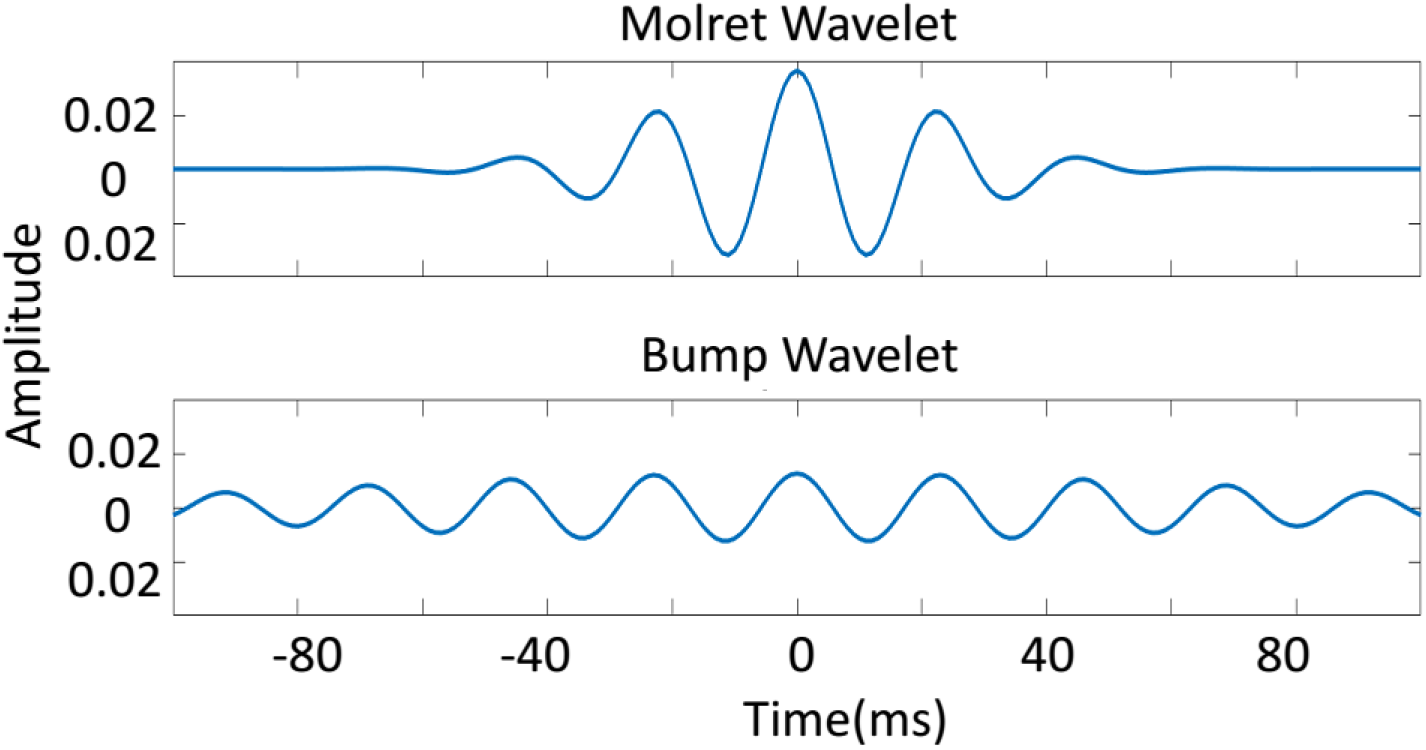
Morlet and Bump Mother Wavelets ^57^. (Up): Morlet wavelet is created from a Complex exponential and a Gaussian window and is closely related to human perception. Morlet wavelets are Gaussian shaped in the frequency domain, so they minimize the ripple effects that can be misinterpreted as oscillations. Morlet transforms also preserve the temporal resolution of the signal ^58^. (Down): Bump wavelet is bandlimited so it has a better frequency resolution than Morlet wavelet ^59^. Vertical axis shows the wavelet’s amplitude. Time is shown on the horizontal axis.
- **Wigner-Ville Distribution:** One of the most interesting and popular time frequency representations is Wigner-Ville distribution that was introduced by Wigner in ^60^ in quantum mechanics. For a real signal S(t) Wigner-Ville distribution is defined as follow:

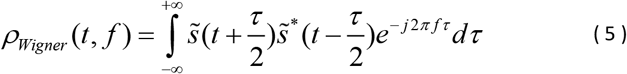

Where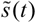 is an analytical signal where its imagery part is Hilbert transform of its real part S(t).
- **Support Vector Machine (SVM):** Support Vector Machine is a supervised learning algorithm that is used for data classification and regression. First, it transforms the input data to a high dimensional feature space with a non-linear transformation (Kernel). Then, construct a decision surface in the new space ^61^. In this study, two different kernels are used, polynomial and Radial Basis Function (RBF). SVM is primarily a two-class classifier but K SVM can be used to classify a K-class data, each SVM for separate one class from other K-1 ^62^.
- **Random Forest:** Classification accuracies can remarkably improve with growing an ensemble of trees and letting them vote for the most popular class ^63^. Random Forest is an ensemble learning method used for classification and regression ^63,64^. It creates a set of decision trees from a randomly selected subset of the training dataset and for prediction it collects votes from different decision trees. In fact, this method combines Breiman’s bagging sampling approach ^65^, and the random selection of features, introduced independently by ^66^ and ^67^, in order to construct a collection of decision trees with controlled variation ^64^.
- **Linear Discriminative Analysis (LDA):** Linear Discriminative Analysis is a dimension reduction algorithm that also is used in supervised classification ^68^. The goal of LDA classifier is to find a hyperplane so that the projections of each class’s mean on this hyperplane have minimum intraclass and maximum between class distances ^69^.
- **K-Nearest neighbors (KNN):** K Nearest neighbors is a supervised classification algorithm that predicts the label of a point, based on labels of its nearest points ^70^.
- **Dummy Classifier:** Dummy classifier specifies a chance level classification when the dataset is imbalanced and includes a different number of trials for each class. These kinds of classifiers use a strategy to classify input data. The strategy used in this study for the dummy classifiers is ‘Most Frequent’ which means that the dummy classifier corresponds each input to the label which has the most frequency in the input dataset no matter what the input is.
- **K-fold Cross Validation:** In this method, the dataset randomly is divided to K equal parts. The Model is trained with K-1 parts of the dataset. Predictions of the model evaluate on the remaining one part. By respectively using this approach so that, each one of K parts used in training and testing process, optimal model can be chosen ^71^.

## Results

In this study, we provided a comprehensive comparison of different methods for classification of EEG signals. In addition, we investigated the possibility of decoding motor imagery of the individual fingers taking advantage of the EEG signals. Due to the importance of subject-independent classification for practical purposes, the subject-independent classification has also been conducted. In the preprocessing stage, all signals are filtered and the features are then extracted and classified. Independent Component Analysis (ICA) is used to reduce artifacts in data. To select the most relevant parts of the data, ERP method is used. Classification stage was done in both time domain and time-frequency domain. Time-frequency representation of the signal is obtained with different methods like Morlet wavelet, Bump wavelet and Wigner-Ville distribution. Moreover, in order to investigate dimension reduction effects on classifiers performance, classification stage was done in two scenarios (Before PCA/After PCA). To identify the chance level, dummy classifiers with the “most frequent” strategy are used.

### A. Which Classifiers perform better in time domain?

Many experimental studies revealed that analysis of non-stationary signals, like EEG, separately in time domain, or in frequency domain cannot provide a complete representation of the data, and led to information loss^72–74^. Time-Frequency analysis resolves this problem ^75^. In order to study the impact of Time-Frequency analysis on the classification performances, we classified EEG signals in both time domain and time-frequency domain. Our results showed that the used classifiers performed better in the time-domain. For time-domain classification, before PCA application, all classifiers significantly won the dummy classifier for all subjects except 2 Subjects (Subject F and H) (see **Figure 5A** and Table S1). In the cases of Subject F and Subject H only LDA and SVM (both poly and RBF kernel) classifiers had significantly higher accuracies than dummy classifiers (5×2 Cross-Validation Test; P< 0.01; Table S1).

**Figure 5.**
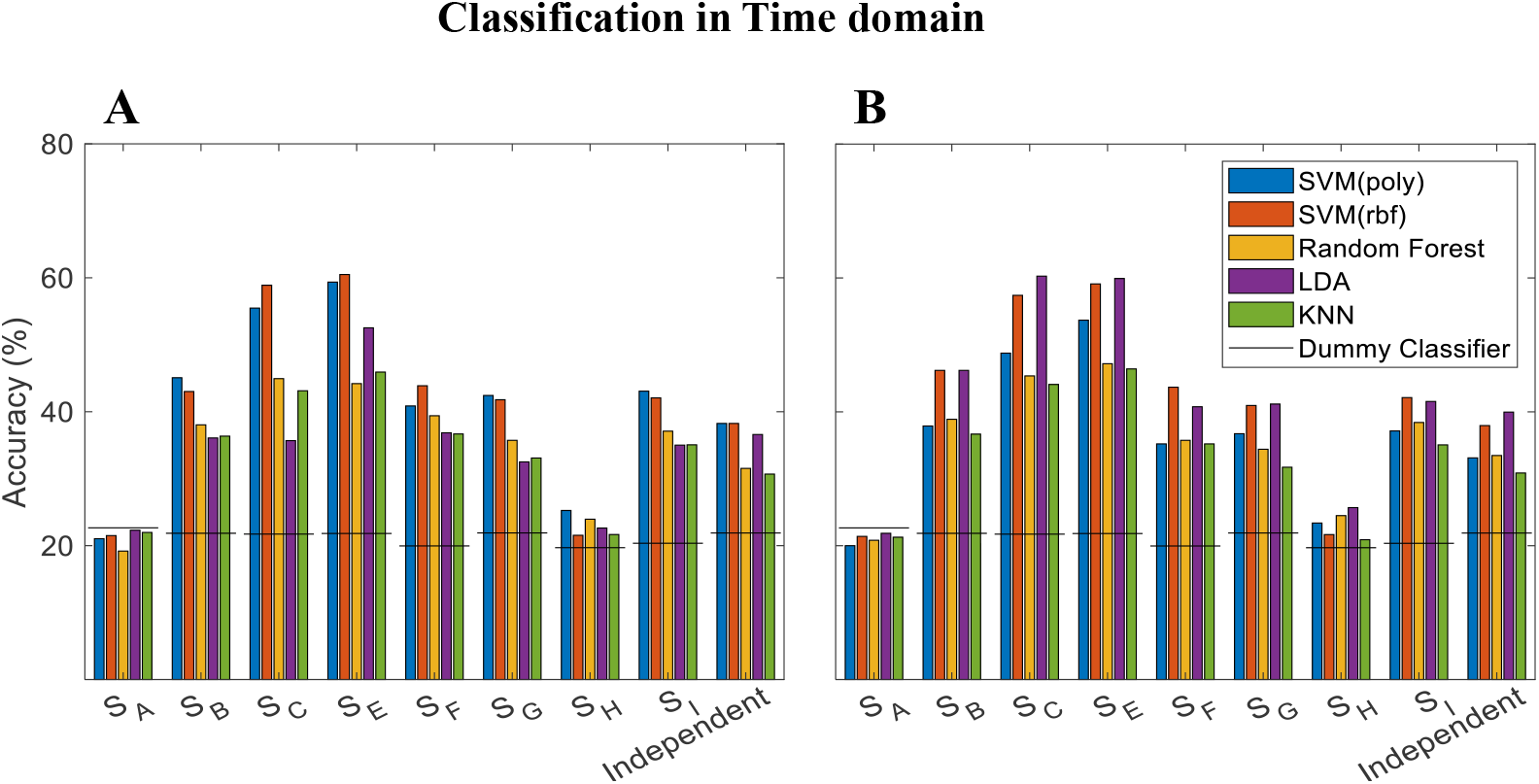
Classification in time domain. Vertical axis shows accuracies of classifying five finger motor imageries for different subjects and classifiers. Different subjects are shown in the horizontal axis and various classifiers are specified with different colors. Horizontal lines are representing the accuracies of the dummy classifier belongs to each subject. (A): Classification accuracy in time domain before dimension reduction. All classifiers performed better than dummy classifier, except in the case of subject A, which none of the classifiers reached the accuracy of dummy classifier about 22.66%. Subject C (58.89% by SVM with RBF kernel) and subject E (60.49% by SVM with RBF kernel) obtained the most accurate result. Also, SVM classifiers (with both kernels) performed the best classification accuracies among the other classifiers. (B): Classification accuracies for time domain signal after dimension reduction. In this case the 600-dimension feature vector is reduced to a 100-dimension feature vector using PCA algorithm. SubjectC (60.27%; by the LDA) and subjectE (59.92% by the LDA) have the most accurate results.

After PCA application all the classifiers showed significantly higher accuracies than the dummy classifier (5×2 Cross-Validation Test; P < 0.001; **Figure 5B** and Table S2). In time-frequency classification through Morlet wavelet, with or without PCA application, only for Subject C and Subject E, all of the classifiers significantly won the dummy classifier (5×2 Cross-Validation Test; P < 0.0001; **Figure 6A, C** and Table3S, 4S). However, after using Morlet wavelet, none of them won the dummy classifier, for Subject H (5×2 Cross-Validation Test; Minimum P > 0.065; Table S3, S4). Instead, with Wigner-Ville transform, all of the classifiers were able to significantly outperform the dummy classifier for Subjects C and G (Before PCA; 5×2 Cross-Validation Test; P < 0.005; **Figure *7*A**; Table S5) and Subjects C and E (After PCA; 5×2 Cross-Validation Test; P < 0.001; **Figure *7*B**; Table S6). In addition, like the Morlet wavelet, we observed that none of the classifiers could achieve significantly better performances than the dummy classifier for Subject H (before and after PCA). In Bump wavelet case, none of the classifiers could won the dummy classifier significantly (**Figure 6B, D**).

**Figure 6.**
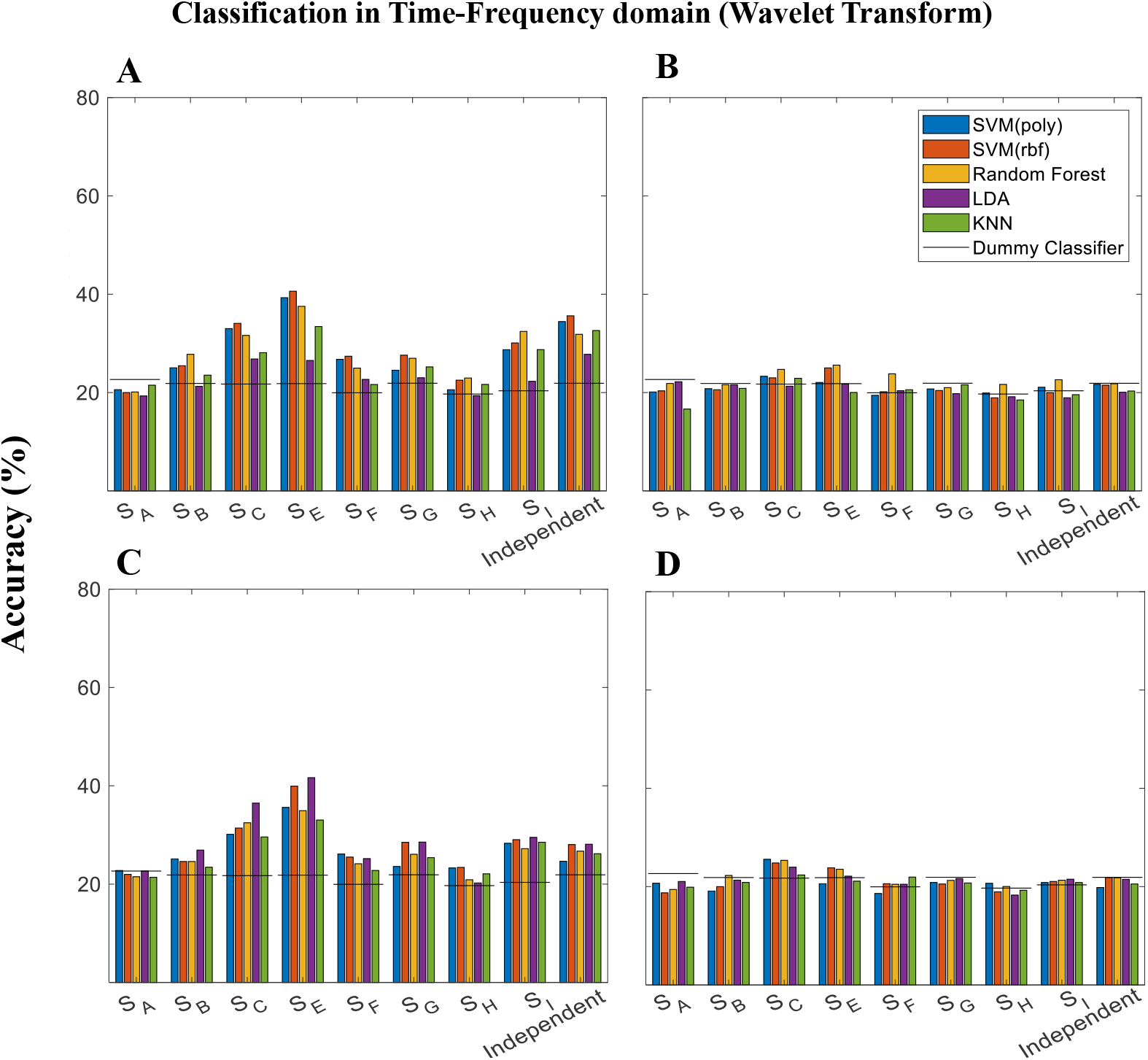
Classification in Time-Frequency domain (Wavelet transform). Vertical axis shows accuracies of classifying five finger motor imageries for different subjects and classifiers. Different subjects are shown in the horizontal axis and various classifiers are specified with different colors. Horizontal lines are representing the accuracies of the dummy classifier belongs to each subject. (A): Classification accuracies for time-frequency domain signal before dimension reduction. Time-frequency representation of the signal is achieved with Morlet wavelet transform. In this case, our feature vector is a 22800-dimension vector. Except for subject A, which none of the classifiers reached the accuracy of dummy classifier about 22.66%, for all the other subjects, almost all classifiers performed better than dummy classifier. Among all subjects, subjectC (with 34.08% accuracy for SVM with RBF kernel) and subjectE (with 40.61% accuracy for SVM with RBF kernel) have more accurate results. (B): Classification accuracies for time-frequency domain signal before dimension reduction. Time-frequency representation of the signal is achieved with Bump wavelet transform. The feature vector is a 13800-dimension vector. In this case, almost none of the classifiers could perform better than the dummy classifier. (C): Classification accuracies for time-frequency domain signal after dimension reduction via PCA. Time-frequency representation of the signal is achieved with Morlet wavelet transform. In this case, our feature vector is transformed from 22800 dimensions to a 100-dimension vector with PCA dimension reduction algorithm. Except subject A, which none of the classifiers reach the accuracy of dummy classifier about 22.66%, for all other subjects, almost all classifiers perform better than dummy classifier. (D): Classification accuracies for time-frequency domain signal after dimension reduction with principal component analysis (PCA). Time-frequency representation of the signal is achieved with Bump wavelet transform. In this case, our feature vector is transformed from 13800 dimensions to a 100-dimension vector with PCA dimension reduction algorithm.

We observed that all the classifiers perform better than the dummy classifier in the time domain. But for a complete comparison, classification accuracies are compared between time domain and time-frequency domain (Wigner-Ville transform). Our results showed that the used classifiers had significantly higher accuracies in time domain for all subjects (One-way ANOVA test; P < 0.001), except Subject H. Mean accuracies of 10-fold cross validation in both domains and results of ANOVA statistical test are reported in Table 13S.

We then compared training time between two domains. **Table *1*** shows the training time of all classifiers for Subject E in both domains. Except for the Random Forest classifier, all classifiers terminated longer training time in the time-frequency domain. Training time for all individual subjects are reported in Table 7S-12S. These results show that although time-frequency representation of data captures complete information of the signal, it does not necessarily mean that classifiers perform better in this domain.

### B. SVM or LDA?

In order to find the best classifier, “5×2 Cross-Validation Test” was conducted and all classifiers were compared in pairs in all domains (see Table S1-S6). Our findings demonstrated that SVM yield significantly higher classification accuracies than other classifiers in time domain before PCA application (see Table S1). In the time domain before PCA application SVM significantly won other classifiers in Subject B, C, E and G (5×2 Cross-Validation Test; P < 0.005; Table S1). For Subject I, SVM outperformed KNN (5×2 Cross-Validation Test; P-Value < 0.05, Table S1). SVM also significantly outperformed the dummy classifier for Subject H, (5×2 Cross-Validation Test; P < 0.05; Table S1). In addition, for Subject F, SVM showed significantly better performance than LDA (5×2 Cross-Validation Test; P < 0.004; Table S1), but there is no significant difference among other classifiers for this subject (5×2 Cross-Validation Test; P > 0.5; Table S1). There was no significant difference between polynomial and RBF kernel in SVM classifier (5×2 Cross-Validation Test; P > 0.05; Table S1), except for Subject C which RBF kernel significantly outperformed the polynomial kernel (5×2 Cross-Validation Test; P < 0.004; Table S1).

We found that although SVM is the best classifier before PCA application for time domain classification, it has a strong competitor in after PCA application. For Subject B, C, E and G, after PCA application, LDA gave significantly better performance than the other classifiers except SVM (5×2 Cross-Validation Test; P < 0.05; Table S2). According to our results, there was no significant difference between these two classifiers after PCA application (5×2 Cross-Validation Test; P > 0.05; Table S2). So, in order to find the best classifier in time domain we compared training time of these two classifiers for different subjects. Results of this comparison are reported in **Table *2*** and illustrated that LDA needs considerably shorter training time which is an advantage compared to SVM. The results of “5×2 Cross-Validation Test” for time-frequency domain are tabulated in Table S3-S6. Despite the time domain, in the time-frequency domain there was no meaningful significant difference between the classifiers (5×2 Cross-Validation Test; P > 0.05).

### C. Dimension reduction improves LDA performance

One of the difficulties we usually encounter in signal classifications, especially in EEG classification is the high dimension of the data. Dimension reduction is an important step to resolve this problem ^76^. However, reducing the dimension of data, might remove relevant information in data. So, in order to investigate the dimension reduction effects, in this study, classifications are done in two distinct scenarios. In the first scenario (Before PCA) data was classified without dimension reduction. While in the second scenario (After PCA), data dimension was reduced using the PCA algorithm before the classification stage. In order to better visualize, classification accuracy bar plots for Before PCA and After PCA scenarios are shown in **Figure 5**-7. We observed that there is no considerable difference for most of the classifiers except for LDA. LDA accuracies considerably increased after dimension reduction by PCA (One-way ANOVA; P < 0.001; Table S14-16). Statistical analysis that reported in Table S14-16, revealed that dimension reduction with PCA caused significant increase of classification accuracy for LDA in most of the subjects.

### D. Independent Case

Due to the importance of the Subject-Independent classification for BCI systems, in the current study the Subject-Independent classification is also considered. Our findings show the feasibility of Subject-Independent decoding of EEG signals from individual finger motor imagery. According to our results (see **Figure 5** and Table 1S, 2S), all of the classifiers were able to significantly outperform the dummy classifier in the time domain classification (with or without PCA application; 5×2 Cross-Validation Test; P < 0.5; Table 1S, 2S). Our results also revealed that SVM and LDA have the best performance among other classifiers in before PCA scenario (5×2 Cross-Validation Test; P < 0.05; Table 1S). Like the Subject-Dependent case, LDA yielded significantly higher accuracies than the other classifiers after PCA application (5×2 Cross-Validation Test; P < 0.005; Table 2S). In the time-frequency classification (see **Figure 6** and Figure 7), Except for the Bump wavelet, almost all of the classifiers gave significantly higher accuracies than the dummy classifier (5×2 Cross-Validation Test; P < 0.05; Table 3S-6S). But there was no meaningful significant difference among the classifiers (5×2 Cross-Validation Test; P > 0.1; Table 3S-6S).

**Table 1.** Mean Training Times of all classifiers for Subject E before PCA application

| Classifier | Time Domain(sec) | Time-Frequency Domain<br>(Wigner-Ville) (sec) |
| --- | --- | --- |
| SVM (poly) | 2.729 | 441.86 |
| SVM (RBF) | 1.864 | 452.789 |
| Random Forest | 0.0 | 0.0 |
| LDA | 0.418 | 169.06 |
| KNN | 0.0 | 14.515 |

**Table 2.** Mean Training Time (Second) for after PCA scenario in time domain classification.

| Subjects | Classifier |  |
| --- | --- | --- |
|  | SVM (RBF) | LDA |
| Subject B | 0.35 | 0.02 |
| Subject C | 0.10 | 0.01 |
| Subject E | 0.62 | 0.03 |
| Subject F | 0.12 | 0.01 |
| Subject G | 0.40 | 0.03 |
| Subject H | 0.15 | 0.01 |
| Subject I | 0.40 | 0.01 |

**Figure 7.**
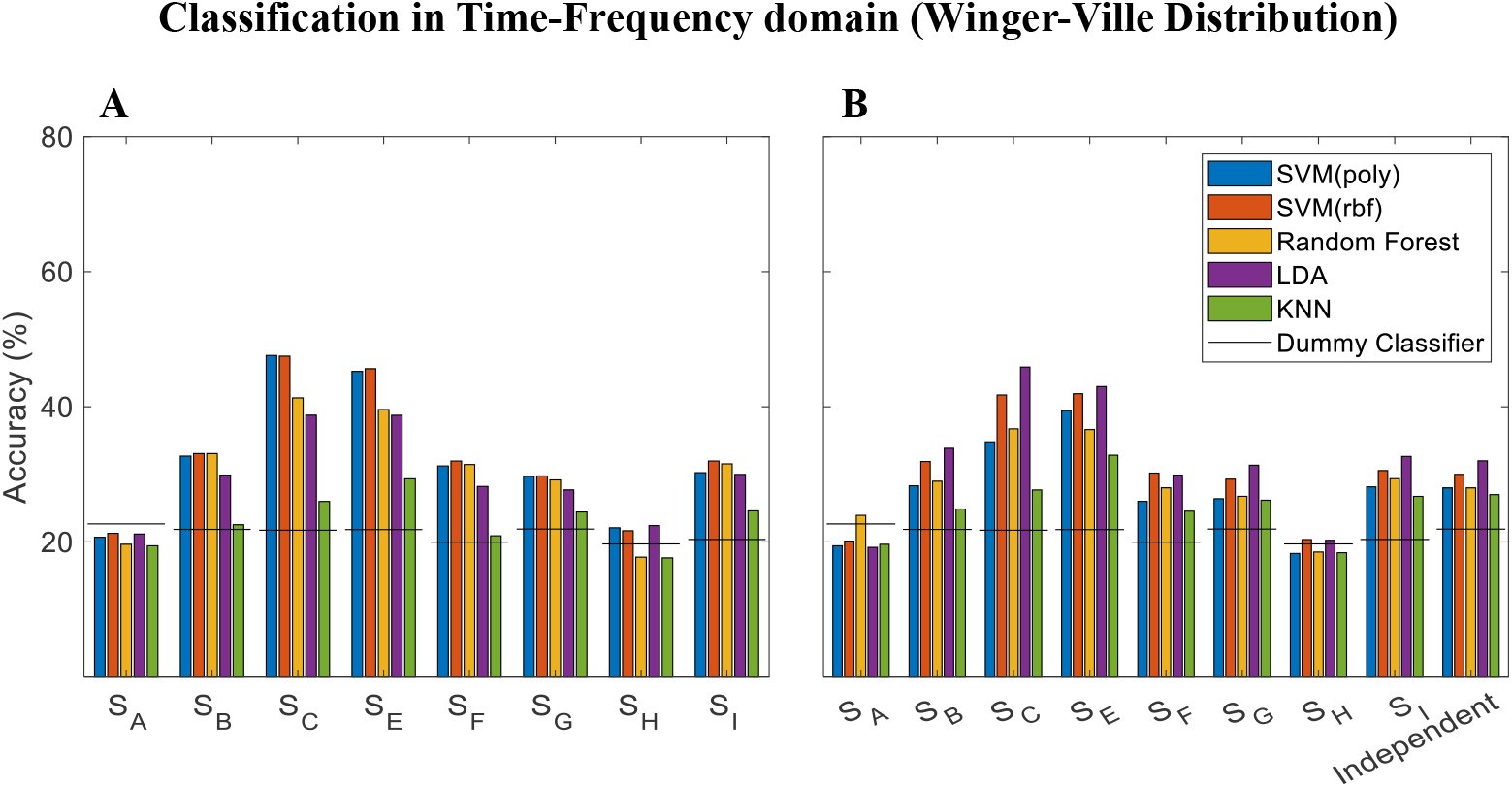
Classification in Time-Frequency domain (Wigner-Ville Distribution). Vertical axis shows accuracies of classifying five finger motor imageries for different subjects and different classifiers. The subjects are shown in horizontal axis and the classifiers are specified with different colors. The Horizontal lines are accuracies of the dummy classifier. (A): Classification accuracies before dimension reduction. Except subject A, none of the classifiers reached the accuracy of dummy classifier about 22.66%. For all other subjects, almost all classifiers performed better than dummy classifier. Among all subjects, subjectC (with 47.50% accuracy for SVM with RBF kernel) and subjectE (with 45.64% accuracy for SVM with RBF kernel) reached more accurate results. In this case, SVM has the best performance of almost all of the subjects. (B): Classification accuracies after dimension reduction with PCA. In this case, our feature vector is transformed from 120000 dimensions to a 150-dimension vector with PCA dimension reduction algorithm. Subject A, who’s never reached to the dummy classifier accuracy of 22.66%, reached 23.93% accuracy with random forest classifier.

## Discussion

Brain-Computer Interfaces (BCIs) systems and cognitive architectures are suggest helpful ways of communication with the world for people, and can improve their quality of life^77,78^. BCIs interpret the person’s brain signal to the person’s intention or mental states. Therefore, an important step in a BCI system is providing the most accurate brain signals. In this study, we investigated the feasibility of decoding a person’s intention from EEG signals that are recorded during the motor imagery of individual finger. Different methods for signals processing and classification are used and discussed. EEG signals are studied in both time domain and timefrequency domain. In preprocessing step, ICA is used in order to reduce the noise of the signal. Relevant EEG channels are also selected according to the Movement homunculus map. Time-frequency representation of the signal is achieved with different methods like: Morlet wavelet transformation, Bump wavelet transformation and Wigner-Ville transformation. Then signals are classified with different machine learning methods like SVM, LDA, Random Forest and KNN. Classification is done in two different scenarios, with and without dimension reduction. We found that, despite the power of time-frequency representations to reveal features of the non-stationary signals, classification in the time domain gives better accuracy and needs less time for training. SVM significantly yielded the best performance among different classifiers in the first scenario (before dimension reduction; 5×2 Cross-Validation Test; P < 0.01; Table 1S) and LDA performed as well as SVM in second scenario (after dimension reduction; 5×2 Cross-Validation Test; P < 0.005; Table 2S).

Although EEG signals have non-stationary nature and expected to have better classification performance in time-frequency domain ^79^, this study showed that classification in the time domain has better accuracy compared to the time-frequency domain. This can be due to the high dimension of the data set in the time-frequency domain. In the current study, the dataset in time domain was a 600-dimension vector which converted to a 120000-dimension vector after it transformed to the time-frequency representation using Wigner-Ville distribution which had best performance in time-frequency domain. We used PCA for dimension reduction and speed up the classification process. One of the important advantages of PCA is its ability to reduce the dimension without much loss of information^80^. Although, for most of the classifiers in this study PCA did not improve the classification accuracy significantly, but it reduced the training times. Moreover, comparing the classification results we revealed that LDA classifier performed significantly better after dimension reduction using PCA (Table S7, S8, Table S9, S10, Table S11, S12). For LDA, the intraclass covariance matrix can become singular due to high dimensionality of input data in comparison with low number of training vector^81^. So, this improvement in classification accuracies can be justified with the fact that dimension reduction with PCA that helps to prevent the intraclass covariance matrix to become singular, and improves the classification accuracies, especially for datasets which have low training samples ^81,82^.

We found that SVM significantly performed in all domains. SVM performed better in non-linear and high-dimension pattern recognition problems and revealed a good generalization even when dealing with low training samples ^83,84^. This is because SVM aims to find the hyperplane that classifies the data correctly and keeps the margin maximum. Maximizing the margin also keep the Vapnik and Chervonenkis dimension small, and the larger margin lead to a better generalization, and prevent the classifier from overfitting in high-dimensional datasets^85,86^. For more details please see ^86^.

In summary, we illustrated the possibility of classification of individual finger motor imagery from EEG signals through a comprehensive study and comparison between different methods for representation and classification of EEG signals. The current study provides a good perspective for the future studies for signal processing and design Brain-Computer Interfaces. Our results indicated that despite expectations based on the fact that time-frequency representations better reveal features of non-stationary signals like EEG, classification in the time domain has the advantage of lower data dimensionality and gives significantly higher accuracies and jointly less training times. Results of our study show the advantage of using PCA+LDA for classification. Dimension reduction with PCA before classification with LDA significantly improves accuracy and causes LDA to perform as good as SVM.

