## Supplementary Data for "EEG Signal Processing to Control a Finger Rehabilitation System"

### STATISTICAL ANALYSIS

K-fold Cross Validation, calculated mean accuracies of different classifiers and the best classifier can be found by comparing these accuracies. But it is hard to find out if these accuracies are real or are just due to statistical accident. To resolve this problem, statistical significance tests are designed. Doing a statistical significant test for a classification problem is not as easy as it seems.

At first glance, we could use the 10-fold cross validation method to evaluate classifiers on exactly the same split of the data. Because each classifier evaluated on the same 10 test sets, this would give samples of paired measures between two classifiers. Then we could select and use the paired Student's t-test to check if the difference in the mean accuracy between the two models is statistically significant or is just due to statistical accident. Hundreds of published papers used this method named "Cross-Validated Paired t-test" to compare their classifiers[1].

In this method there is an overlap between training sets on each of the trials. So, the accuracies are not independent and it is in contradiction with the initial assumptions of the Paired t-test. To resolve this problem[2], proposed a new method named "5x2 Cross-Validation Test". The idea of this method is to run a 2-fold cross validation 5 times. In this case, training sets do not have overlap and generate 10 different estimations. In each of the 5 iterations, data split to two parts (50% training and 50% test data) and classifiers fit to the training set and evaluate their performance on the test set. Then the training and test sets rotated and performance measures calculated again. This gives two performance difference measures as follows:

$$p^{(1)} = p_A^{(1)} - p_B^{(1)}$$
$$p^{(2)} = p_A^{(2)} - p_B^{(2)}$$

Where $p_A$ and $p_B$ are performance measures of classifiers A and B respectively. Then, we estimate the estimate mean and variance of the differences:

$$\bar{p} = \frac{p^{(1)} + p^{(2)}}{2}$$

$$s^2 = \left(p^{(1)} - \bar{p}\right)^2 + \left(p^{(2)} - \bar{p}\right)^2$$

The variance of the difference is computed for the 5 iterations and then used to compute the t statistic as follows:

$$t = \frac{p_1^{(1)}}{\sqrt{\frac{1}{5}\sum_{i=1}^{5} s_i^2}}$$

Where $p_1^{(1)}$ is the $p^{(1)}$ from very first iteration[3]. In this study "MLxtend"[4], which is a data science python library, is used to do "5x2 Cross-Validation Test".

**Table 1S**, “5x2 Cross-Validation Test” in Time Domain, Before PCA scenario.
SVM (kernel='poly', degree=3, C=100000),
SVM(kernel='rbf', gamma=0.001, C=100),
RandomForestClassifier(max_depth=5, random_state=20),
KNeighborsClassifier(n_neighbors = 60).

| **Subjects** | **Classifiers** | **Accuracies %** | **t Statistics** | **P-Value** |
|---|---|---|---|---|
| Subject B | SVM (poly)<br>SVM (RBF) | 40<br>44 | -2.413 | 0.061 |
| | SVM (poly)<br>Random Forest | 40<br>36 | 0.537 | 0.614 |
| | SVM (poly)<br>LDA | 40<br>30 | 3.754 | *0.013 |
| | SVM (poly)<br>KNN | 40<br>34 | 1.259 | 0.264 |
| | SVM (RBF)<br>Random Forest | 44<br>36 | 3.446 | *0.018 |
| | SVM (RBF)<br>LDA | 44<br>30 | 6.675 | **0.001 |
| | SVM (RBF)<br>KNN | 44<br>34 | 4.231 | **0.008 |
| | Random Forest<br>LDA | 36<br>30 | 3.578 | *0.016 |
| | Random Forest<br>KNN | 36<br>34 | 1.054 | 0.340 |
| | LDA<br>KNN | 30<br>34 | -2.386 | 0.063 |
| | Dummy<br>SVM (poly) | 22<br>40 | -8.965 | ***0.000 |
| | Dummy<br>SVM (RBF) | 20<br>44 | -17.312 | ***0.000 |
| | Dummy<br>Random Forest | 21<br>37 | -8.328 | ***0.000 |
| | Dummy<br>LDA | 20<br>30 | -3.277 | *0.022 |
| | Dummy<br>KNN | 21<br>35 | -4.794 | **0.005 |
| Subject C | SVM (poly)<br>SVM (RBF) | 51<br>54 | -5.152 | **0.004 |
| | SVM (poly)<br>Random Forest | 51<br>44 | 4.777 | **0.005 |
| | SVM (poly)<br>LDA | 51<br>31 | 5.435 | **0.003 |
| | SVM (poly)<br>KNN | 51<br>42 | 1.760 | 0.139 |
| | SVM (RBF)<br>Random Forest | 54<br>43 | 9.477 | ***0.000 |
| | SVM (RBF)<br>LDA | 54<br>31 | 6.117 | **0.002 |
| | SVM (RBF)<br>KNN | 54<br>40 | 2.890 | *0.034 |
| | Random Forest<br>LDA | 43<br>31 | 3.886 | *0.012 |
| | Random Forest<br>KNN | 43<br>41 | -0.357 | 0.736 |
| | LDA<br>KNN | 32<br>42 | -5.522 | **0.003 |
| | Dummy<br>SVM (poly) | 20<br>51 | -13.022 | ***0.000 |
| | Dummy<br>SVM (RBF) | 20<br>53 | -15.510 | ***0.000 |
| | Dummy<br>Random Forest | 20<br>43 | -14.033 | ***0.000 |
| | Dummy | 20 | -4.374 | **0.007 |

| | | | | |
|---|---|---|---|---|
| | LDA | 31 | | |
| | Dummy<br>KNN | 20<br>41 | -9.896 | ***0.000 |
| Subject E | SVM (poly)<br>SVM (RBF) | 55<br>58 | -2.229 | 0.076 |
| | SVM (poly)<br>Random Forest | 55<br>43 | 8.779 | ***0.000 |
| | SVM (poly)<br>LDA | 55<br>48 | 11.951 | ***0.000 |
| | SVM (poly)<br>KNN | 55<br>44 | 9.289 | ***0.000 |
| | SVM (RBF)<br>Random Forest | 58<br>43 | 18.370 | ***0.000 |
| | SVM (RBF)<br>LDA | 58<br>48 | 13.224 | ***0.000 |
| | SVM (RBF)<br>KNN | 58<br>45 | 10.110 | ***0.000 |
| | Random Forest<br>LDA | 43<br>48 | -3.074 | *0.028 |
| | Random Forest<br>KNN | 43<br>44 | -0.678 | 0.528 |
| | LDA<br>KNN | 48<br>44 | 3.454 | *0.018 |
| | Dummy<br>SVM (poly) | 20<br>60 | -32.654 | ***0.000 |
| | Dummy<br>SVM (RBF) | 21<br>57 | -23.619 | ***0.000 |
| | Dummy<br>Random Forest | 21<br>43 | -11.246 | ***0.000 |
| | Dummy<br>LDA | 21<br>48 | -20.455 | ***0.000 |
| | Dummy<br>KNN | 20<br>44 | -17.767 | ***0.000 |
| Subject F | SVM (poly)<br>SVM (RBF) | 37<br>40 | -0.351 | 0.740 |
| | SVM (poly)<br>Random Forest | 38<br>37 | 0.699 | 0.516 |
| | SVM (poly)<br>LDA | 40<br>27 | 7.083 | *0.001 |
| | SVM (poly)<br>KNN | 38<br>36 | 2.404 | 0.061 |
| | SVM (RBF)<br>Random Forest | 42<br>37 | 1.069 | 0.334 |
| | SVM (RBF)<br>LDA | 41<br>26 | 5.002 | **0.004 |
| | SVM (RBF)<br>KNN | 41<br>36 | 1.945 | 0.109 |
| | Random Forest<br>LDA | 36<br>25 | 2.997 | *0.030 |
| | Random Forest<br>KNN | 36<br>34 | 0.746 | 0.489 |
| | LDA<br>KNN | 25<br>33 | -2.839 | *0.036 |
| | Dummy<br>SVM (poly) | 20<br>38 | -7.326 | **0.001 |
| | Dummy<br>SVM (RBF) | 20<br>41 | -5.688 | **0.002 |
| | Dummy<br>Random Forest | 19<br>36 | -4.328 | **0.008 |
| | Dummy<br>LDA | 20<br>26 | -0.717 | 0.506 |
| | Dummy<br>KNN | 20<br>34 | -3.500 | *0.017 |
| Subject G | SVM (poly)<br>SVM (RBF) | 37<br>38 | -2.315 | 0.068 |
| | SVM (poly) | 37 | 4.440 | **0.007 |

| | | | | |
|---|---|---|---|---|
| | Random Forest | 34 | | |
| | SVM (poly)<br>LDA | 37<br>30 | 6.292 | **0.001 |
| | SVM (poly)<br>KNN | 37<br>30 | 2.447 | 0.058 |
| | SVM (RBF)<br>Random Forest | 38<br>34 | 3.845 | *0.012 |
| | SVM (RBF)<br>LDA | 38<br>28 | 6.502 | **0.001 |
| | SVM (RBF)<br>KNN | 38<br>30 | 5.292 | **0.003 |
| | Random Forest<br>LDA | 35<br>28 | 5.280 | **0.003 |
| | Random Forest<br>KNN | 34<br>30 | 0.335 | 0.751 |
| | LDA<br>KNN | 29<br>30 | -1.450 | 0.207 |
| | Dummy<br>SVM (poly) | 20<br>37 | -8.880 | ***0.000 |
| | Dummy<br>SVM (RBF) | 20<br>38 | -11.955 | ***0.000 |
| | Dummy<br>Random Forest | 21<br>35 | -5.407 | **0.003 |
| | Dummy<br>LDA | 21<br>29 | -3.266 | *0.022 |
| | Dummy<br>KNN | 21<br>30 | -8.475 | ***0.000 |
| Subject H | SVM (poly)<br>SVM (RBF) | 24<br>21 | 0.000 | 1.000 |
| | SVM (poly)<br>Random Forest | 24<br>22 | -0865 | 0.427 |
| | SVM (poly)<br>LDA | 24<br>21 | 1.040 | 0.346 |
| | SVM (poly)<br>KNN | 24<br>21 | 0.747 | 0.489 |
| | SVM (RBF)<br>Random Forest | 21<br>22 | -0.365 | 0.730 |
| | SVM (RBF)<br>LDA | 21<br>21 | 0.914 | 0.403 |
| | SVM (RBF)<br>KNN | 21<br>20 | 0.532 | 0.617 |
| | Random Forest<br>LDA | 22<br>20 | 1.192 | 0.287 |
| | Random Forest<br>KNN | 22<br>20 | 1.285 | 0.255 |
| | LDA<br>KNN | 21<br>20 | -0.693 | 0.519 |
| | Dummy<br>SVM (poly) | 20<br>23 | -2.597 | *0.048 |
| | Dummy<br>SVM (RBF) | 20<br>21 | -2.711 | *0.042 |
| | Dummy<br>Random Forest | 20<br>22 | -2.218 | 0.077 |
| | Dummy<br>LDA | 19<br>20 | -1.562 | 0.179 |
| | Dummy<br>KNN | 19<br>20 | -1.638 | 0.162 |

| | | | | |
|---|---|---|---|---|
| Subject I | SVM (poly)<br>SVM (RBF) | 36<br>40 | -1.116 | 0.315 |
| | SVM (poly)<br>Random Forest | 36<br>35 | 0.707 | 0.511 |
| | SVM (poly)<br>LDA | 36<br>30 | 3.793 | *0.013 |

| | | | | |
|---|---|---|---|---|
| | SVM (poly)<br>KNN | 37<br>34 | -0.120 | 0.909 |
| | SVM (RBF)<br>Random Forest | 39<br>35 | 3.316 | *0.021 |
| | SVM (RBF)<br>LDA | 40<br>30 | 3.561 | *0.016 |
| | SVM (RBF)<br>KNN | 40<br>35 | 2.087 | 0.091 |
| | Random Forest<br>LDA | 36<br>30 | 2.721 | *0.042 |
| | Random Forest<br>KNN | 36<br>34 | -1.633 | 0.163 |
| | LDA<br>KNN | 29<br>34 | -3.646 | *0.015 |
| | Dummy<br>SVM (poly) | 21<br>35 | -8.768 | ***0.000 |
| | Dummy<br>SVM (RBF) | 20<br>38 | -9.188 | ***0.000 |
| | Dummy<br>Random Forest | 21<br>36 | -11.123 | ***0.000 |
| | Dummy<br>LDA | 20<br>30 | -7.553 | **0.001 |
| | Dummy<br>KNN | 20<br>34 | -10.626 | ***0.000 |
| Independent | SVM (poly)<br>SVM (RBF) | 35<br>37 | -2.508 | 0.054 |
| | SVM (poly)<br>Random Forest | 34<br>23 | 2.010 | 0.101 |
| | SVM (poly)<br>LDA | 34<br>35 | -1.321 | 0.244 |
| | SVM (poly)<br>KNN | 34<br>30 | 4.098 | 0.009** |
| | SVM (RBF)<br>Random Forest | 37<br>32 | 6.295 | 0.001** |
| | SVM (RBF)<br>LDA | 37<br>35 | 0.908 | 0.405 |
| | SVM (RBF)<br>KNN | 37<br>30 | 5.999 | 0.002** |
| | Random Forest<br>LDA | 32<br>36 | -3.630 | 0.015* |
| | Random Forest<br>KNN | 32<br>30 | 3.956 | 0.011* |
| | LDA<br>KNN | 36<br>30 | 6.421 | 0.001** |
| | Dummy<br>SVM (poly) | 21<br>34 | -9.029 | 0.000*** |
| | Dummy<br>SVM (RBF) | 21<br>37 | -12.692 | 0.000*** |
| | Dummy<br>Random Forest | 21<br>32 | -14.834 | 0.000*** |
| | Dummy<br>LDA | 21<br>36 | -13.381 | 0.000*** |
| | Dummy<br>KNN | 21<br>30 | -12.021 | 0.000*** |

**Table 2S**, “5x2 Cross-Validation Test” in Time Domain, After PCA scenario.
SVM (kernel='poly', degree=3, C=100000),
SVM(kernel='rbf', gamma=0.001, C=100),
RandomForestClassifier(max_depth=5, random_state=20),
KNeighborsClassifier(n_neighbors = 60).

| Subjects | Classifiers | Accuracies % | t Statistics | P-Value |
|---|---|---|---|---|
| Subject B | SVM (poly)<br>SVM (RBF) | 37<br>44 | -4.458 | **0.007 |
| | SVM (poly)<br>Random Forest | 37<br>34 | -0.740 | 0.493 |
| | SVM (poly)<br>LDA | 37<br>44 | -2.626 | *0.047 |
| | SVM (poly)<br>KNN | 37<br>34 | 1.563 | 0.179 |
| | SVM (RBF)<br>Random Forest | 44<br>36 | 3.863 | *0.012 |
| | SVM (RBF)<br>LDA | 44<br>43 | 1.279 | 0.257 |
| | SVM (RBF)<br>KNN | 44<br>34 | 6.732 | **0.001 |
| | Random Forest<br>LDA | 36<br>44 | -2.975 | *0.031 |
| | Random Forest<br>KNN | 36<br>34 | 0.877 | 0.421 |
| | LDA<br>KNN | 44<br>34 | 3.904 | *0.011 |
| | Dummy<br>SVM (poly) | 20<br>38 | -6.595 | **0.001 |
| | Dummy<br>SVM (RBF) | 19<br>44 | -13.021 | ***0.000 |
| | Dummy<br>Random Forest | 20<br>35 | -4.637 | **0.006 |
| | Dummy<br>LDA | 20<br>44 | -11.270 | ***0.000 |
| | Dummy<br>KNN | 20<br>34 | -5.533 | **0.003 |
| Subject C | SVM (poly)<br>SVM (RBF) | 47<br>54 | -1.602 | 0.170 |
| | SVM (poly)<br>Random Forest | 48<br>44 | 1.276 | 0.258 |
| | SVM (poly)<br>LDA | 48<br>53 | -2.015 | 0.100 |
| | SVM (poly)<br>KNN | 47<br>42 | 1.994 | 0.103 |
| | SVM (RBF)<br>Random Forest | 54<br>44 | 6.460 | **0.001 |
| | SVM (RBF)<br>LDA | 54<br>55 | -0.953 | 0.384 |
| | SVM (RBF)<br>KNN | 54<br>41 | 2.780 | *0.039 |
| | Random Forest<br>LDA | 44<br>56 | -4.010 | *0.010 |
| | Random Forest<br>KNN | 44<br>41 | 1.218 | 0.278 |
| | LDA<br>KNN | 54<br>41 | 3.685 | *0.014 |
| | Dummy<br>SVM (poly) | 20<br>47 | -22.512 | ***0.000 |
| | Dummy | 20 | -12.259 | ***0.000 |

| | | | | |
|---|---|---|---|---|
| | SVM (RBF) | 54 | | |
| | Dummy<br>Random Forest | 19<br>44 | -11.844 | ***0.000 |
| | Dummy<br>LDA | 20<br>56 | -13.074 | ***0.000 |
| | Dummy<br>KNN | 19<br>41 | -8.413 | ***0.000 |
| Subject E | SVM (poly)<br>SVM (RBF) | 51<br>57 | -5.778 | **0.002 |
| | SVM (poly)<br>Random Forest | 51<br>46 | 6.241 | **0.002 |
| | SVM (poly)<br>LDA | 51<br>58 | -7.213 | **0.001 |
| | SVM (poly)<br>KNN | 51<br>45 | 10.523 | ***0.000 |
| | SVM (RBF)<br>Random Forest | 57<br>46 | 15.775 | ***0.000 |
| | SVM (RBF)<br>LDA | 57<br>58 | -1.211 | 0.280 |
| | SVM (RBF)<br>KNN | 57<br>45 | 12.745 | ***0.000 |
| | Random Forest<br>LDA | 46<br>58 | -10.435 | ***0.000 |
| | Random Forest<br>KNN | 46<br>44 | -0.586 | 0.583 |
| | LDA<br>KNN | 58<br>45 | 8.365 | ***0.000 |
| | Dummy<br>SVM (poly) | 21<br>52 | -29.161 | ***0.000 |
| | Dummy<br>SVM (RBF) | 22<br>57 | -24.462 | ***0.000 |
| | Dummy<br>Random Forest | 21<br>46 | -9.800 | ***0.000 |
| | Dummy<br>LDA | 21<br>58 | -21.442 | ***0.000 |
| | Dummy<br>KNN | 22<br>45 | -23.283 | ***0.000 |
| Subject F | SVM (poly)<br>SVM (RBF) | 35<br>41 | -3.139 | *0.026 |
| | SVM (poly)<br>Random Forest | 34<br>34 | 1.313 | 0.246 |
| | SVM (poly)<br>LDA | 33<br>38 | -7.779 | **0.001 |
| | SVM (poly)<br>KNN | 33<br>34 | -0.568 | 0.594 |
| | SVM (RBF)<br>Random Forest | 40<br>34 | 4.672 | **0.005 |
| | SVM (RBF)<br>LDA | 40<br>38 | -0.104 | 0.921 |
| | SVM (RBF)<br>KNN | 41<br>35 | 1.898 | 0.116 |
| | Random Forest<br>LDA | 32<br>39 | -4.744 | **0.005 |
| | Random Forest<br>KNN | 34<br>35 | -0.726 | 0.500 |
| | LDA<br>KNN | 40<br>35 | 2.289 | 0.071 |
| | Dummy<br>SVM (poly) | 20<br>35 | -5.005 | **0.004 |
| | Dummy<br>SVM (RBF) | 20<br>41 | -5.800 | **0.002 |
| | Dummy<br>Random Forest | 19<br>33 | -2.693 | *0.043 |
| | Dummy<br>LDA | 20<br>40 | -16.449 | ***0.000 |
| | Dummy | 19 | -3.817 | *0.012 |

| | | | | |
|---|---|---|---|---|
| | KNN | 33 | | |
| Subject G | SVM (poly)<br>SVM (RBF) | 34<br>38 | -1.801 | 0.132 |
| | SVM (poly)<br>Random Forest | 34<br>32 | 0.643 | 0.549 |
| | SVM (poly)<br>LDA | 34<br>39 | -3.024 | *0.029 |
| | SVM (poly)<br>KNN | 35<br>30 | 0.698 | 0.516 |
| | SVM (RBF)<br>Random Forest | 39<br>33 | 2.017 | 0.100 |
| | SVM (RBF)<br>LDA | 38<br>39 | -0.120 | 0.909 |
| | SVM (RBF)<br>KNN | 39<br>30 | 2.669 | *0.044 |
| | Random Forest<br>LDA | 33<br>38 | -3.805 | *0.013 |
| | Random Forest<br>KNN | 33<br>30 | -0.036 | 0.973 |
| | LDA<br>KNN | 39<br>29 | 3.214 | *0.024 |
| | Dummy<br>SVM (poly) | 21<br>34 | -10.095 | ***0.000 |
| | Dummy<br>SVM (RBF) | 21<br>38 | -10.925 | ***0.000 |
| | Dummy<br>Random Forest | 21<br>33 | -6.361 | **0.001 |
| | Dummy<br>LDA | 21<br>38 | -9.102 | ***0.000 |
| | Dummy<br>KNN | 21<br>30 | -6.156 | **0.002 |
| Subject H | SVM (poly)<br>SVM (RBF) | 22<br>22 | 0.409 | 0.699 |
| | SVM (poly)<br>Random Forest | 22<br>21 | 0.802 | 0.459 |
| | SVM (poly)<br>LDA | 23<br>25 | 0.952 | 0.385 |
| | SVM (poly)<br>KNN | 22<br>22 | 1.337 | 0.239 |
| | SVM (RBF)<br>Random Forest | 22<br>23 | 1.609 | 0.168 |
| | SVM (RBF)<br>LDA | 21<br>23 | 0.783 | 0.469 |
| | SVM (RBF)<br>KNN | 22<br>21 | 0.921 | 0.399 |
| | Random Forest<br>LDA | 24<br>25 | -0.117 | 0.912 |
| | Random Forest<br>KNN | 23<br>21 | 0.399 | 0.706 |
| | LDA<br>KNN | 25<br>20 | -0.099 | 0.925 |
| | Dummy<br>SVM (poly) | 21<br>22 | -1.954 | 0.108 |
| | Dummy<br>SVM (RBF) | 20<br>21 | -2.681 | *0.044 |
| | Dummy<br>Random Forest | 20<br>22 | -0.487 | 0.647 |
| | Dummy<br>LDA | 20<br>24 | -1.701 | 0.150 |
| | Dummy<br>KNN | 20<br>19 | -1.362 | 0.231 |
| Subject I | SVM (poly)<br>SVM (RBF) | 37<br>40 | -0.169 | 0.872 |
| | SVM (poly)<br>Random Forest | 37<br>37 | 0.755 | 0.485 |
| | SVM (poly) | 38 | -0.634 | 0.554 |

| | | | | |
|---|---|---|---|---|
| | LDA | 41 | | |
| | SVM (poly)<br>KNN | 38<br>35 | 1.086 | 0.327 |
| | SVM (RBF)<br>Random Forest | 39<br>37 | 0.785 | 0.468 |
| | SVM (RBF)<br>LDA | 40<br>41 | -0.190 | 0.857 |
| | SVM (RBF)<br>KNN | 39<br>35 | 2.114 | 0.088 |
| | Random Forest<br>LDA | 37<br>40 | -1.872 | 0.120 |
| | Random Forest<br>KNN | 36<br>35 | 3.080 | *0.027 |
| | LDA<br>KNN | 41<br>35 | 5.216 | **0.003 |
| | Dummy<br>SVM (poly) | 21<br>37 | -6.486 | **0.001 |
| | Dummy<br>SVM (RBF) | 21<br>39 | -10.119 | ***0.000 |
| | Dummy<br>Random Forest | 21<br>36 | -10.996 | ***0.000 |
| | Dummy<br>LDA | 21<br>41 | -12.982 | ***0.000 |
| | Dummy<br>KNN | 21<br>34 | -8.116 | ***0.000 |
| Independent | SVM (poly)<br>SVM (RBF) | 33<br>37 | -2.687 | 0.043* |
| | SVM (poly)<br>Random Forest | 33<br>34 | -0.644 | 0.548 |
| | SVM (poly)<br>LDA | 33<br>40 | -10.732 | 0.000*** |
| | SVM (poly)<br>KNN | 34<br>31 | 6.006 | 0.002** |
| | SVM (RBF)<br>Random Forest | 37<br>34 | 0.892 | 0.413 |
| | SVM (RBF)<br>LDA | 37<br>40 | -3.415 | 0.019* |
| | SVM (RBF)<br>KNN | 37<br>30 | 11.514 | 0.000*** |
| | Random Forest<br>LDA | 34<br>40 | -2.238 | 0.075 |
| | Random Forest<br>KNN | 34<br>30 | 2.993 | 0.030* |
| | LDA<br>KNN | 40<br>30 | 7.743 | 0.001** |
| | Dummy<br>SVM (poly) | 21<br>34 | -15.892 | 0.000*** |
| | Dummy<br>SVM (RBF) | 21<br>37 | -13.714 | 0.000*** |
| | Dummy<br>Random Forest | 21<br>34 | -6.423 | 0.001** |
| | Dummy<br>LDA | 21<br>40 | -12.661 | 0.000*** |
| | Dummy<br>KNN | 21<br>30 | -10.790 | 0.000*** |

**Table 3S,** "5x2 Cross-Validation Test" in Time-frequency Domain (Morlet), Before PCA scenario,
SVM (kernel='poly', degree = 3, C = 100000),
SVM (kernel='rbf', gamma=0.001, C=100)
RandomForestClassifier (max_depth=5, random_state=20),
KNeighborsClassifier (n_neighbors = 60).

| Subjects | Classifiers | Accuracies % | t Statistics | P-Value |
|---|---|---|---|---|
| Subject B | SVM (poly)<br>SVM (RBF) | 24<br>25 | -0.577 | 0.589 |
| | SVM (poly)<br>Random Forest | 25<br>26 | -0.304 | 0.774 |
| | SVM (poly)<br>LDA | 24<br>22 | 4.385 | **0.007 |
| | SVM (poly)<br>KNN | 24<br>23 | 2.010 | 0.101 |
| | SVM (RBF)<br>Random Forest | 25<br>26 | 0.247 | 0.815 |
| | SVM (RBF)<br>LDA | 25<br>22 | 3.165 | *0.025 |
| | SVM (RBF)<br>KNN | 25<br>24 | 1.462 | 0.204 |
| | Random Forest<br>LDA | 26<br>22 | 3.033 | *0.029 |
| | Random Forest<br>KNN | 25<br>23 | 1.086 | 0.327 |
| | LDA<br>KNN | 22<br>23 | -2.353 | 0.065 |
| | Dummy<br>SVM (poly) | 21<br>25 | -6.044 | **0.002 |
| | Dummy<br>SVM (RBF) | 21<br>25 | -3.972 | *0.011 |
| | Dummy<br>Random Forest | 20<br>26 | -2.516 | 0.053 |
| | Dummy<br>LDA | 19<br>22 | 0.484 | 0.649 |
| | Dummy<br>KNN | 21<br>24 | -2.637 | *0.046 |
| Subject C | SVM (poly)<br>SVM (RBF) | 31<br>32 | -0.984 | 0.370 |
| | SVM (poly)<br>Random Forest | 31<br>32 | -3.487 | *0.018 |
| | SVM (poly)<br>LDA | 30<br>28 | 2.335 | 0.067 |
| | SVM (poly)<br>KNN | 31<br>29 | -1.023 | 0.353 |
| | SVM (RBF)<br>Random Forest | 31<br>32 | -0.715 | 0.507 |
| | SVM (RBF)<br>LDA | 33<br>27 | 3.370 | *0.020 |
| | SVM (RBF)<br>KNN | 32<br>29 | -0.155 | 0.883 |
| | Random Forest<br>LDA | 31<br>27 | 3.600 | *0.016 |
| | Random Forest<br>KNN | 31<br>28 | 0.318 | 0.763 |
| | LDA<br>KNN | 27<br>29 | -1.869 | 0.121 |
| | Dummy<br>SVM (poly) | 21<br>31 | -4.554 | **0.006 |
| | Dummy<br>SVM (RBF) | 20<br>33 | -6.600 | **0.001 |
| | Dummy | 20 | -6.601 | **0.001 |

| | | | | |
|---|---|---|---|---|
| | Random Forest | 31 | | |
| | Dummy<br>LDA | 21<br>27 | -3.864 | *0.012 |
| | Dummy<br>KNN | 21<br>30 | -3.451 | *0.018 |
| Subject E | SVM (poly)<br>SVM (RBF) | 37<br>39 | -1.842 | 0.125 |
| | SVM (poly)<br>Random Forest | 36<br>37 | -0.663 | 0.537 |
| | SVM (poly)<br>LDA | 36<br>27 | 12.021 | ***0.000 |
| | SVM (poly)<br>KNN | 37<br>33 | 3.441 | *0.018 |
| | SVM (RBF)<br>Random Forest | 38<br>37 | 0.858 | 0.430 |
| | SVM (RBF)<br>LDA | 38<br>37 | 24.171 | ***0.000 |
| | SVM (RBF)<br>KNN | 38<br>34 | 4.206 | **0.008 |
| | Random Forest<br>LDA | 37<br>27 | 6.889 | **0.001 |
| | Random Forest<br>KNN | 37<br>34 | 3.495 | *0.017 |
| | LDA<br>KNN | 27<br>33 | -3.358 | *0.020 |
| | Dummy<br>SVM (poly) | 21<br>37 | -9.308 | ***0.000 |
| | Dummy<br>SVM (RBF) | 21<br>39 | -15.160 | ***0.000 |
| | Dummy<br>Random Forest | 21<br>36 | -8.288 | ***0.000 |
| | Dummy<br>LDA | 22<br>27 | -4.440 | **0.007 |
| | Dummy<br>KNN | 21<br>33 | -4.938 | **0.004 |
| Subject F | SVM (poly)<br>SVM (RBF) | 25<br>26 | -0.246 | 0.816 |
| | SVM (poly)<br>Random Forest | 26<br>24 | 1.827 | 0.127 |
| | SVM (poly)<br>LDA | 26<br>25 | 2.842 | *0.036 |
| | SVM (poly)<br>KNN | 26<br>24 | 2.423 | 0.060 |
| | SVM (RBF)<br>Random Forest | 26<br>25 | 1.991 | 0.103 |
| | SVM (RBF)<br>LDA | 27<br>25 | 3.023 | *0.029 |
| | SVM (RBF)<br>KNN | 26<br>23 | 3.448 | *0.018 |
| | Random Forest<br>LDA | 24<br>25 | 0.350 | 0.740 |
| | Random Forest<br>KNN | 25<br>23 | 0.508 | 0.633 |
| | LDA<br>KNN | 24<br>23 | 0.135 | 0.898 |
| | Dummy<br>SVM (poly) | 19<br>26 | -6.859 | **0.001 |
| | Dummy<br>SVM (RBF) | 20<br>27 | -5.870 | **0.002 |
| | Dummy<br>Random Forest | 20<br>25 | -0.591 | 0.580 |
| | Dummy<br>LDA | 20<br>24 | -0.364 | 0.731 |
| | Dummy<br>KNN | 20<br>22 | -0.201 | 0.848 |

| | | | | |
|---|---|---|---|---|
| Subject G | SVM (poly)<br>SVM (RBF) | 26<br>27 | -1.270 | 0.260 |
| | SVM (poly)<br>Random Forest | 25<br>27 | -0.050 | 0.962 |
| | SVM (poly)<br>LDA | 25<br>22 | 2.755 | *0.040 |
| | SVM (poly)<br>KNN | 26<br>25 | 0.641 | 0.550 |
| | SVM (RBF)<br>Random Forest | 27<br>27 | 1.711 | 0.148 |
| | SVM (RBF)<br>LDA | 26<br>23 | 6.748 | **0.001 |
| | SVM (RBF)<br>KNN | 27<br>24 | 1.415 | 0.216 |
| | Random Forest<br>LDA | 27<br>22 | 2.369 | 0.064 |
| | Random Forest<br>KNN | 26<br>24 | 0.644 | 0.548 |
| | LDA<br>KNN | 23<br>25 | -0.791 | 0.465 |
| | Dummy<br>SVM (poly) | 22<br>25 | -3.473 | *0.018 |
| | Dummy<br>SVM (RBF) | 22<br>27 | -4.892 | **0.005 |
| | Dummy<br>Random Forest | 21<br>26 | -3.072 | *0.028 |
| | Dummy<br>LDA | 21<br>22 | -1.574 | 0.176 |
| | Dummy<br>KNN | 21<br>25 | -2.457 | 0.057 |
| Subject H | SVM (poly)<br>SVM (RBF) | 21<br>23 | -2.099 | 0.090 |
| | SVM (poly)<br>Random Forest | 21<br>22 | -2.204 | 0.079 |
| | SVM (poly)<br>LDA | 22<br>21 | -0.887 | 0.416 |
| | SVM (poly)<br>KNN | 22<br>22 | -1.106 | 0.319 |
| | SVM (RBF)<br>Random Forest | 22<br>22 | -0.625 | 0.559 |
| | SVM (RBF)<br>LDA | 22<br>21 | 1.416 | 0.216 |
| | SVM (RBF)<br>KNN | 22<br>21 | 0.256 | 0.808 |
| | Random Forest<br>LDA | 22<br>21 | 1.643 | 0.161 |
| | Random Forest<br>KNN | 22<br>21 | 0.875 | 0.422 |
| | LDA<br>KNN | 20<br>21 | -0.606 | 0.571 |
| | Dummy<br>SVM (poly) | 21<br>22 | -0.927 | 0.396 |
| | Dummy<br>SVM (RBF) | 21<br>23 | -2.139 | 0.085 |
| | Dummy<br>Random Forest | 21<br>23 | -1.906 | 0.115 |
| | Dummy<br>LDA | 20<br>21 | -1.540 | 0.184 |
| | Dummy<br>KNN | 19<br>22 | -1.395 | 0.222 |
| Subject I | SVM (poly)<br>SVM (RBF) | 28<br>28 | 0.427 | 0.687 |
| | SVM (poly)<br>Random Forest | 28<br>31 | -2.741 | *0.041 |
| | SVM (poly)<br>LDA | 27<br>23 | 1.059 | 0.338 |

| | | | | |
|---|---|---|---|---|
| | SVM (poly)<br>KNN | 27<br>27 | 0.769 | 0.477 |
| | SVM (RBF)<br>Random Forest | 27<br>31 | -3.142 | *0.026 |
| | SVM (RBF)<br>LDA | 28<br>24 | 0.857 | 0.431 |
| | SVM (RBF)<br>KNN | 27<br>26 | 0.303 | 0.774 |
| | Random Forest<br>LDA | 30<br>23 | 4.008 | *0.010 |
| | Random Forest<br>KNN | 31<br>27 | 2.189 | 0.080 |
| | LDA<br>KNN | 24<br>26 | -0.473 | 0.656 |
| | Dummy<br>SVM (poly) | 21<br>28 | -3.588 | *0.016 |
| | Dummy<br>SVM (RBF) | 20<br>27 | -4.290 | **0.008 |
| | Dummy<br>Random Forest | 21<br>31 | -7.468 | **0.001 |
| | Dummy<br>LDA | 20<br>23 | -1.826 | 0.127 |
| | Dummy<br>KNN | 21<br>26 | -2.292 | 0.070 |
| Independent | SVM (poly)<br>SVM (RBF) | 25<br>28 | -4.513 | 0.006** |
| | SVM (poly)<br>Random Forest | 25<br>28 | -4.927 | 0.004** |
| | SVM (poly)<br>LDA | 25<br>21 | 6.689 | 0.001** |
| | SVM (poly)<br>KNN | 25<br>25 | -1.929 | 0.112 |
| | SVM (RBF)<br>Random Forest | 28<br>28 | -0.616 | 0.565 |
| | SVM (RBF)<br>LDA | 28<br>22 | 6.300 | 0.001** |
| | SVM (RBF)<br>KNN | 28<br>25 | 3.551 | 0.016* |
| | Random Forest<br>LDA | 28<br>21 | 6.537 | 0.001** |
| | Random Forest<br>KNN | 28<br>25 | 3.800 | 0.013* |
| | LDA<br>KNN | 21<br>25 | -4.118 | 0.009** |
| | Dummy<br>SVM (poly) | 21<br>24 | -1.934 | 0.111 |
| | Dummy<br>SVM (RBF) | 21<br>29 | -4.969 | 0.004** |
| | Dummy<br>Random Forest | 21<br>28 | -6.517 | 0.001** |
| | Dummy<br>LDA | 21<br>21 | 0.319 | 0.762 |
| | Dummy<br>KNN | 21<br>25 | -4.020 | 0.010* |

**Table 4S,** “5x2 Cross-Validation Test” in Time-frequency Domain (Morlet), After PCA scenario,
SVM (kernel='poly', degree = 3, C = 100000),
SVM (kernel='rbf', gamma=0.001, C=100),
RandomForestClassifier (max_depth=5, random_state=20),
KNeighborsClassifier (n_neighbors = 60)

| **Subjects** | **Classifiers** | **Accuracies %** | **t Statistics** | **P-Value** |
|---|---|---|---|---|
| Subject B | SVM (poly)<br>SVM (RBF) | 23<br>24 | -0.911 | 0.404 |
| | SVM (poly)<br>Random Forest | 24<br>23 | 0.917 | 0.504 |
| | SVM (poly)<br>LDA | 24<br>25 | -2.262 | 0.073 |
| | SVM (poly)<br>KNN | 24<br>24 | 0.698 | 0.517 |
| | SVM (RBF)<br>Random Forest | 24<br>23 | 2.828 | *0.037 |
| | SVM (RBF)<br>LDA | 24<br>26 | -0.714 | 0.507 |
| | SVM (RBF)<br>KNN | 24<br>23 | 1.025 | 0.352 |
| | Random Forest<br>LDA | 23<br>26 | -3.069 | *0.028 |
| | Random Forest<br>KNN | 23<br>23 | -0.541 | 0.612 |
| | LDA<br>KNN | 26<br>24 | 1.434 | 0.211 |
| | Dummy<br>SVM (poly) | 21<br>24 | -2.381 | 0.063 |
| | Dummy<br>SVM (RBF) | 21<br>25 | -2.662 | *0.045 |
| | Dummy<br>Random Forest | 21<br>24 | 0.328 | 0.756 |
| | Dummy<br>LDA | 21<br>24 | -3.499 | *0.017 |
| | Dummy<br>KNN | 21<br>24 | -1.480 | 0.199 |
| Subject C | SVM (poly)<br>SVM (RBF) | 30<br>32 | -1.832 | 0.126 |
| | SVM (poly)<br>Random Forest | 29<br>27 | -0.214 | 0.819 |
| | SVM (poly)<br>LDA | 28<br>32 | -2.040 | 0.097 |
| | SVM (poly)<br>KNN | 28<br>29 | -1.328 | 0.241 |
| | SVM (RBF)<br>Random Forest | 32<br>29 | -1.690 | 0.152 |
| | SVM (RBF)<br>LDA | 31<br>34 | -1.221 | 0.277 |
| | SVM (RBF)<br>KNN | 32<br>28 | -0.083 | 0.937 |
| | Random Forest<br>LDA | 29<br>32 | -1.971 | 0.106 |
| | Random Forest<br>KNN | 27<br>28 | -0.370 | 0.727 |
| | LDA<br>KNN | 31<br>29 | 1.400 | 0.220 |
| | Dummy<br>SVM (poly) | 20<br>29 | -7.023 | **0.001 |
| | Dummy<br>SVM (RBF) | 21<br>31 | -4.864 | **0.005 |
| | Dummy<br>Random Forest | 20<br>28 | -5.565 | **0.003 |

| | | | | |
|---|---|---|---|---|
| | Dummy<br>LDA | 20<br>32 | -6.733 | **0.001 |
| | Dummy<br>KNN | 20<br>29 | -3.853 | *0.012 |
| Subject E | SVM (poly)<br>SVM (RBF) | 35<br>38 | -3.188 | *0.024 |
| | SVM (poly)<br>Random Forest | 35<br>34 | 1.773 | 0.136 |
| | SVM (poly)<br>LDA | 35<br>39 | -3.829 | 0.012 |
| | SVM (poly)<br>KNN | 35<br>34 | 1.470 | 0.202 |
| | SVM (RBF)<br>Random Forest | 39<br>34 | 2.483 | 0.056 |
| | SVM (RBF)<br>LDA | 39.2<br>39.3 | -2.728 | *0.041 |
| | SVM (RBF)<br>KNN | 39<br>34 | 4.090 | **0.009 |
| | Random Forest<br>LDA | 35<br>39 | -3.444 | *0.018 |
| | Random Forest<br>KNN | 35<br>34 | 0.445 | 0.675 |
| | LDA<br>KNN | 40<br>34 | 4.595 | **0.006 |
| | Dummy<br>SVM (poly) | 21<br>35 | -10.495 | ***0.000 |
| | Dummy<br>SVM (RBF) | 21<br>39 | -10.350 | ***0.000 |
| | Dummy<br>Random Forest | 21<br>35 | -6.190 | **0.002 |
| | Dummy<br>LDA | 21<br>39 | -10.514 | ***0.000 |
| | Dummy<br>KNN | 22<br>34 | -3.991 | *0.010 |
| Subject F | SVM (poly)<br>SVM (RBF) | 25<br>25 | -0.666 | 0.535 |
| | SVM (poly)<br>Random Forest | 25<br>25 | 1.578 | 0.175 |
| | SVM (poly)<br>LDA | 25<br>25 | -0.083 | 0.937 |
| | SVM (poly)<br>KNN | 25<br>23 | 3.133 | *0.026 |
| | SVM (RBF)<br>Random Forest | 26<br>24 | 2.724 | *0.042 |
| | SVM (RBF)<br>LDA | 25<br>25 | 0.688 | 0.522 |
| | SVM (RBF)<br>KNN | 27<br>22 | 1.974 | 0.105 |
| | Random Forest<br>LDA | 24<br>25 | -2.718 | *0.042 |
| | Random Forest<br>KNN | 25<br>24 | 0.504 | 0.635 |
| | LDA<br>KNN | 24<br>23 | 1.237 | 0.259 |
| | Dummy<br>SVM (poly) | 20<br>25 | -4.642 | **0.006 |
| | Dummy<br>SVM (RBF) | 20<br>25 | -2.935 | *0.032 |
| | Dummy<br>Random Forest | 19<br>23 | -2.540 | 0.052 |
| | Dummy<br>LDA | 20<br>24 | -3.891 | *0.012 |
| | Dummy<br>KNN | 21<br>22 | 0.328 | 0.756 |
| Subject G | SVM (poly)<br>SVM (RBF) | 24<br>27 | -2.449 | 0.058 |

| | | | | |
|---|---|---|---|---|
| | SVM (poly)<br>Random Forest | 24<br>24 | 1.357 | 0.233 |
| | SVM (poly)<br>LDA | 24<br>28 | -1.399 | 0.221 |
| | SVM (poly)<br>KNN | 24<br>25 | 0.684 | 0.524 |
| | SVM (RBF)<br>Random Forest | 27<br>25 | 4.062 | *0.010 |
| | SVM (RBF)<br>LDA | 27<br>28 | -0.904 | 0.408 |
| | SVM (RBF)<br>KNN | 27<br>25 | 0.591 | 0.580 |
| | Random Forest<br>LDA | 26<br>27 | -2.376 | 0.063 |
| | Random Forest<br>KNN | 25<br>25 | -0.247 | 0.814 |
| | LDA<br>KNN | 27<br>24 | 1.908 | 0.115 |
| | Dummy<br>SVM (poly) | 21<br>25 | -5.316 | **0.003 |
| | Dummy<br>SVM (RBF) | 21<br>27 | -5.313 | **0.003 |
| | Dummy<br>Random Forest | 22<br>26 | -2.269 | 0.072 |
| | Dummy<br>LDA | 20<br>28 | -4.048 | *0.010 |
| | Dummy<br>KNN | 21<br>24 | -3.787 | *0.013 |
| Subject H | SVM (poly)<br>SVM (RBF) | 20<br>22 | -1.477 | 0.200 |
| | SVM (poly)<br>Random Forest | 21<br>21 | 0.908 | 0.405 |
| | SVM (poly)<br>LDA | 21<br>23 | 0.329 | 0.755 |
| | SVM (poly)<br>KNN | 21<br>21 | -1.815 | 0.129 |
| | SVM (RBF)<br>Random Forest | 23<br>22 | 1.674 | 0.155 |
| | SVM (RBF)<br>LDA | 22<br>23 | 1.747 | 0.141 |
| | SVM (RBF)<br>KNN | 23<br>22 | -0.654 | 0.542 |
| | Random Forest<br>LDA | 20<br>23 | -0.914 | 0.403 |
| | Random Forest<br>KNN | 21<br>22 | -1.892 | 0.117 |
| | LDA<br>KNN | 22<br>22 | -0.640 | 0.550 |
| | Dummy<br>SVM (poly) | 19<br>22 | -0.752 | 0.486 |
| | Dummy<br>SVM (RBF) | 20<br>23 | -2.417 | 0.060 |
| | Dummy<br>Random Forest | 20<br>21 | 0.180 | 0.864 |
| | Dummy<br>LDA | 20<br>21 | -0.243 | 0.817 |
| | Dummy<br>KNN | 20<br>22 | -1.378 | 0.227 |
| Subject I | SVM (poly)<br>SVM (RBF) | 26<br>28 | -1.708 | 0.148 |
| | SVM (poly)<br>Random Forest | 27<br>26 | -0.762 | 0.481 |
| | SVM (poly)<br>LDA | 27<br>28 | -1.744 | 0.142 |
| | SVM (poly)<br>KNN | 26<br>27 | -1.678 | 0.154 |

| | | | | |
|---|---|---|---|---|
| | SVM (RBF)<br>Random Forest | 28<br>25 | 0.601 | 0.574 |
| | SVM (RBF)<br>LDA | 28<br>28 | -0.093 | 0.930 |
| | SVM (RBF)<br>KNN | 28<br>26 | 0.743 | 0.491 |
| | Random Forest<br>LDA | 25<br>29 | -1.551 | 0.182 |
| | Random Forest<br>KNN | 26<br>27 | 0.305 | 0.773 |
| | LDA<br>KNN | 28<br>27 | 1.858 | 0.122 |
| | Dummy<br>SVM (poly) | 21<br>27 | -1.409 | 0.218 |
| | Dummy<br>SVM (RBF) | 20<br>29 | -3.838 | *0.012 |
| | Dummy<br>Random Forest | 21<br>25 | -2.866 | *0.035 |
| | Dummy<br>LDA | 21<br>29 | -8.907 | ***0.000 |
| | Dummy<br>KNN | 21<br>27 | -3.468 | *0.018 |
| Independent | SVM (poly)<br>SVM (RBF) | 24<br>28 | -5.699 | 0.002** |
| | SVM (poly)<br>Random Forest | 24<br>26 | -3.085 | 0.027* |
| | SVM (poly)<br>LDA | 23<br>27 | -6.520 | 0.001** |
| | SVM (poly)<br>KNN | 24<br>25 | -2.494 | 0.055 |
| | SVM (RBF)<br>Random Forest | 28<br>26 | 2.023 | 0.099 |
| | SVM (RBF)<br>LDA | 28<br>28 | 0.339 | 0.748 |
| | SVM (RBF)<br>KNN | 28<br>25 | 3.363 | 0.020* |
| | Random Forest<br>LDA | 26<br>27 | -0.465 | 0.662 |
| | Random Forest<br>KNN | 26<br>25 | 1.810 | 0.130 |
| | LDA<br>KNN | 28<br>25 | 4.123 | 0.009** |
| | Dummy<br>SVM (poly) | 21<br>23 | -0.676 | 0.529 |
| | Dummy<br>SVM (RBF) | 21<br>28 | -4.682 | 0.005** |
| | Dummy<br>Random Forest | 21<br>26 | -5.835 | 0.002** |
| | Dummy<br>LDA | 21<br>28 | -4.940 | 0.004** |
| | Dummy<br>KNN | 21<br>25 | -3.159 | 0.025* |

**Table 5S,** "5x2 Cross-Validation Test" in Time-frequency Domain (Wigner), Before PCA scenario,
SVM (kernel='poly', degree = 3, C = 100000),
SVM (kernel='rbf', gamma=0.001, C=100),
RandomForestClassifier (max_depth=5, random_state=20),
KNeighborsClassifier (n_neighbors = 60)

| Subjects | Classifiers | Accuracies % | t Statistics | P-Value |
|---|---|---|---|---|
| Subject B | SVM (poly)<br>SVM (RBF) | 32<br>34 | 0.893 | 0.413 |
| | SVM (poly)<br>Random Forest | 32<br>31 | 0.602 | 0.574 |
| | SVM (poly)<br>LDA | 32<br>30 | 2.257 | 0.074 |
| | SVM (poly)<br>KNN | 32<br>22 | 4.536 | **0.006 |
| | SVM (RBF)<br>Random Forest | 33<br>32 | 0.038 | 0.971 |
| | SVM (RBF)<br>LDA | 34<br>30 | 1.200 | 0.284 |
| | SVM (RBF)<br>KNN | 32<br>22 | 4.503 | **0.006 |
| | Random Forest<br>LDA | 32<br>29 | 1.164 | 0.297 |
| | Random Forest<br>KNN | 32<br>23 | 4.096 | **0.009 |
| | LDA<br>KNN | 31<br>22 | 3.737 | *0.013 |
| | Dummy<br>SVM (poly) | 21<br>31 | -4.886 | **0.005 |
| | Dummy<br>SVM (RBF) | 20<br>33 | -4.323 | **0.008 |
| | Dummy<br>Random Forest | 21<br>31 | -5.709 | **0.002 |
| | Dummy<br>LDA | 21<br>30 | -6.257 | **0.002 |
| | Dummy<br>KNN | 22<br>22 | 0.512 | 0.630 |
| Subject C | SVM (poly)<br>SVM (RBF) | 43<br>42 | -1.294 | 0.252 |
| | SVM (poly)<br>Random Forest | 43<br>39 | 1.856 | 0.123 |
| | SVM (poly)<br>LDA | 43<br>39 | 3.088 | *0.027 |
| | SVM (poly)<br>KNN | 43<br>25 | 9.188 | ***0.000 |
| | SVM (RBF)<br>Random Forest | 43<br>40 | 2.597 | *0.048 |
| | SVM (RBF)<br>LDA | 43<br>39 | 5.398 | **0.003 |
| | SVM (RBF)<br>KNN | 42<br>25 | 7.912 | **0.001 |
| | Random Forest<br>LDA | 43<br>39 | 5.398 | **0.003 |
| | Random Forest<br>KNN | 39<br>26 | 3.415 | *0.019 |
| | LDA<br>KNN | 39<br>25 | 4.490 | **0.006 |
| | Dummy<br>SVM (poly) | 19<br>43 | -9.426 | ***0.000 |
| | Dummy<br>SVM (RBF) | 19<br>42 | -8.637 | ***0.000 |
| | Dummy<br>Random Forest | 19<br>37 | -8.590 | ***0.000 |
| | Dummy<br>LDA | 19<br>38 | -7.786 | **0.001 |

| | | | | |
|---|---|---|---|---|
| | Dummy<br>KNN | 20<br>25 | -2.709 | *0.042 |
| Subject E | SVM (poly)<br>SVM (RBF) | 42<br>42 | 0.653 | 0.542 |
| | SVM (poly)<br>Random Forest | 43<br>36 | 1.687 | 0.152 |
| | SVM (poly)<br>LDA | 42<br>38 | 2.707 | *0.042 |
| | SVM (poly)<br>KNN | 42<br>27 | 6.744 | 0.001 |
| | SVM (RBF)<br>Random Forest | 43<br>37 | 1.549 | 0.182 |
| | SVM (RBF)<br>LDA | 42<br>38 | 2.927 | *0.033 |
| | SVM (RBF)<br>KNN | 42<br>27 | 6.947 | **0.001 |
| | Random Forest<br>LDA | 38<br>37 | 0.470 | 0.658 |
| | Random Forest<br>KNN | 36<br>27 | 3.865 | *0.012 |
| | LDA<br>KNN | 37<br>27 | 3.751 | *0.013 |
| | Dummy<br>SVM (poly) | 22<br>42 | -12.390 | ***0.000 |
| | Dummy<br>SVM (RBF) | 22<br>43 | -11.214 | ***0.000 |
| | Dummy<br>Random Forest | 21<br>37 | -11.718 | ***0.000 |
| | Dummy<br>LDA | 21<br>37 | -5.937 | **0.002 |
| | Dummy<br>KNN | 21<br>27 | -1.235 | 0.272 |
| Subject F | SVM (poly)<br>SVM (RBF) | 29<br>30 | -0.953 | 0.384 |
| | SVM (poly)<br>Random Forest | 30<br>28 | 0.971 | 0.465 |
| | SVM (poly)<br>LDA | 30<br>27 | 0.215 | 0.838 |
| | SVM (poly)<br>KNN | 30<br>21 | 2.439 | 0.059 |
| | SVM (RBF)<br>Random Forest | 28<br>27 | 1.208 | 0.281 |
| | SVM (RBF)<br>LDA | 30<br>26 | 0.379 | 0.721 |
| | SVM (RBF)<br>KNN | 29<br>21 | 2.762 | *0.040 |
| | Random Forest<br>LDA | 28<br>26 | -0.235 | 0.823 |
| | Random Forest<br>KNN | 28<br>22 | 2.264 | 0.073 |
| | LDA<br>KNN | 26<br>21 | 2.972 | *0.031 |
| | Dummy<br>SVM (poly) | 19<br>31 | -3.678 | *0.014 |
| | Dummy<br>SVM (RBF) | 20<br>30 | -4.438 | **0.007 |
| | Dummy<br>Random Forest | 20<br>27 | -3.493 | *0.017 |
| | Dummy<br>LDA | 20<br>27 | -2.508 | 0.054 |
| | Dummy<br>KNN | 20<br>21 | 0.426 | 0.688 |
| Subject G | SVM (poly)<br>SVM (RBF) | 30<br>30 | 0.526 | 0.621 |
| | SVM (poly)<br>Random Forest | 28<br>28 | 1.265 | 0.262 |

| | | | | |
|---|---|---|---|---|
| | SVM (poly)<br>LDA | 29<br>28 | 1.651 | 0.160 |
| | SVM (poly)<br>KNN | 30<br>23 | 6.007 | **0.002 |
| | SVM (RBF)<br>Random Forest | 29<br>28 | 1.165 | 0.297 |
| | SVM (RBF)<br>LDA | 29<br>27 | 1.905 | 0.115 |
| | SVM (RBF)<br>KNN | 28<br>24 | 3.715 | *0.014 |
| | Random Forest<br>LDA | 28<br>27 | 1.342 | 0.237 |
| | Random Forest<br>KNN | 28<br>23 | 3.563 | *0.016 |
| | LDA<br>KNN | 27<br>23 | 1.532 | 0.186 |
| | Dummy<br>SVM (poly) | 21<br>29 | -9.333 | ***0.000 |
| | Dummy<br>SVM (RBF) | 21<br>29 | -6.597 | **0.001 |
| | Dummy<br>Random Forest | 21<br>29 | -6.066 | **0.002 |
| | Dummy<br>LDA | 20<br>28 | -3.826 | *0.012 |
| | Dummy<br>KNN | 21<br>24 | -3.361 | *0.020 |
| Subject H | SVM (poly)<br>SVM (RBF) | 22<br>21 | 0.481 | 0.651 |
| | SVM (poly)<br>Random Forest | 22<br>21 | -0.173 | 0.870 |
| | SVM (poly)<br>LDA | 22<br>21 | 0.489 | 0.645 |
| | SVM (poly)<br>KNN | 21<br>19 | 1.190 | 0.287 |
| | SVM (RBF)<br>Random Forest | 21<br>21 | -0.364 | 0.731 |
| | SVM (RBF)<br>LDA | 20<br>21 | 0.364 | 0.730 |
| | SVM (RBF)<br>KNN | 20<br>20 | 1.267 | 0.261 |
| | Random Forest<br>LDA | 20<br>21 | 0.552 | 0.605 |
| | Random Forest<br>KNN | 21<br>20 | 1.197 | 0.285 |
| | LDA<br>KNN | 21<br>20 | 1.022 | 0.354 |
| | Dummy<br>SVM (poly) | 19<br>22 | -1.250 | 0.267 |
| | Dummy<br>SVM (RBF) | 21<br>21 | -1.038 | 0.347 |
| | Dummy<br>Random Forest | 20<br>21 | -1.983 | 0.104 |
| | Dummy<br>LDA | 22<br>21 | -0.768 | 0.477 |
| | Dummy<br>KNN | 20<br>19 | 0.345 | 0.744 |
| Subject I | SVM (poly)<br>SVM (RBF) | 29<br>30 | -1.291 | 0.253 |
| | SVM (poly)<br>Random Forest | 29<br>31 | 0.686 | 0.523 |
| | SVM (poly)<br>LDA | 29<br>29 | 1.774 | 0.136 |
| | SVM (poly)<br>KNN | 30<br>23 | 4.814 | **0.005 |
| | SVM (RBF)<br>Random Forest | 31<br>31 | 1.633 | 0.163 |

| | | | | |
|---|---|---|---|---|
| | SVM (RBF)<br>LDA | 31<br>29 | 3.017 | *0.030 |
| | SVM (RBF)<br>KNN | 30<br>25 | 6.736 | **0.001 |
| | Random Forest<br>LDA | 31<br>29 | 0.287 | 0.786 |
| | Random Forest<br>KNN | 30<br>24 | 5.116 | **0.004 |
| | LDA<br>KNN | 29<br>23 | 6.940 | **0.001 |
| | Dummy<br>SVM (poly) | 21<br>29 | -6.817 | **0.001 |
| | Dummy<br>SVM (RBF) | 22<br>31 | -8.368 | ***0.000 |
| | Dummy<br>Random Forest | 21<br>30 | -6.460 | **0.001 |
| | Dummy<br>LDA | 21<br>28 | -9.573 | ***0.000 |
| | Dummy<br>KNN | 21<br>24 | 0.000 | 1.000 |

**Table 6S,** “5x2 Cross-Validation Test” in Time-frequency Domain (Wigner), After PCA (150) scenario, SVM (kernel='poly', degree = 3, C = 100000),
SVM (kernel='rbf', gamma=0.001, C=100),
RandomForestClassifier (max_depth=5, random_state=20),
KNeighborsClassifier (n_neighbors = 60).

| Subjects | Classifiers | Accuracies % | t Statistics | P-Value |
|---|---|---|---|---|
| Subject B | SVM (poly)<br>SVM (RBF) | 28<br>32 | -1.714 | 0.147 |
| | SVM (poly)<br>Random Forest | 28<br>28 | -1.280 | 0.257 |
| | SVM (poly)<br>LDA | 28<br>33 | -2.020 | 0.090 |
| | SVM (poly)<br>KNN | 28<br>25 | 2.267 | 0.073 |
| | SVM (RBF)<br>Random Forest | 33<br>28 | 2.252 | 0.074 |
| | SVM (RBF)<br>LDA | 31<br>32 | -0.809 | 0.456 |
| | SVM (RBF)<br>KNN | 32<br>26 | 3.190 | *0.024 |
| | Random Forest<br>LDA | 28<br>32 | -3.891 | *0.012 |
| | Random Forest<br>KNN | 28<br>25 | 3.351 | *0.020 |
| | LDA<br>KNN | 32<br>25 | 3.844 | *0.012 |
| | Dummy<br>SVM (poly) | 21<br>28 | -2.805 | *0.038 |
| | Dummy<br>SVM (RBF) | 21<br>31 | -3.810 | *0.013 |
| | Dummy<br>Random Forest | 21<br>28 | -5.213 | **0.003 |
| | Dummy<br>LDA | 21<br>32 | -6.520 | **0.001 |
| | Dummy<br>KNN | 21<br>25 | -0.737 | 0.494 |

| | | | | |
|---|---|---|---|---|
| Subject C | SVM (poly)<br>SVM (RBF) | 33<br>39 | -1.486 | 0.197 |
| | SVM (poly)<br>Random Forest | 32<br>33 | 0.825 | 0.447 |
| | SVM (poly)<br>LDA | 33<br>39 | -1.183 | 0.290 |
| | SVM (poly)<br>KNN | 34<br>27 | 1.819 | 0.129 |
| | SVM (RBF)<br>Random Forest | 38<br>33 | 2.844 | *0.036 |
| | SVM (RBF)<br>LDA | 39<br>40 | 0.833 | 0.443 |
| | SVM (RBF)<br>KNN | 40<br>27 | 3.159 | *0.025 |
| | Random Forest<br>LDA | 33<br>39 | -1.699 | 0.150 |
| | Random Forest<br>KNN | 33<br>27 | 1.316 | 0.245 |
| | LDA<br>KNN | 39<br>27 | 1.695 | 0.151 |
| | Dummy<br>SVM (poly) | 19<br>33 | -3.005 | *0.030 |
| | Dummy<br>SVM (RBF) | 19<br>40 | -5.690 | **0.002 |
| | Dummy<br>Random Forest | 21<br>33 | -4.648 | **0.006 |

| | | | | |
|---|---|---|---|---|
| | Dummy<br>LDA | 20<br>38 | -7.234 | **0.001 |
| | Dummy<br>KNN | 20<br>26 | -2.698 | *0.043 |
| Subject E | SVM (poly)<br>SVM (RBF) | 38<br>40 | -2.152 | 0.084 |
| | SVM (poly)<br>Random Forest | 38<br>35 | 3.401 | *0.019 |
| | SVM (poly)<br>LDA | 38<br>41 | -1.859 | 0.122 |
| | SVM (poly)<br>KNN | 38<br>32 | 4.700 | **0.005 |
| | SVM (RBF)<br>Random Forest | 40<br>36 | 3.473 | *0.018 |
| | SVM (RBF)<br>LDA | 41<br>42 | -0.238 | 0.822 |
| | SVM (RBF)<br>KNN | 41<br>33 | 4.040 | *0.010 |
| | Random Forest<br>LDA | 35<br>42 | -2.243 | 0.075 |
| | Random Forest<br>KNN | 35<br>32 | 0.891 | 0.414 |
| | LDA<br>KNN | 41<br>33 | 4.622 | **0.006 |
| | Dummy<br>SVM (poly) | 21<br>38 | -11.319 | ***0.000 |
| | Dummy<br>SVM (RBF) | 21<br>40 | -12.669 | ***0.000 |
| | Dummy<br>Random Forest | 21<br>35 | -9.668 | ***0.000 |
| | Dummy<br>LDA | 21<br>41 | -10.052 | ***0.000 |
| | Dummy<br>KNN | 22<br>32 | -4.900 | **0.004 |
| Subject F | SVM (poly)<br>SVM (RBF) | 26<br>30 | -0.742 | 0.492 |
| | SVM (poly)<br>Random Forest | 26<br>26 | 0.472 | 0.657 |
| | SVM (poly)<br>LDA | 24<br>28 | -2.957 | *0.032 |
| | SVM (poly)<br>KNN | 25<br>23 | 0.562 | 0.598 |
| | SVM (RBF)<br>Random Forest | 29<br>26 | 0.469 | 0.659 |
| | SVM (RBF)<br>LDA | 29<br>28 | -1.548 | 0.182 |
| | SVM (RBF)<br>KNN | 28<br>25 | 2.952 | *0.032 |
| | Random Forest<br>LDA | 27<br>29 | -1.007 | 0.360 |
| | Random Forest<br>KNN | 26<br>24 | 2.075 | 0.093 |
| | LDA<br>KNN | 28<br>23 | 4.904 | **0.004 |
| | Dummy<br>SVM (poly) | 20<br>26 | -0.892 | 0.413 |
| | Dummy<br>SVM (RBF) | 20<br>28 | -6.069 | **0.002 |
| | Dummy<br>Random Forest | 19<br>25 | -6.484 | **0.001 |
| | Dummy<br>LDA | 20<br>29 | -4.984 | **0.004 |
| | Dummy<br>KNN | 20<br>23 | -0.847 | 0.436 |
| Subject G | SVM (poly)<br>SVM (RBF) | 26<br>28 | -1.769 | 0.137 |

| | | | | |
|---|---|---|---|---|
| | SVM (poly)<br>Random Forest | 25<br>25 | -0.242 | 0.819 |
| | SVM (poly)<br>LDA | 25<br>30 | -1.371 | 0.229 |
| | SVM (poly)<br>KNN | 26<br>25 | 0.613 | 0.567 |
| | SVM (RBF)<br>Random Forest | 29<br>26 | 3.062 | *0.028 |
| | SVM (RBF)<br>LDA | 29<br>30 | 1.077 | 0.330 |
| | SVM (RBF)<br>KNN | 28<br>25 | 2.650 | *0.045 |
| | Random Forest<br>LDA | 26<br>30 | -1.592 | 0.172 |
| | Random Forest<br>KNN | 26<br>25 | 0.571 | 0.593 |
| | LDA<br>KNN | 29<br>24 | 3.070 | *0.028 |
| | Dummy<br>SVM (poly) | 21<br>26 | -2.145 | 0.085 |
| | Dummy<br>SVM (RBF) | 21<br>29 | -10.710 | ***0.000 |
| | Dummy<br>Random Forest | 21<br>25 | -5.522 | **0.003 |
| | Dummy<br>LDA | 21<br>30 | -5.681 | **0.002 |
| | Dummy<br>KNN | 20<br>25 | -2.063 | 0.094 |
| Subject H | SVM (poly)<br>SVM (RBF) | 20<br>19 | -0.608 | 0.570 |
| | SVM (poly)<br>Random Forest | 20<br>20 | -0.099 | 0.925 |
| | SVM (poly)<br>LDA | 21<br>21 | 0.297 | 0.779 |
| | SVM (poly)<br>KNN | 21<br>19 | 2.139 | 0.085 |
| | SVM (RBF)<br>Random Forest | 19<br>20 | 0.391 | 0.712 |
| | SVM (RBF)<br>LDA | 21<br>21 | 0.951 | 0.385 |
| | SVM (RBF)<br>KNN | 20<br>19 | 1.307 | 0.248 |
| | Random Forest<br>LDA | 21<br>21 | -2.210 | 0.078 |
| | Random Forest<br>KNN | 20<br>19 | 1.001 | 0.363 |
| | LDA<br>KNN | 20<br>18 | 1.123 | 0.312 |
| | Dummy<br>SVM (poly) | 20<br>20 | -1.388 | 0.224 |
| | Dummy<br>SVM (RBF) | 20<br>21 | -1.657 | 0.158 |
| | Dummy<br>Random Forest | 21<br>19 | -0.727 | 0.500 |
| | Dummy<br>LDA | 21<br>21 | -1.174 | 0.239 |
| | Dummy<br>KNN | 21<br>18 | -0.166 | 0.875 |
| Subject I | SVM (poly)<br>SVM (RBF) | 28<br>31 | -1.780 | 0.135 |
| | SVM (poly)<br>Random Forest | 28<br>28 | 0.665 | 0.535 |
| | SVM (poly)<br>LDA | 28<br>30 | -2.231 | 0.076 |
| | SVM (poly)<br>KNN | 29<br>25 | 2.270 | 0.072 |

| | | | | |
|---|---|---|---|---|
| | SVM (RBF)<br>Random Forest | 30<br>27 | 2.418 | 0.060 |
| | SVM (RBF)<br>LDA | 31<br>30 | 1.081 | 0.329 |
| | SVM (RBF)<br>KNN | 30<br>26 | 3.475 | *0.018 |
| | Random Forest<br>LDA | 28<br>30 | -2.114 | 0.088 |
| | Random Forest<br>KNN | 28<br>26 | 1.498 | 0.194 |
| | LDA<br>KNN | 31<br>25 | 3.097 | *0.027 |
| | Dummy<br>SVM (poly) | 21<br>28 | -2.761 | *0.040 |
| | Dummy<br>SVM (RBF) | 22<br>31 | -5.360 | **0.003 |
| | Dummy<br>Random Forest | 21<br>27 | -4.456 | **0.007 |
| | Dummy<br>LDA | 21<br>31 | -6.076 | **0.002 |
| | Dummy<br>KNN | 20<br>25 | -1.169 | 0.295 |
| Independent | SVM (poly)<br>SVM (RBF) | 26<br>29 | -4.571 | 0.006** |
| | SVM (poly)<br>Random Forest | 26<br>26 | 0.019 | 0.986 |
| | SVM (poly)<br>LDA | 26<br>29 | -1.756 | 0.139 |
| | SVM (poly)<br>KNN | 27<br>25 | 1.795 | 0.133 |
| | SVM (RBF)<br>Random Forest | 26<br>26 | 4.449 | 0.007** |
| | SVM (RBF)<br>LDA | 29<br>29 | 2.084 | 0.092 |
| | SVM (RBF)<br>KNN | 29<br>25 | 4.653 | 0.006** |
| | Random Forest<br>LDA | 26<br>29 | -1.110 | 0.318 |
| | Random Forest<br>KNN | 26<br>25 | 1.011 | 0.358 |
| | LDA<br>KNN | 29<br>25 | 3.028 | 0.029* |
| | Dummy<br>SVM (poly) | 21<br>27 | -3.350 | 0.020* |
| | Dummy<br>SVM (RBF) | 21<br>29 | -9.051 | 0.000*** |
| | Dummy<br>Random Forest | 21<br>27 | -3.886 | 0.012* |
| | Dummy<br>LDA | 21<br>29 | -5.018 | 0.004** |
| | Dummy<br>KNN | 21<br>25 | -2.513 | 0.054 |

Table 7S, 10-fold cross validation, Accuracies (%) and Training Times (Second) for each fold in Time domain, Before PCA scenario.

| | | | | | | | | | | | |
|---|---|---|---|---|---|---|---|---|---|---|---|
| Subject B | SVM (Poly) | Accuracy | 45.50 | 40.74 | 45.50 | 39.68 | 43.38 | 40.74 | 40.21 | 38.09 | 41.79 | 37.56 |
| | | Time | 2.08 | 1.76 | 1.62 | 1.79 | 1.95 | 2.11 | 1.99 | 1.81 | 1.79 | 1.74 |
| | SVM (RBF) | Accuracy | 50.79 | 46.56 | 46.03 | 50.79 | 52.91 | 42.85 | 47.61 | 44.97 | 42.85 | 39.68 |
| | | Time | 0.97 | 0.76 | 0.80 | 0.80 | 0.78 | 0.85 | 1.02 | 0.85 | 0.92 | 0.80 |
| | Random Forest | Accuracy | 40.74 | 32.80 | 38.62 | 36.50 | 39.68 | 38.09 | 42.32 | 39.15 | 40.74 | 31.21 |
| | | Time | 0.0 | 0.0 | 0.0 | 0.0 | 0.0 | 0.0 | 0.0 | 0.0 | 0.0 | 0.0 |
| | LDA | Accuracy | 37.56 | 32.80 | 40.74 | 35.97 | 43.91 | 36.50 | 35.97 | 31.21 | 31.74 | 38.09 |
| | | Time | 0.25 | 0.26 | 0.25 | 0.24 | 0.25 | 0.29 | 0.28 | 0.25 | 0.26 | 0.24 |
| | KNN | Accuracy | 37.03 | 37.56 | 34.39 | 36.50 | 38.62 | 34.39 | 40.74 | 34.39 | 31.21 | 34.92 |
| | | Time | 0.0 | 0.0 | 0.0 | 0.0 | 0.0 | 0.0 | 0.0 | 0.0 | 0.0 | 0.0 |
| | Dummy | Accuracy | 22.75 | 19.57 | 21.16 | 23.28 | 26.45 | 18.51 | 18.51 | 21.69 | 22.22 | 24.33 |
| | | Time | | | | | | | | | | |
| Subject C | SVM (Poly) | Accuracy | 58.51 | 60.63 | 54.25 | 50 | 59.57 | 53.19 | 53.19 | 61.70 | 52.12 | 53.76 |
| | | Time | 0.18 | 0.22 | 0.18 | 0.18 | 0.20 | 0.18 | 0.20 | 0.20 | 0.18 | 0.18 |
| | SVM (RBF) | Accuracy | 57.44 | 61.70 | 55.31 | 58.51 | 57.44 | 53.19 | 59.57 | 68.08 | 55.31 | 53.76 |
| | | Time | 0.14 | 0.18 | 0.18 | 0.20 | 0.18 | 0.18 | 0.17 | 0.19 | 0.19 | 0.18 |
| | Random Forest | Accuracy | 44.68 | 41.48 | 48.93 | 42.55 | 45.74 | 45.74 | 46.80 | 40.42 | 39.36 | 45.16 |
| | | Time | 0.0 | 0.0 | 0.0 | 0.0 | 0.0 | 0.0 | 0.0 | 0.0 | 0.0 | 0.0 |
| | LDA | Accuracy | 39.36 | 39.36 | 28.72 | 39.36 | 37.23 | 31.91 | 35.10 | 38.29 | 38.29 | 46.23 |
| | | Time | 0.16 | 0.15 | 0.20 | 0.15 | 0.15 | 0.18 | 0.14 | 0.14 | 0.14 | 0.15 |
| | KNN | Accuracy | 41.48 | 54.25 | 41.48 | 45.74 | 50 | 36.17 | 38.29 | 56.38 | 46.80 | 47.11 |
| | | Time | 0.0 | 0.0 | 0.0 | 0.0 | 0.0 | 0.0 | 0.0 | 0.0 | 0.0 | 0.0 |
| | Dummy | Accuracy | 25.53 | 20.21 | 23.40 | 24.46 | 22.34 | 14.89 | 20.21 | 22.34 | 19.14 | 24.73 |
| | | Time | | | | | | | | | | |
| Subject E | SVM (Poly) | Accuracy | 57.54 | 59.64 | 60.21 | 56.33 | 56.69 | 55.98 | 61.98 | 58.09 | 55.98 | 53.87 |
| | | Time | 2.54 | 2.85 | 2.57 | 2.43 | 3.49 | 2.43 | 3.22 | 2.76 | 2.57 | 2.43 |
| | SVM (RBF) | Accuracy | 60.35 | 63.85 | 61.97 | 60.56 | 61.26 | 57.74 | 63.38 | 58.45 | 60.56 | 58.80 |
| | | Time | 1.77 | 1.97 | 1.74 | 1.64 | 1.80 | 2.14 | 2.24 | 1.80 | 1.85 | 1.69 |
| | Random Forest | Accuracy | 40.35 | 43.15 | 43.66 | 43.66 | 44.36 | 46.83 | 44.71 | 42.95 | 44.36 | 46.12 |

| | | | | | | | | | | | |
|---|---|---|---|---|---|---|---|---|---|---|---|
| | | Time | 0.0 | 0.0 | 0.0 | 0.0 | 0.0 | 0.0 | 0.0 | 0.0 | 0.0 | 0.0 |
| | LDA | Accuracy | 54.38 | 54.73 | 56.33 | 54.22 | 53.52 | 51.76 | 55.98 | 55.98 | 53.52 | 54.22 |
| | | Time | 0.49 | 0.40 | 0.34 | 0.47 | 0.49 | 0.49 | 0.41 | 0.36 | 0.36 | 0.37 |
| | KNN | Accuracy | 47.01 | 51.92 | 45.07 | 46.83 | 50.70 | 50.00 | 46.47 | 44.36 | 53.87 | 42.95 |
| | | Time | 0.0 | 0.0 | 0.01 | 0.0 | 0.00 | 0.00 | 0.0 | 0.0 | 0.0 | 0.0 |
| | Dummy | Accuracy | 23.15 | 20.00 | 20.77 | 20.77 | 21.12 | 23.23 | 22.18 | 22.18 | 21.83 | 22.88 |
| | | Time | | | | | | | | | | |
| Subject F | SVM (Poly) | Accuracy | 48.95 | 41.66 | 36.45 | 38.54 | 48.95 | 50.00 | 33.33 | 40.00 | 34.73 | 40.00 |
| | | Time | 0.28 | 0.26 | 0.24 | 0.26 | 0.28 | 0.28 | 0.24 | 0.26 | 0.28 | 0.26 |
| | SVM (RBF) | Accuracy | 48.95 | 42.70 | 39.58 | 40.62 | 48.95 | 48.95 | 41.66 | 42.10 | 40.00 | 38.94 |
| | | Time | 0.18 | 0.20 | 0.20 | 0.21 | 0.21 | 0.22 | 0.20 | 0.20 | 0.20 | 0.21 |
| | Random Forest | Accuracy | 41.66 | 39.58 | 36.45 | 45.83 | 50.00 | 51.04 | 34.37 | 32.63 | 32.63 | 42.10 |
| | | Time | 0.0 | 0.0 | 0.0 | 0.0 | 0.0 | 0.0 | 0.0 | 0.0 | 0.0 | 0.0 |
| | LDA | Accuracy | 33.33 | 29.16 | 28.12 | 29.16 | 32.29 | 27.08 | 21.87 | 26.31 | 24.21 | 30.52 |
| | | Time | 0.17 | 0.14 | 0.14 | 0.14 | 0.14 | 0.15 | 0.15 | 0.17 | 0.15 | 0.15 |
| | KNN | Accuracy | 37.50 | 37.50 | 34.37 | 33.33 | 41.66 | 45.83 | 33.33 | 28.42 | 32.63 | 32.63 |
| | | Time | 0.0 | 0.0 | 0.0 | 0.0 | 0.0 | 0.0 | 0.0 | 0.0 | 0.0 | 0.0 |
| | Dummy | Accuracy | 19.79 | 26.04 | 17.70 | 21.87 | 26.04 | 26.24 | 17.70 | 20.00 | 21.05 | 21.05 |
| | | Time | | | | | | | | | | |
| Subject G | SVM (Poly) | Accuracy | 41.57 | 40.00 | 37.36 | 40.52 | 37.36 | 39.68 | 38.09 | 42.85 | 42.85 | 39.68 |
| | | Time | 1.89 | 1.90 | 1.88 | 1.91 | 1.72 | 1.90 | 1.93 | 1.89 | 1.59 | 1.81 |
| | SVM (RBF) | Accuracy | 44.73 | 40.52 | 39.47 | 45.26 | 39.47 | 38.09 | 45.50 | 43.91 | 37.56 | 43.91 |
| | | Time | 0.82 | 0.82 | 0.83 | 0.83 | 0.82 | 0.82 | 0.87 | 0.82 | 0.80 | 0.82 |
| | Random Forest | Accuracy | 41.05 | 34.73 | 38.42 | 37.89 | 28.94 | 35.97 | 37.03 | 37.03 | 31.21 | 40.74 |
| | | Time | 0.0 | 0.0 | 0.0 | 0.0 | 0.0 | 0.0 | 0.0 | 0.0 | 0.0 | 0.0 |
| | LDA | Accuracy | 36.31 | 34.21 | 32.10 | 33.68 | 29.47 | 33.86 | 32.80 | 37.56 | 29.62 | 31.74 |
| | | Time | 0.24 | 0.28 | 0.24 | 0.23 | 0.25 | 0.24 | 0.25 | 0.24 | 0.24 | 0.24 |
| | KNN | Accuracy | 30.52 | 31.57 | 31.57 | 34.21 | 29.47 | 25.92 | 35.44 | 36.50 | 33.86 | 31.74 |
| | | Time | 0.0 | 0.0 | 0.0 | 0.0 | 0.0 | 0.0 | 0.0 | 0.0 | 0.0 | 0.0 |
| | Dummy | Accuracy | 20.52 | 19.47 | 25.26 | 23.15 | 21.57 | 20.10 | 22.22 | 24.86 | 21.16 | 20.63 |

| | | | | | | | | | | | |
|---|---|---|---|---|---|---|---|---|---|---|---|
| | | Time | | | | | | | | | |
| Subject H | SVM (Poly) | Accuracy | 23.91 | 26.08 | 22.82 | 28.26 | 21.73 | 20.65 | 26.08 | 22.82 | 28.26 | 25.27 |
| | | Time | 0.57 | 0.78 | 0.68 | 0.81 | 0.70 | 0.71 | 0.74 | 0.70 | 0.70 | 0.71 |
| | SVM (RBF) | Accuracy | 19.56 | 26.08 | 19.56 | 23.91 | 20.65 | 17.39 | 15.21 | 21.73 | 30.34 | 26.37 |
| | | Time | 0.21 | 0.25 | 0.30 | 0.26 | 0.30 | 0.26 | 0.28 | 0.25 | 0.28 | 0.26 |
| | Random Forest | Accuracy | 26.08 | 20.65 | 13.04 | 19.56 | 26.08 | 17.39 | 20.65 | 22.82 | 29.34 | 28.57 |
| | | Time | 0.0 | 0.0 | 0.0 | 0.0 | 0.0 | 0.0 | 0.0 | 0.0 | 0.0 | 0.0 |
| | LDA | Accuracy | 23.91 | 30.43 | 22.82 | 22.82 | 19.56 | 18.47 | 16.30 | 26.08 | 19.56 | 20.87 |
| | | Time | 0.18 | 0.15 | 0.17 | 0.21 | 0.15 | 0.15 | 0.14 | 0.17 | 0.15 | 0.14 |
| | KNN | Accuracy | 26.08 | 19.56 | 17.39 | 18.47 | 25.00 | 20.65 | 26.08 | 17.39 | 20.65 | 17.58 |
| | | Time | 0.0 | 0.0 | 0.0 | 0.0 | 0.0 | 0.0 | 0.0 | 0.0 | 0.0 | 0.0 |
| | Dummy | Accuracy | 27.17 | 16.30 | 15.21 | 22.82 | 10.86 | 18.47 | 17.39 | 21.73 | 22.82 | 26.37 |
| | | Time | | | | | | | | | | |
| Subject I | SVM (Poly) | Accuracy | 39.06 | 35.60 | 36.64 | 32.46 | 33.50 | 36.64 | 33.50 | 36.64 | 32.98 | 42.40 |
| | | Time | 4.66 | 5.39 | 5.31 | 6.27 | 5.12 | 5.46 | 4.92 | 4.41 | 5.21 | 5.14 |
| | SVM (RBF) | Accuracy | 42.18 | 38.74 | 48.69 | 44.50 | 40.31 | 46.07 | 36.64 | 40.31 | 37.69 | 40.83 |
| | | Time | 0.89 | 0.95 | 0.96 | 0.94 | 0.94 | 0.94 | 0.92 | 0.89 | 0.93 | 0.92 |
| | Random Forest | Accuracy | 38.54 | 39.26 | 43.97 | 33.50 | 38.21 | 41.88 | 37.17 | 34.03 | 33.50 | 33.50 |
| | | Time | 0.0 | 0.0 | 0.0 | 0.0 | 0.0 | 0.0 | 0.0 | 0.0 | 0.0 | 0.0 |
| | LDA | Accuracy | 39.06 | 34.55 | 35.60 | 29.84 | 34.55 | 38.74 | 31.41 | 35.60 | 32.98 | 42.40 |
| | | Time | 0.28 | 0.24 | 0.24 | 0.24 | 0.23 | 0.23 | 0.24 | 0.23 | 0.24 | 0.23 |
| | KNN | Accuracy | 34.89 | 37.69 | 37.17 | 32.46 | 34.55 | 35.07 | 35.07 | 41.36 | 30.89 | 35.60 |
| | | Time | 0.0 | 0.0 | 0.0 | 0.0 | 0.0 | 0.0 | 0.0 | 0.0 | 0.0 | 0.0 |
| | Dummy | Accuracy | 21.35 | 21.46 | 21.46 | 14.13 | 26.17 | 18.84 | 26.17 | 27.22 | 17.27 | 24.08 |
| | | Time | | | | | | | | | | |

Table 8S, 10-fold cross validation, Accuracies (%) and Training Times (Second) for each fold in Time domain, After PCA (100) scenario.

| | | | | | | | | | | | |
|---|---|---|---|---|---|---|---|---|---|---|---|
| Subject B | SVM (Poly) | Accuracy | 41.26 | 39.15 | 36.50 | 39.68 | 42.85 | 30.15 | 38.09 | 42.85 | 36.50 | 39.68 |
| | | Time | 0.43 | 0.45 | 0.42 | 0.43 | 0.42 | 0.41 | 0.43 | 0.43 | 0.42 | 0.43 |
| | SVM (RBF) | Accuracy | 51.32 | 47.08 | 46.56 | 49.73 | 53.43 | 44.44 | 48.67 | 46.03 | 43.38 | 40.21 |
| | | Time | 0.34 | 0.37 | 0.40 | 0.34 | 0.33 | 0.36 | 0.34 | 0.33 | 0.36 | 0.36 |
| | Random Forest | Accuracy | 39.15 | 36.50 | 39.68 | 39.15 | 44.44 | 37.56 | 38.62 | 37.56 | 44.44 | 35.44 |
| | | Time | 0.0 | 0.0 | 0.0 | 0.0 | 0.0 | 0.0 | 0.0 | 0.0 | 0.0 | 0.0 |
| | LDA | Accuracy | 51.32 | 43.91 | 47.08 | 49.20 | 50.26 | 43.38 | 49.20 | 44.44 | 42.32 | 42.32 |
| | | Time | 0.03 | 0.03 | 0.03 | 0.01 | 0.01 | 0.03 | 0.03 | 0.03 | 0.01 | 0.01 |
| | KNN | Accuracy | 37.56 | 38.09 | 33.33 | 38.09 | 39.15 | 35.97 | 41.26 | 33.33 | 29.62 | 32.80 |
| | | Time | 0.0 | 0.0 | 0.0 | 0.0 | 0.0 | 0.0 | 0.0 | 0.0 | 0.0 | 0.0 |
| | Dummy | Accuracy | 22.75 | 19.57 | 21.16 | 23.28 | 26.45 | 18.51 | 18.51 | 21.69 | 22.22 | 24.33 |
| | | Time | | | | | | | | | | |
| Subject C | SVM (Poly) | Accuracy | 50.00 | 51.06 | 46.80 | 47.87 | 51.06 | 50.00 | 44.68 | 59.57 | 48.93 | 48.38 |
| | | Time | 0.10 | 0.11 | 0.11 | 0.09 | 0.09 | 0.10 | 0.09 | 0.09 | 0.10 | 0.10 |
| | SVM (RBF) | Accuracy | 58.51 | 59.57 | 54.25 | 57.44 | 56.38 | 54.25 | 62.76 | 67.02 | 51.06 | 53.76 |
| | | Time | 0.07 | 0.12 | 0.11 | 0.12 | 0.09 | 0.12 | 0.10 | 0.10 | 0.09 | 0.09 |
| | Random Forest | Accuracy | 44.68 | 42.55 | 41.48 | 51.06 | 52.12 | 39.36 | 47.87 | 44.68 | 53.19 | 43.01 |
| | | Time | 0.0 | 0.0 | 0.0 | 0.0 | 0.0 | 0.0 | 0.0 | 0.0 | 0.0 | 0.0 |
| | LDA | Accuracy | 61.70 | 61.70 | 60.63 | 62.76 | 59.57 | 63.82 | 59.57 | 62.76 | 60.63 | 52.68 |
| | | Time | 0.01 | 0.01 | 0.01 | 0.01 | 0.03 | 0.01 | 0.01 | 0.01 | 0.01 | 0.00 |
| | KNN | Accuracy | 40.42 | 54.25 | 45.74 | 44.68 | 44.68 | 39.36 | 37.23 | 51.06 | 48.93 | 44.08 |
| | | Time | 0.0 | 0.0 | 0.0 | 0.0 | 0.0 | 0.0 | 0.0 | 0.0 | 0.0 | 0.0 |
| | Dummy | Accuracy | 25.53 | 20.21 | 23.40 | 24.46 | 22.34 | 14.89 | 20.21 | 22.34 | 19.14 | 24.73 |
| | | Time | | | | | | | | | | |
| Subject E | SVM (Poly) | Accuracy | 55.78 | 57.89 | 54.57 | 53.52 | 54.57 | 51.05 | 53.87 | 50.70 | 54.22 | 51.40 |
| | | Time | 0.91 | 0.88 | 0.89 | 0.90 | 0.90 | 0.87 | 0.85 | 0.89 | 0.84 | 0.92 |
| | SVM (RBF) | Accuracy | 57.89 | 61.75 | 59.85 | 59.50 | 58.45 | 54.92 | 63.02 | 57.04 | 58.80 | 56.33 |
| | | Time | 0.62 | 0.65 | 0.63 | 0.60 | 0.62 | 0.61 | 0.61 | 0.61 | 0.61 | 0.58 |
| | Random Forest | Accuracy | 44.21 | 47.36 | 50.00 | 49.64 | 51.76 | 48.94 | 49.29 | 42.95 | 52.46 | 42.60 |

| | | | | | | | | | | | |
|---|---|---|---|---|---|---|---|---|---|---|---|
| | | Time | 0.0 | 0.0 | 0.0 | 0.0 | 0.0 | 0.0 | 0.0 | 0.0 | 0.0 | 0.0 |
| | LDA | Accuracy | 60.00 | 64.91 | 58.80 | 57.74 | 62.32 | 59.85 | 64.08 | 59.50 | 60.56 | 55.98 |
| | | Time | 0.03 | 0.03 | 0.03 | 0.03 | 0.04 | 0.03 | 0.04 | 0.04 | 0.04 | 0.03 |
| | KNN | Accuracy | 43.85 | 52.28 | 46.47 | 46.83 | 52.11 | 47.18 | 44.36 | 43.30 | 51.05 | 44.36 |
| | | Time | 0.0 | 0.01 | 0.0 | 0.0 | 0.0 | 0.0 | 0.0 | 0.0 | 0.0 | 0.01 |
| | Dummy | Accuracy | 23.15 | 20.00 | 20.77 | 20.77 | 21.12 | 23.23 | 22.18 | 22.18 | 21.83 | 22.88 |
| | | Time | | | | | | | | | | |
| Subject F | SVM (Poly) | Accuracy | 38.54 | 29.16 | 36.45 | 35.41 | 45.83 | 35.41 | 37.50 | 30.52 | 37.89 | 32.63 |
| | | Time | 0.12 | 0.10 | 0.13 | 0.13 | 0.10 | 0.09 | 0.11 | 0.12 | 0.10 | 0.11 |
| | SVM (RBF) | Accuracy | 50.00 | 40.62 | 42.70 | 38.54 | 46.87 | 45.83 | 39.58 | 41.05 | 37.89 | 41.05 |
| | | Time | 0.12 | 0.13 | 0.12 | 0.12 | 0.13 | 0.14 | 0.13 | 0.14 | 0.13 | 0.13 |
| | Random Forest | Accuracy | 40.62 | 36.45 | 29.16 | 34.37 | 40.62 | 35.41 | 34.37 | 35.78 | 35.78 | 42.10 |
| | | Time | 0.0 | 0.0 | 0.0 | 0.0 | 0.0 | 0.0 | 0.0 | 0.0 | 0.0 | 0.0 |
| | LDA | Accuracy | 43.75 | 40.62 | 39.58 | 41.66 | 52.08 | 44.79 | 43.75 | 43.15 | 35.78 | 38.94 |
| | | Time | 0.01 | 0.01 | 0.01 | 0.01 | 0.03 | 0.01 | 0.03 | 0.01 | 0.01 | 0.01 |
| | KNN | Accuracy | 35.41 | 35.41 | 32.29 | 28.12 | 44.79 | 43.75 | 35.41 | 25.26 | 30.52 | 32.63 |
| | | Time | 0.0 | 0.01 | 0.0 | 0.0 | 0.0 | 0.0 | 0.0 | 0.0 | 0.0 | 0.0 |
| | Dummy | Accuracy | 19.79 | 26.04 | 17.70 | 21.87 | 26.04 | 26.04 | 17.70 | 20.00 | 21.05 | 21.05 |
| | | Time | | | | | | | | | | |
| Subject G | SVM (Poly) | Accuracy | 40.00 | 33.15 | 33.15 | 40.00 | 32.10 | 33.33 | 37.56 | 33.86 | 41.79 | 35.97 |
| | | Time | 0.46 | 0.46 | 0.45 | 0.46 | 0.43 | 0.43 | 0.45 | 0.46 | 0.43 | 0.46 |
| | SVM (RBF) | Accuracy | 45.26 | 41.05 | 38.42 | 46.31 | 38.42 | 35.44 | 45.50 | 42.85 | 39.15 | 41.79 |
| | | Time | 0.41 | 0.38 | 0.43 | 0.38 | 0.40 | 0.39 | 0.35 | 0.37 | 0.36 | 0.39 |
| | Random Forest | Accuracy | 36.31 | 35.26 | 33.15 | 34.73 | 32.10 | 31.74 | 40.21 | 34.91 | 33.33 | 33.86 |
| | | Time | 0.0 | 0.0 | 0.0 | 0.0 | 0.0 | 0.0 | 0.0 | 0.0 | 0.0 | 0.0 |
| | LDA | Accuracy | 41.05 | 38.42 | 37.89 | 42.10 | 41.05 | 39.15 | 39.68 | 46.56 | 39.68 | 40.74 |
| | | Time | 0.03 | 0.03 | 0.03 | 0.03 | 0.03 | 0.01 | 0.03 | 0.03 | 0.01 | 0.03 |
| | KNN | Accuracy | 33.68 | 30.52 | 31.57 | 33.15 | 30.00 | 29.62 | 31.21 | 38.62 | 35.97 | 33.33 |
| | | Time | 0.0 | 0.0 | 0.0 | 0.0 | 0.0 | 0.0 | 0.0 | 0.0 | 0.01 | 0.0 |
| | Dummy | Accuracy | 20.52 | 19.47 | 25.26 | 23.15 | 21.57 | 20.10 | 22.22 | 24.86 | 21.16 | 20.63 |

| | | | | | | | | | | | |
|---|---|---|---|---|---|---|---|---|---|---|---|
| | | Time | | | | | | | | | | |
| Subject H | SVM (Poly) | Accuracy | 28.60 | 22.82 | 21.73 | 21.73 | 27.17 | 18.47 | 25.00 | 23.91 | 23.91 | 18.68 |
| | | Time | 0.18 | 0.19 | 0.18 | 0.15 | 0.17 | 0.14 | 0.16 | 0.15 | 0.16 | 0.17 |
| | SVM (RBF) | Accuracy | 19.56 | 26.08 | 18.47 | 22.82 | 19.56 | 16.30 | 16.30 | 21.73 | 29.34 | 27.47 |
| | | Time | 0.12 | 0.13 | 0.13 | 0.17 | 0.16 | 0.15 | 0.14 | 0.16 | 0.15 | 0.12 |
| | Random Forest | Accuracy | 23.91 | 27.17 | 14.13 | 20.65 | 16.30 | 25.00 | 20.65 | 32.60 | 32.60 | 27.47 |
| | | Time | 0.0 | 0.0 | 0.0 | 0.0 | 0.0 | 0.0 | 0.0 | 0.0 | 0.0 | 0.0 |
| | LDA | Accuracy | 25.00 | 27.17 | 15.21 | 23.91 | 21.73 | 18.47 | 28.26 | 26.08 | 34.78 | 34.06 |
| | | Time | 0.01 | 0.01 | 0.01 | 0.01 | 0.01 | 0.01 | 0.01 | 0.01 | 0.01 | 0.01 |
| | KNN | Accuracy | 26.08 | 18.47 | 18.47 | 17.39 | 25.00 | 21.73 | 26.08 | 18.47 | 19.56 | 17.58 |
| | | Time | 0.0 | 0.0 | 0.0 | 0.0 | 0.0 | 0.0 | 0.0 | 0.0 | 0.0 | 0.0 |
| | Dummy | Accuracy | 27.17 | 16.30 | 15.21 | 22.82 | 10.86 | 18.47 | 17.39 | 21.73 | 22.82 | 26.37 |
| | | Time | | | | | | | | | | |
| Subject I | SVM (Poly) | Accuracy | 38.54 | 34.55 | 38.21 | 37.69 | 41.88 | 36.64 | 35.60 | 41.36 | 35.07 | 29.31 |
| | | Time | 0.74 | 0.74 | 0.74 | 0.67 | 0.74 | 0.74 | 0.65 | 0.65 | 0.70 | 0.64 |
| | SVM (RBF) | Accuracy | 41.14 | 38.74 | 49.21 | 45.54 | 40.31 | 45.54 | 37.69 | 38.74 | 37.17 | 41.36 |
| | | Time | 0.37 | 0.45 | 0.42 | 0.39 | 0.40 | 0.44 | 0.37 | 0.36 | 0.43 | 0.38 |
| | Random Forest | Accuracy | 38.54 | 37.17 | 42.40 | 35.07 | 41.36 | 41.36 | 39.26 | 35.07 | 35.60 | 41.36 |
| | | Time | 0.0 | 0.0 | 0.0 | 0.0 | 0.0 | 0.0 | 0.0 | 0.0 | 0.0 | 0.0 |
| | LDA | Accuracy | 40.62 | 42.93 | 43.45 | 36.12 | 40.83 | 50.78 | 40.31 | 42.40 | 39.79 | 49.73 |
| | | Time | 0.01 | 0.03 | 0.01 | 0.01 | 0.03 | 0.01 | 0.03 | 0.03 | 0.03 | 0.01 |
| | KNN | Accuracy | 34.37 | 35.07 | 37.17 | 31.93 | 36.64 | 33.50 | 36.64 | 39.79 | 36.12 | 35.60 |
| | | Time | 0.01 | 0.0 | 0.01 | 0.0 | 0.0 | 0.01 | 0.0 | 0.0 | 0.0 | 0.0 |
| | Dummy | Accuracy | 21.35 | 21.46 | 21.46 | 14.13 | 26.17 | 18.84 | 26.17 | 27.22 | 17.27 | 24.08 |
| | | Time | | | | | | | | | | |
| Independent | SVM (Poly) | Accuracy | 36.53 | 35.65 | 35.03 | 34.00 | 33.12 | 32.59 | 35.77 | 33.03 | 36.56 | 35.15 |
| | | Time | 52.71 | 55.13 | 49.49 | 55.15 | 50.74 | 50.25 | 48.38 | 48.42 | 49.99 | 48.35 |
| | SVM (RBF) | Accuracy | 40.49 | 38.73 | 39.61 | 36.82 | 37.88 | 39.20 | 39.91 | 38.85 | 39.82 | 38.14 |
| | | Time | 18.67 | 18.55 | 17.72 | 16.58 | 17.48 | 16.59 | 16.61 | 16.42 | 16.69 | 16.61 |
| | Random Forest | Accuracy | 34.59 | 36.09 | 32.39 | 35.24 | 33.83 | 32.59 | 34.36 | 35.77 | 37.62 | 32.51 |

| | | | | | | | | | | | | |
|---|---|---|---|---|---|---|---|---|---|---|---|---|
| | | Time | 0.0 | 0.01 | 0.0 | 0.0 | 0.0 | 0.0 | 0.0 | 0.0 | 0.01 | 0.0 |
| | LDA | Accuracy | 45.07 | 41.54 | 41.98 | 37.53 | 38.76 | 40.52 | 41.93 | 41.14 | 41.05 | 39.47 |
| | | Time | 0.15 | 0.13 | 0.14 | 0.16 | 0.14 | 0.14 | 0.15 | 0.14 | 0.15 | 0.15 |
| | KNN | Accuracy | 32.21 | 29.84 | 30.89 | 29.77 | 30.30 | 30.83 | 32.51 | 29.86 | 31.80 | 30.13 |
| | | Time | 0.0 | 0.01 | 0.0 | 0.0 | 0.0 | 0.0 | 0.0 | 0.0 | 0.0 | 0.0 |
| | Dummy | Accuracy | 21.03 | 22.53 | 20.51 | 22.02 | 23.08 | 21.76 | 20.96 | 21.05 | 22.81 | 22.55 |
| | | Time | | | | | | | | | | |

Table 9S, 10-fold cross validation, Accuracies (%) and Training Times (Second) for each fold in Time-Frequency domain (Morlet), Before PCA scenario.

| | | | | | | | | | | | | |
|---|---|---|---|---|---|---|---|---|---|---|---|---|
| Subject B | SVM (Poly) | Accuracy | 23.80 | 23.80 | 26.98 | 24.86 | 22.75 | 25.39 | 27.51 | 21.16 | 28.57 | 24.86 |
| | | Time | 44.77 | 44.46 | 44.22 | 45.46 | 47.18 | 43.81 | 46.38 | 46.62 | 45.66 | 49.26 |
| | SVM (RBF) | Accuracy | 22.75 | 26.98 | 29.10 | 32.80 | 23.80 | 25.92 | 29.62 | 24.86 | 23.80 | 22.22 |
| | | Time | 47.29 | 43.90 | 44.01 | 43.92 | 45.84 | 43.70 | 45.61 | 46.26 | 43.72 | 47.00 |
| | Random Forest | Accuracy | 25.92 | 26.45 | 30.68 | 29.62 | 28.04 | 21.69 | 30.15 | 28.57 | 30.68 | 30.15 |
| | | Time | 0.0 | 0.0 | 0.0 | 0.0 | 0.0 | 0.0 | 0.0 | 0.0 | 0.0 | 0.0 |
| | LDA | Accuracy | 28.57 | 23.80 | 12.16 | 20.63 | 24.33 | 21.16 | 18.51 | 19.04 | 19.04 | 25.39 |
| | | Time | 42.21 | 43.21 | 42.00 | 42.10 | 42.19 | 42.27 | 42.00 | 42.13 | 42.21 | 42.33 |
| | KNN | Accuracy | 20.10 | 23.28 | 21.16 | 23.80 | 21.16 | 22.75 | 29.62 | 21.16 | 29.10 | 22.75 |
| | | Time | 0.03 | 0.04 | 0.06 | 0.04 | 0.06 | 0.04 | 0.04 | 0.03 | 0.04 | 0.07 |
| | Dummy | Accuracy | 22.75 | 19.57 | 21.16 | 23.28 | 26.45 | 18.51 | 18.51 | 21.69 | 22.22 | 24.33 |
| | | Time | | | | | | | | | | |
| Subject C | SVM (Poly) | Accuracy | 30.85 | 38.29 | 44.68 | 35.10 | 27.65 | 41.48 | 32.97 | 32.97 | 28.72 | 33.33 |
| | | Time | 12.46 | 13.01 | 12.76 | 13.06 | 13.55 | 12.48 | 12.28 | 13.13 | 12.78 | 11.10 |
| | SVM (RBF) | Accuracy | 31.91 | 40.42 | 30.85 | 36.17 | 23.40 | 36.17 | 30.85 | 36.17 | 29.78 | 32.25 |
| | | Time | 12.20 | 11.98 | 11.10 | 12.51 | 12.28 | 11.99 | 12.33 | 12.94 | 12.18 | 11.19 |
| | Random Forest | Accuracy | 26.59 | 35.10 | 37.23 | 34.04 | 35.10 | 30.85 | 35.10 | 31.91 | 36.17 | 29.03 |
| | | Time | 0.0 | 0.0 | 0.0 | 0.0 | 0.01 | 0.0 | 0.0 | 0.0 | 0.01 | 0.0 |
| | LDA | Accuracy | 25.53 | 29.78 | 21.27 | 24.46 | 17.02 | 26.59 | 22.34 | 28.72 | 30.85 | 22.58 |

| | | | | | | | | | | | | |
|---|---|---|---|---|---|---|---|---|---|---|---|---|
| | | Time | 5.32 | 4.39 | 4.62 | 5.40 | 4.41 | 5.35 | 5.30 | 4.55 | 5.61 | 4.39 |
| | KNN | Accuracy | 26.98 | 27.65 | 33.00 | 28.59 | 30.15 | 27.65 | 27.65 | 32.97 | 26.59 | 34.40 |
| | | Time | 0.01 | 0.03 | 0.01 | 0.03 | 0.02 | 0.04 | 0.03 | 0.01 | 0.03 | 0.03 |
| | Dummy | Accuracy | 25.53 | 20.21 | 23.40 | 24.46 | 22.34 | 14.89 | 20.21 | 22.34 | 19.14 | 24.73 |
| | | Time | | | | | | | | | | |
| Subject E | SVM (Poly) | Accuracy | 41.75 | 36.84 | 40.14 | 38.73 | 39.43 | 44.01 | 39.08 | 40.49 | 35.21 | 41.19 |
| | | Time | 88.56 | 95.85 | 79.86 | 82.93 | 81.81 | 77.28 | 79.78 | 81.51 | 77.17 | 84.44 |
| | SVM (RBF) | Accuracy | 41.75 | 35.08 | 39.78 | 41.90 | 42.60 | 45.07 | 42.25 | 42.95 | 41.54 | 41.54 |
| | | Time | 84.23 | 87.61 | 91.24 | 75.67 | 76.68 | 75.01 | 75.79 | 75.71 | 74.35 | 75.75 |
| | Random Forest | Accuracy | 37.19 | 35.43 | 39.43 | 38.73 | 45.07 | 42.60 | 41.19 | 35.91 | 36.61 | 36.26 |
| | | Time | 0.0 | 0.0 | 0.0 | 0.0 | 0.0 | 0.0 | 0.0 | 0.0 | 0.0 | 0.0 |
| | LDA | Accuracy | 24.56 | 24.91 | 26.08 | 25.70 | 29.22 | 26.05 | 24.29 | 26.76 | 25.70 | 29.57 |
| | | Time | 105.7 | 103.7 | 86.41 | 86.29 | 86.54 | 86.70 | 86.59 | 86.35 | 86.37 | 86.36 |
| | KNN | Accuracy | 35.43 | 31.22 | 35.91 | 28.87 | 30.98 | 34.85 | 34.15 | 37.32 | 38.38 | 33.09 |
| | | Time | 0.06 | 0.09 | 0.09 | 0.06 | 0.10 | 0.14 | 0.07 | 0.07 | 0.15 | 0.07 |
| | Dummy | Accuracy | 23.15 | 20.00 | 20.77 | 20.77 | 21.12 | 23.23 | 21.56 | 19.97 | 22.14 | 20.64 |
| | | Time | | | | | | | | | | |
| Subject F | SVM (Poly) | Accuracy | 18.75 | 25.00 | 18.75 | 23.95 | 28.12 | 29.16 | 25.00 | 33.68 | 28.42 | 25.26 |
| | | Time | 11.90 | 13.86 | 13.09 | 13.26 | 13.41 | 12.96 | 11.82 | 12.40 | 12.45 | 11.75 |
| | SVM (RBF) | Accuracy | 22.91 | 31.25 | 22.91 | 21.87 | 27.08 | 32.29 | 23.95 | 31.57 | 25.26 | 21.05 |
| | | Time | 12.38 | 12.66 | 13.29 | 13.01 | 13.31 | 12.43 | 13.55 | 13.13 | 12.93 | 11.87 |
| | Random Forest | Accuracy | 21.87 | 28.12 | 21.87 | 28.12 | 27.08 | 30.20 | 27.08 | 29.47 | 33.68 | 25.26 |
| | | Time | 0.0 | 0.0 | 0.0 | 0.0 | 0.0 | 0.0 | 0.0 | 0.0 | 0.0 | 0.0 |
| | LDA | Accuracy | 26.04 | 29.16 | 16.66 | 20.83 | 26.04 | 20.83 | 30.20 | 29.47 | 25.26 | 18.94 |
| | | Time | 4.42 | 5.23 | 5.31 | 5.25 | 5.03 | 5.49 | 4.83 | 5.03 | 5.22 | 4.49 |
| | KNN | Accuracy | 19.79 | 23.95 | 16.66 | 23.95 | 30.20 | 26.04 | 26.04 | 22.10 | 22.10 | 26.31 |
| | | Time | 0.01 | 0.03 | 0.03 | 0.03 | 0.03 | 0.03 | 0.03 | 0.01 | 0.01 | 0.03 |
| | Dummy | Accuracy | 19.79 | 26.04 | 17.70 | 21.87 | 26.04 | 26.04 | 17.70 | 20.00 | 21.05 | 21.05 |
| | | Time | | | | | | | | | | |
| Subje ct G | SVM (Poly) | Accuracy | 28.94 | 25.78 | 22.10 | 30.52 | 27.36 | 25.39 | 24.86 | 23.28 | 23.80 | 28.57 |

| | | | | | | | | | | | |
|---|---|---|---|---|---|---|---|---|---|---|---|
| | | Time | 49.35 | 45.83 | 47.97 | 45.36 | 52.89 | 52.44 | 43.55 | 45.52 | 43.65 | 46.30 |
| | SVM (RBF) | Accuracy | 29.47 | 27.36 | 28.42 | 24.21 | 27.36 | 26.98 | 29.10 | 28.57 | 29.10 | 30.68 |
| | | Time | 47.02 | 46.83 | 46.28 | 44.99 | 46.44 | 48.11 | 41.25 | 43.04 | 40.45 | 43.35 |
| | Random Forest | Accuracy | 26.31 | 34.73 | 24.73 | 26.31 | 30.52 | 26.45 | 29.62 | 25.39 | 31.21 | 26.98 |
| | | Time | 0.0 | 0.0 | 0.0 | 0.0 | 0.0 | 0.0 | 0.0 | 0.0 | 0.0 | 0.0 |
| | LDA | Accuracy | 23.15 | 22.10 | 19.47 | 27.36 | 22.10 | 22.22 | 20.63 | 19.57 | 20.10 | 17.98 |
| | | Time | 47.60 | 46.46 | 44.07 | 45.24 | 44.54 | 43.67 | 44.06 | 45.26 | 44.54 | 43.97 |
| | KNN | Accuracy | 22.10 | 29.47 | 23.15 | 29.47 | 21.57 | 30.68 | 26.45 | 23.80 | 27.51 | 21.69 |
| | | Time | 0.04 | 0.04 | 0.03 | 0.04 | 0.09 | 0.03 | 0.04 | 0.04 | 0.06 | 0.04 |
| | Dummy | Accuracy | 20.52 | 19.47 | 25.26 | 23.15 | 21.57 | 20.10 | 22.22 | 24.86 | 21.16 | 20.63 |
| | | Time | | | | | | | | | | |
| Subject H | SVM (Poly) | Accuracy | 30.43 | 27.17 | 17.39 | 19.56 | 21.73 | 18.47 | 17.39 | 15.21 | 20.64 | 28.57 |
| | | Time | 12.06 | 12.53 | 12.06 | 11.54 | 12.96 | 12.38 | 10.62 | 10.29 | 10.28 | 11.15 |
| | SVM (RBF) | Accuracy | 26.08 | 26.08 | 22.82 | 22.82 | 19.56 | 30.43 | 16.30 | 16.30 | 23.91 | 28.57 |
| | | Time | 11.57 | 11.13 | 10.93 | 10.62 | 11.33 | 11.09 | 9.37 | 9.31 | 9.99 | 9.55 |
| | Random Forest | Accuracy | 21.73 | 20.65 | 23.91 | 27.17 | 21.73 | 26.08 | 16.30 | 25.00 | 30.43 | 29.67 |
| | | Time | 0.0 | 0.0 | 0.0 | 0.0 | 0.0 | 0.0 | 0.0 | 0.0 | 0.0 | 0.0 |
| | LDA | Accuracy | 18.47 | 15.21 | 17.39 | 27.17 | 14.13 | 21.73 | 17.39 | 21.73 | 21.73 | 21.97 |
| | | Time | 4.24 | 4.61 | 4.22 | 4.97 | 4.81 | 4.31 | 3.96 | 3.99 | 3.96 | 4.03 |
| | KNN | Accuracy | 21.73 | 20.65 | 16.30 | 21.73 | 17.39 | 28.26 | 16.30 | 19.56 | 18.47 | 27.47 |
| | | Time | 0.0 | 0.02 | 0.03 | 0.01 | 0.04 | 0.01 | 0.01 | 0.03 | 0.01 | 0.01 |
| | Dummy | Accuracy | 27.17 | 16.30 | 15.21 | 22.82 | 10.86 | 18.47 | 17.39 | 21.73 | 22.82 | 26.37 |
| | | Time | | | | | | | | | | |
| Subject I | SVM (Poly) | Accuracy | 32.81 | 37.17 | 29.84 | 35.07 | 29.84 | 25.13 | 18.84 | 25.65 | 21.98 | 30.89 |
| | | Time | 46.75 | 47.37 | 50.26 | 48.49 | 46.53 | 43.83 | 46.54 | 44.27 | 44.92 | 43.81 |
| | SVM (RBF) | Accuracy | 32.29 | 31.93 | 25.65 | 34.03 | 27.74 | 31.41 | 22.51 | 32.98 | 31.41 | 26.70 |
| | | Time | 47.24 | 48.57 | 45.44 | 49.03 | 48.91 | 43.71 | 44.73 | 44.57 | 45.22 | 43.54 |
| | Random Forest | Accuracy | 34.89 | 34.55 | 35.07 | 23.03 | 35.60 | 28.79 | 27.74 | 35.05 | 27.22 | 32.46 |
| | | Time | 0.0 | 0.0 | 0.0 | 0.0 | 0.0 | 0.0 | 0.0 | 0.0 | 0.0 | 0.0 |
| | LDA | Accuracy | 19.27 | 26.70 | 21.46 | 23.56 | 20.41 | 23.56 | 24.08 | 21.98 | 23.56 | 21.46 |

| | | | | | | | | | | | |
|---|---|---|---|---|---|---|---|---|---|---|---|
| | | Time | 49.83 | 48.85 | 48.10 | 50.52 | 42.97 | 43.13 | 44.84 | 42.66 | 42.12 | 42.11 |
| | KNN | Accuracy | 28.64 | 31.93 | 26.70 | 27.22 | 28.79 | 24.60 | 30.36 | 25.13 | 26.70 | 29.31 |
| | | Time | 0.04 | 0.06 | 0.04 | 0.06 | 0.04 | 0.04 | 0.03 | 0.04 | 0.04 | 0.04 |
| | Dummy | Accuracy | 21.35 | 21.46 | 21.46 | 14.13 | 26.17 | 18.84 | 26.17 | 27.22 | 17.27 | 24.08 |
| | | Time | | | | | | | | | | |

Table 10S, 10-fold cross validation, Accuracies (%) and Training Times (Second) for each fold in Time-Frequency domain (Morlet), After PCA (100) scenario.

| | | | | | | | | | | | |
|---|---|---|---|---|---|---|---|---|---|---|---|
| Subject B | SVM (Poly) | Accuracy | 20.10 | 23.80 | 23.80 | 25.39 | 25.39 | 28.04 | 30.15 | 22.75 | 21.69 | 21.16 |
| | | Time | 0.42 | 0.41 | 0.40 | 0.40 | 0.40 | 0.40 | 0.39 | 0.39 | 0.39 | 0.40 |
| | SVM (RBF) | Accuracy | 23.80 | 24.33 | 26.98 | 21.69 | 23.28 | 26.45 | 28.57 | 21.16 | 24.33 | 22.75 |
| | | Time | 0.83 | 0.76 | 0.76 | 0.75 | 0.78 | 0.71 | 0.78 | 0.76 | 0.74 | 0.73 |
| | Random Forest | Accuracy | 20.10 | 23.28 | 22.22 | 28.04 | 26.45 | 21.16 | 26.98 | 19.57 | 25.39 | 28.57 |
| | | Time | 0.0 | 0.0 | 0.0 | 0.0 | 0.0 | 0.0 | 0.0 | 0.0 | 0.0 | 0.0 |
| | LDA | Accuracy | 21.69 | 28.57 | 29.10 | 23.80 | 25.92 | 28.04 | 28.57 | 26.98 | 28.04 | 30.15 |
| | | Time | 0.02 | 0.01 | 0.03 | 0.01 | 0.01 | 0.03 | 0.03 | 0.01 | 0.01 | 0.01 |
| | KNN | Accuracy | 24.86 | 20.10 | 25.39 | 25.39 | 28.04 | 25.92 | 28.57 | 21.16 | 24.33 | 23.80 |
| | | Time | 0.0 | 0.01 | 0.0 | 0.0 | 0.0 | 0.0 | 0.0 | 0.0 | 0.0 | 0.0 |
| | Dummy | Accuracy | 22.75 | 19.57 | 21.16 | 23.28 | 26.45 | 18.51 | 18.51 | 21.69 | 22.22 | 24.33 |
| | | Time | | | | | | | | | | |
| Subject C | SVM (Poly) | Accuracy | 25.53 | 30.85 | 41.48 | 35.10 | 26.59 | 26.59 | 28.72 | 34.04 | 28.72 | 33.33 |
| | | Time | 0.10 | 0.13 | 0.11 | 0.10 | 0.12 | 0.10 | 0.11 | 0.10 | 0.09 | 0.10 |
| | SVM (RBF) | Accuracy | 29.78 | 38.29 | 40.42 | 30.85 | 24.46 | 39.36 | 38.29 | 39.36 | 29.78 | 33.33 |
| | | Time | 0.18 | 0.23 | 0.21 | 0.23 | 0.23 | 0.20 | 0.18 | 0.20 | 0.18 | 0.20 |
| | Random Forest | Accuracy | 28.72 | 37.23 | 35.10 | 31.91 | 36.17 | 32.97 | 35.10 | 35.10 | 32.97 | 29.03 |
| | | Time | 0.0 | 0.0 | 0.0 | 0.0 | 0.0 | 0.0 | 0.0 | 0.0 | 0.0 | 0.0 |
| | LDA | Accuracy | 35.10 | 40.42 | 40.42 | 43.61 | 35.10 | 40.42 | 36.17 | 39.36 | 35.10 | 30.10 |
| | | Time | 0.01 | 0.03 | 0.01 | 0.02 | 0.01 | 0.01 | 0.01 | 0.01 | 0.02 | 0.01 |
| | KNN | Accuracy | 28.72 | 30.85 | 39.36 | 31.91 | 29.78 | 28.72 | 26.59 | 38.29 | 27.65 | 22.58 |
| | | Time | 0.0 | 0.0 | 0.0 | 0.0 | 0.0 | 0.0 | 0.0 | 0.0 | 0.0 | 0.0 |

| | | | | | | | | | | | |
|---|---|---|---|---|---|---|---|---|---|---|---|
| | Dummy | Accuracy | 25.53 | 20.21 | 23.40 | 24.46 | 22.34 | 14.89 | 20.21 | 22.34 | 19.14 | 24.73 |
| | | Time | | | | | | | | | | |
| Subject E | SVM (Poly) | Accuracy | 36.84 | 30.52 | 40.49 | 33.09 | 37.67 | 33.80 | 36.97 | 38.02 | 37.32 | 35.21 |
| | | Time | 0.94 | 0.87 | 0.87 | 0.85 | 0.89 | 0.87 | 0.87 | 0.87 | 0.87 | 0.86 |
| | SVM (RBF) | Accuracy | 41.75 | 40.00 | 45.42 | 42.60 | 41.90 | 38.73 | 42.60 | 39.08 | 41.19 | 42.25 |
| | | Time | 1.12 | 1.06 | 1.01 | 1.01 | 1.01 | 0.98 | 0.98 | 0.98 | 0.96 | 0.99 |
| | Random Forest | Accuracy | 36.14 | 36.49 | 36.61 | 33.09 | 35.56 | 36.97 | 35.21 | 31.33 | 41.90 | 38.02 |
| | | Time | 0.0 | 0.0 | 0.0 | 0.0 | 0.0 | 0.0 | 0.0 | 0.0 | 0.0 | 0.0 |
| | LDA | Accuracy | 38.94 | 39.59 | 42.60 | 41.90 | 44.36 | 44.71 | 43.66 | 35.91 | 41.90 | 41.45 |
| | | Time | 0.04 | 0.03 | 0.03 | 0.03 | 0.04 | 0.04 | 0.03 | 0.04 | 0.04 | 0.03 |
| | KNN | Accuracy | 36.14 | 29.47 | 36.26 | 31.33 | 33.45 | 35.56 | 38.38 | 35.21 | 38.38 | 35.56 |
| | | Time | 0.0 | 0.0 | 0.0 | 0.0 | 0.0 | 0.0 | 0.0 | 0.0 | 0.0 | 0.0 |
| | Dummy | Accuracy | 23.15 | 20.00 | 20.77 | 20.77 | 21.12 | 23.23 | 22.18 | 22.18 | 21.83 | 22.88 |
| | | Time | | | | | | | | | | |
| Subject F | SVM (Poly) | Accuracy | 19.79 | 26.04 | 19.79 | 22.91 | 35.41 | 29.16 | 28.12 | 26.31 | 20.00 | 24.21 |
| | | Time | 0.09 | 0.10 | 0.12 | 0.09 | 0.10 | 0.10 | 0.10 | 0.10 | 0.10 | 0.12 |
| | SVM (RBF) | Accuracy | 23.95 | 20.83 | 15.62 | 18.75 | 28.12 | 34.37 | 23.95 | 26.31 | 24.10 | 27.36 |
| | | Time | 0.20 | 0.24 | 0.21 | 0.20 | 0.26 | 0.24 | 0.17 | 0.21 | 0.18 | 0.20 |
| | Random Forest | Accuracy | 22.91 | 22.91 | 23.95 | 22.91 | 29.16 | 27.08 | 23.95 | 25.26 | 30.52 | 30.52 |
| | | Time | 0.0 | 0.0 | 0.0 | 0.0 | 0.0 | 0.0 | 0.0 | 0.0 | 0.0 | 0.0 |
| | LDA | Accuracy | 25.00 | 30.20 | 18.75 | 19.79 | 34.37 | 26.04 | 30.20 | 31.57 | 37.89 | 22.10 |
| | | Time | 0.01 | 0.01 | 0.03 | 0.03 | 0.03 | 0.01 | 0.01 | 0.01 | 0.01 | 0.01 |
| | KNN | Accuracy | 20.83 | 20.83 | 17.70 | 26.04 | 29.16 | 28.12 | 27.08 | 21.05 | 26.31 | 23.15 |
| | | Time | 0.0 | 0.0 | 0.0 | 0.0 | 0.0 | 0.0 | 0.0 | 0.0 | 0.0 | 0.0 |
| | Dummy | Accuracy | 19.79 | 26.04 | 17.70 | 21.87 | 26.04 | 26.04 | 17.70 | 20.00 | 21.05 | 21.05 |
| | | Time | | | | | | | | | | |
| Subject G | SVM (Poly) | Accuracy | 25.78 | 25.26 | 25.78 | 32.10 | 24.21 | 25.92 | 26.45 | 17.98 | 21.69 | 20.63 |
| | | Time | 0.41 | 0.41 | 0.39 | 0.42 | 0.40 | 0.43 | 0.40 | 0.40 | 0.42 | 0.39 |
| | SVM (RBF) | Accuracy | 31.57 | 28.42 | 32.63 | 28.42 | 27.36 | 31.21 | 29.62 | 20.63 | 30.15 | 29.10 |
| | | Time | 0.57 | 0.62 | 0.57 | 0.60 | 0.67 | 0.56 | 0.62 | 0.54 | 0.54 | 0.54 |

| | | | | | | | | | | | |
|---|---|---|---|---|---|---|---|---|---|---|---|
| | Random Forest | Accuracy | 27.36 | 26.31 | 24.73 | 27.89 | 25.78 | 24.33 | 30.15 | 23.80 | 26.98 | 26.45 |
| | | Time | 0.0 | 0.0 | 0.0 | 0.0 | 0.0 | 0.0 | 0.0 | 0.0 | 0.0 | 0.0 |
| | LDA | Accuracy | 31.05 | 28.94 | 29.47 | 30.00 | 30.52 | 32.80 | 30.68 | 25.92 | 37.03 | 28.57 |
| | | Time | 0.02 | 0.03 | 0.03 | 0.03 | 0.03 | 0.03 | 0.03 | 0.01 | 0.03 | 0.03 |
| | KNN | Accuracy | 24.21 | 24.73 | 27.36 | 31.05 | 21.05 | 28.04 | 26.98 | 24.33 | 22.75 | 23.28 |
| | | Time | 0.0 | 0.0 | 0.0 | 0.0 | 0.0 | 0.0 | 0.0 | 0.0 | 0.0 | 0.0 |
| | Dummy | Accuracy | 20.52 | 19.47 | 25.26 | 23.15 | 21.57 | 20.10 | 22.22 | 24.86 | 21.16 | 20.63 |
| | | Time | | | | | | | | | | |
| Subject H | SVM (Poly) | Accuracy | 33.69 | 19.56 | 21.73 | 21.73 | 16.30 | 28.26 | 25.00 | 14.13 | 21.73 | 25.27 |
| | | Time | 0.10 | 0.12 | 0.10 | 0.10 | 0.11 | 0.11 | 0.10 | 0.10 | 0.10 | 0.10 |
| | SVM (RBF) | Accuracy | 30.43 | 21.73 | 19.56 | 29.34 | 20.65 | 26.08 | 18.47 | 20.65 | 20.65 | 34.06 |
| | | Time | 0.15 | 0.12 | 0.12 | 0.12 | 0.12 | 0.12 | 0.13 | 0.11 | 0.12 | 0.12 |
| | Random Forest | Accuracy | 28.26 | 26.08 | 15.21 | 27.17 | 7.60 | 27.17 | 11.95 | 18.47 | 17.39 | 30.76 |
| | | Time | 0.0 | 0.0 | 0.0 | 0.0 | 0.0 | 0.0 | 0.0 | 0.0 | 0.0 | 0.0 |
| | LDA | Accuracy | 29.34 | 18.47 | 21.73 | 31.52 | 14.13 | 26.08 | 17.39 | 16.30 | 21.73 | 23.07 |
| | | Time | 0.02 | 0.01 | 0.01 | 0.01 | 0.02 | 0.01 | 0.01 | 0.01 | 0.02 | 0.01 |
| | KNN | Accuracy | 17.39 | 20.65 | 19.56 | 17.39 | 16.30 | 25.00 | 17.39 | 20.65 | 18.47 | 24.17 |
| | | Time | 0.0 | 0.01 | 0.0 | 0.0 | 0.0 | 0.0 | 0.0 | 0.0 | 0.0 | 0.0 |
| | Dummy | Accuracy | 27.17 | 16.30 | 15.21 | 22.82 | 10.86 | 18.47 | 17.39 | 21.73 | 22.82 | 26.37 |
| | | Time | | | | | | | | | | |
| Subject I | SVM (Poly) | Accuracy | 26.56 | 30.89 | 35.07 | 29.31 | 26.70 | 26.17 | 21.46 | 25.65 | 31.93 | 24.60 |
| | | Time | 0.40 | 0.40 | 0.40 | 0.40 | 0.40 | 0.40 | 0.42 | 0.42 | 0.40 | 0.40 |
| | SVM (RBF) | Accuracy | 30.72 | 35.60 | 29.84 | 31.93 | 28.79 | 30.89 | 21.98 | 28.79 | 31.93 | 31.41 |
| | | Time | 0.68 | 0.60 | 0.62 | 0.67 | 0.62 | 0.64 | 0.62 | 0.60 | 0.64 | 0.64 |
| | Random Forest | Accuracy | 29.68 | 28.79 | 25.65 | 25.65 | 25.13 | 21.98 | 29.31 | 32.46 | 22.51 | 30.36 |
| | | Time | 0.0 | 0.0 | 0.0 | 0.0 | 0.0 | 0.0 | 0.0 | 0.0 | 0.0 | 0.0 |
| | LDA | Accuracy | 30.72 | 34.55 | 30.36 | 33.50 | 27.74 | 26.17 | 29.84 | 28.79 | 27.74 | 26.70 |
| | | Time | 0.03 | 0.03 | 0.03 | 0.01 | 0.03 | 0.01 | 0.01 | 0.03 | 0.01 | 0.01 |
| | KNN | Accuracy | 28.64 | 29.84 | 32.98 | 30.36 | 27.74 | 24.60 | 29.84 | 30.36 | 26.17 | 30.89 |
| | | Time | 0.0 | 0.0 | 0.0 | 0.0 | 0.0 | 0.0 | 0.0 | 0.0 | 0.0 | 0.0 |

| | | | | | | | | | | | |
|---|---|---|---|---|---|---|---|---|---|---|---|
| | Dummy | Accuracy | 21.35 | 21.46 | 21.46 | 14.13 | 26.17 | 18.84 | 26.17 | 27.22 | 17.27 | 24.08 |
| | | Time | | | | | | | | | | |
| Independent | SVM (Poly) | Accuracy | 26.40 | 24.91 | 22.88 | 23.52 | 25.63 | 22.29 | 22.37 | 24.14 | 23.61 | 24.49 |
| | | Time | 81.88 | 86.84 | 85.60 | 91.00 | 89.89 | 86.09 | 90.86 | 86.28 | 91.27 | 83.56 |
| | SVM (RBF) | Accuracy | 29.75 | 29.13 | 28.25 | 27.92 | 30.30 | 29.77 | 31.10 | 28.45 | 27.75 | 27.57 |
| | | Time | 22.42 | 20.00 | 20.90 | 20.64 | 19.64 | 22.16 | 19.78 | 20.73 | 20.60 | 19.99 |
| | Random Forest | Accuracy | 28.78 | 27.28 | 25.61 | 24.84 | 29.33 | 26.96 | 28.10 | 28.19 | 25.90 | 27.40 |
| | | Time | 0.0 | 0.0 | 0.0 | 0.0 | 0.0 | 0.0 | 0.0 | 0.0 | 0.0 | 0.0 |
| | LDA | Accuracy | 30.01 | 29.75 | 28.43 | 28.72 | 30.04 | 28.19 | 30.04 | 28.98 | 29.16 | 27.57 |
| | | Time | 0.15 | 0.15 | 0.14 | 0.15 | 0.15 | 0.14 | 0.14 | 0.14 | 0.15 | 0.14 |
| | KNN | Accuracy | 27.64 | 28.69 | 24.47 | 25.81 | 27.31 | 25.28 | 26.07 | 25.81 | 23.78 | 26.16 |
| | | Time | 0.0 | 0.0 | 0.0 | 0.0 | 0.0 | 0.0 | 0.0 | 0.0 | 0.0 | 0.0 |
| | Dummy | Accuracy | 21.03 | 22.53 | 20.51 | 22.02 | 23.08 | 21.76 | 20.96 | 21.05 | 22.81 | 22.55 |
| | | Time | | | | | | | | | | |

Table 11S, 10-fold cross validation, Accuracies (%) and Training Times (Second) for each fold in Time-Frequency domain (Wigner), Before PCA scenario.

| | | | | | | | | | | | |
|---|---|---|---|---|---|---|---|---|---|---|---|
| Subject B | SVM (Poly) | Accuracy | 31.74 | 32.27 | 29.62 | 31.74 | 37.03 | 31.21 | 39.68 | 36.50 | 31.21 | 32.27 |
| | | Time | 235.6 | 212.9 | 212.7 | 215.8 | 214.9 | 213.5 | 212.9 | 216.9 | 211.6 | 211.6 |
| | SVM (RBF) | Accuracy | 35.44 | 34.39 | 31.74 | 36.50 | 34.92 | 29.10 | 38.09 | 38.09 | 28.57 | 34.39 |
| | | Time | 234.4 | 241.5 | 221.2 | 226.2 | 218.6 | 218.2 | 217.6 | 220.5 | 217.6 | 220.7 |
| | Random Forest | Accuracy | 37.56 | 38.62 | 30.68 | 34.92 | 35.97 | 27.51 | 28.57 | 32.80 | 29.62 | 31.74 |
| | | Time | 0.0 | 0.0 | 0.0 | 0.0 | 0.0 | 0.0 | 0.0 | 0.0 | 0.0 | 0.01 |
| | LDA | Accuracy | 27.51 | 28.57 | 27.51 | 33.33 | 31.74 | 28.04 | 32.80 | 30.68 | 26.45 | 22.22 |
| | | Time | 78.08 | 62.69 | 61.39 | 62.90 | 59.07 | 65.50 | 60.32 | 63.54 | 62.11 | 68.21 |
| | KNN | Accuracy | 23.80 | 26.98 | 25.92 | 21.69 | 26.98 | 18.51 | 21.16 | 24.86 | 16.93 | 22.75 |
| | | Time | 0.96 | 0.34 | 0.37 | 0.32 | 0.25 | 0.31 | 0.31 | 1.00 | 0.32 | 0.32 |
| | Dummy | Accuracy | 22.75 | 19.57 | 21.16 | 23.28 | 26.45 | 18.51 | 18.51 | 21.69 | 22.22 | 24.33 |
| | | Time | | | | | | | | | | |

| | | | | | | | | | | | |
|---|---|---|---|---|---|---|---|---|---|---|---|
| Subject C | SVM (Poly) | Accuracy | 44.68 | 58.51 | 43.61 | 53.19 | 47.87 | 43.61 | 41.48 | 51.06 | 37.23 | 43.01 |
| | | Time | 57.48 | 61.66 | 59.26 | 62.81 | 52.65 | 53.27 | 53.08 | 53.82 | 51.60 | 52.95 |
| | SVM (RBF) | Accuracy | 47.87 | 55.31 | 47.87 | 54.25 | 45.74 | 46.80 | 39.36 | 52.12 | 41.48 | 46.23 |
| | | Time | 60.75 | 59.17 | 60.67 | 57.10 | 60.72 | 53.43 | 53.73 | 54.01 | 53.14 | 54.42 |
| | Random Forest | Accuracy | 37.23 | 43.61 | 37.23 | 41.48 | 38.29 | 32.97 | 41.48 | 47.87 | 36.17 | 38.70 |
| | | Time | 0.0 | 0.0 | 0.0 | 0.0 | 0.0 | 0.0 | 0.0 | 0.0 | 0.0 | 0.0 |
| | LDA | Accuracy | 44.68 | 45.74 | 43.61 | 35.10 | 39.36 | 40.42 | 32.97 | 45.74 | 30.85 | 48.38 |
| | | Time | 27.09 | 26.14 | 27.99 | 35.49 | 23.18 | 24.13 | 25.74 | 32.23 | 36.15 | 25.97 |
| | KNN | Accuracy | 23.40 | 29.78 | 23.40 | 34.04 | 21.27 | 22.34 | 29.78 | 18.08 | 23.40 | 34.40 |
| | | Time | 0.15 | 0.12 | 0.17 | 0.15 | 0.10 | 0.17 | 0.20 | 0.17 | 0.15 | 0.12 |
| | Dummy | Accuracy | 25.53 | 20.21 | 23.40 | 24.46 | 22.34 | 14.89 | 20.21 | 22.34 | 19.14 | 24.73 |
| | | Time | | | | | | | | | | |
| Subject E | SVM (Poly) | Accuracy | 39.64 | 38.59 | 47.88 | 40.84 | 48.59 | 46.47 | 44.71 | 42.60 | 47.88 | 41.90 |
| | | Time | 433.0 | 420.7 | 421.1 | 419.9 | 462.0 | 457.0 | 438.4 | 425.9 | 465.1 | 475.6 |
| | SVM (RBF) | Accuracy | 42.45 | 40.35 | 47.88 | 42.95 | 50.35 | 45.77 | 45.42 | 43.30 | 48.94 | 44.01 |
| | | Time | 448.2 | 448.3 | 442.6 | 440.9 | 478.2 | 453.8 | 452.3 | 454.7 | 457.2 | 451.7 |
| | Random Forest | Accuracy | 35.08 | 40.35 | 41.90 | 37.67 | 48.94 | 44.01 | 42.95 | 39.08 | 38.02 | 38.02 |
| | | Time | 0.0 | 0.0 | 0.0 | 0.0 | 0.0 | 0.0 | 0.0 | 0.0 | 0.0 | 0.0 |
| | LDA | Accuracy | 37.89 | 30.17 | 42.60 | 35.56 | 41.19 | 39.43 | 41.90 | 37.67 | 37.67 | 36.97 |
| | | Time | 147.1 | 143.0 | 144.5 | 134.8 | 279.2 | 250.8 | 152.6 | 177.2 | 120.9 | 141.2 |
| | KNN | Accuracy | 31.57 | 27.01 | 32.39 | 31.33 | 30.28 | 23.94 | 30.28 | 27.46 | 28.16 | 28.52 |
| | | Time | 13.9 | 11.6 | 22.7 | 1.99 | 9.69 | 7.08 | 63.03 | 2.99 | 3.81 | 8.36 |
| | Dummy | Accuracy | 23.15 | 20.00 | 20.77 | 20.77 | 21.12 | 23.23 | 22.18 | 22.18 | 21.83 | 22.88 |
| | | Time | | | | | | | | | | |
| Subject F | SVM (Poly) | Accuracy | 28.12 | 39.58 | 27.08 | 34.37 | 37.50 | 37.50 | 34.37 | 24.21 | 29.47 | 32.63 |
| | | Time | 56.63 | 55.77 | 55.28 | 55.71 | 59.52 | 60.81 | 65.53 | 63.79 | 61.98 | 62.28 |
| | SVM (RBF) | Accuracy | 25.00 | 39.58 | 25.00 | 37.50 | 38.54 | 32.29 | 34.37 | 27.36 | 32.63 | 28.42 |
| | | Time | 60.39 | 55.53 | 56.55 | 56.60 | 57.59 | 59.95 | 61.08 | 60.94 | 67.99 | 62.52 |
| | Random Forest | Accuracy | 25.00 | 38.54 | 27.08 | 35.41 | 34.37 | 37.50 | 31.25 | 29.47 | 27.36 | 26.31 |
| | | Time | 0.0 | 0.0 | 0.0 | 0.0 | 0.0 | 0.0 | 0.0 | 0.0 | 0.0 | 0.0 |

| | | | | | | | | | | | |
|---|---|---|---|---|---|---|---|---|---|---|---|
| | LDA | Accuracy | 28.12 | 25.00 | 30.20 | 27.08 | 35.41 | 32.29 | 27.08 | 21.05 | 26.31 | 31.57 |
| | | Time | 24.78 | 24.17 | 24.61 | 24.65 | 25.41 | 25.80 | 26.11 | 26.23 | 25.48 | 27.13 |
| | KNN | Accuracy | 15.62 | 26.04 | 16.66 | 26.04 | 26.04 | 27.08 | 20.83 | 16.84 | 20.00 | 20.00 |
| | | Time | 0.11 | 0.13 | 0.19 | 0.12 | 0.13 | 0.23 | 0.18 | 0.21 | 0.15 | 0.20 |
| | Dummy | Accuracy | 19.79 | 26.04 | 17.70 | 21.87 | 26.04 | 26.04 | 17.70 | 20.00 | 21.05 | 21.05 |
| | | Time | | | | | | | | | | |
| Subject G | SVM (Poly) | Accuracy | 34.21 | 24.21 | 27.89 | 29.47 | 28.42 | 25.39 | 27.51 | 28.57 | 32.27 | 29.10 |
| | | Time | 208.3 | 205.9 | 220.3 | 221.0 | 208.1 | 219.8 | 203.5 | 221.6 | 220.8 | 209.4 |
| | SVM (RBF) | Accuracy | 33.68 | 27.36 | 28.94 | 27.36 | 24.21 | 26.45 | 28.04 | 30.15 | 31.21 | 28.04 |
| | | Time | 208.6 | 212.2 | 210.2 | 221.2 | 213.7 | 214.4 | 216.3 | 219.6 | 217.3 | 212.3 |
| | Random Forest | Accuracy | 33.15 | 26.31 | 37.36 | 33.68 | 24.73 | 28.57 | 30.15 | 27.51 | 29.10 | 29.62 |
| | | Time | 0.0 | 0.0 | 0.03 | 0.01 | 0.0 | 0.0 | 0.0 | 0.0 | 0.0 | 0.0 |
| | LDA | Accuracy | 27.89 | 31.05 | 25.26 | 27.89 | 25.26 | 19.04 | 26.98 | 26.45 | 26.98 | 30.15 |
| | | Time | 67.22 | 85.25 | 99.76 | 68.24 | 69.53 | 63.89 | 70.22 | 64.26 | 67.21 | 60.00 |
| | KNN | Accuracy | 23.15 | 25.26 | 25.26 | 21.05 | 24.21 | 23.28 | 24.33 | 24.86 | 27.51 | 25.39 |
| | | Time | 0.85 | 1.03 | 1.14 | 0.57 | 0.89 | 0.29 | 0.36 | 0.47 | 0.44 | 0.40 |
| | Dummy | Accuracy | 20.52 | 19.47 | 25.26 | 23.15 | 21.57 | 20.10 | 22.22 | 24.86 | 21.16 | 20.63 |
| | | Time | | | | | | | | | | |
| Subject H | SVM (Poly) | Accuracy | 23.91 | 23.91 | 20.65 | 23.91 | 29.34 | 22.82 | 26.08 | 21.73 | 25.00 | 19.78 |
| | | Time | 55.28 | 56.02 | 54.17 | 57.49 | 63.21 | 64.82 | 65.68 | 55.99 | 65.54 | 65.79 |
| | SVM (RBF) | Accuracy | 19.56 | 20.65 | 20.65 | 20.65 | 28.26 | 23.91 | 26.08 | 18.47 | 25.00 | 20.87 |
| | | Time | 58.91 | 55.91 | 58.11 | 54.17 | 63.43 | 61.89 | 64.90 | 54.68 | 62.33 | 61.91 |
| | Random Forest | Accuracy | 23.91 | 20.65 | 13.04 | 18.47 | 17.39 | 20.65 | 17.39 | 28.26 | 22.82 | 25.27 |
| | | Time | 0.0 | 0.0 | 0.0 | 0.0 | 0.0 | 0.0 | 0.0 | 0.0 | 0.0 | 0.0 |
| | LDA | Accuracy | 20.65 | 21.73 | 27.17 | 25.00 | 18.47 | 20.65 | 23.91 | 26.08 | 27.17 | 27.47 |
| | | Time | 24.63 | 24.13 | 24.37 | 25.14 | 26.24 | 26.61 | 26.09 | 25.01 | 24.17 | 27.76 |
| | KNN | Accuracy | 14.13 | 16.30 | 22.82 | 18.47 | 23.91 | 16.30 | 15.21 | 21.73 | 17.39 | 16.48 |
| | | Time | 0.10 | 0.14 | 0.14 | 0.10 | 0.15 | 0.18 | 0.15 | 0.14 | 0.11 | 0.18 |
| | Dummy | Accuracy | 27.17 | 16.30 | 15.21 | 22.82 | 10.86 | 18.47 | 17.39 | 21.73 | 22.82 | 26.37 |
| | | Time | | | | | | | | | | |

| | | | | | | | | | | | | |
|---|---|---|---|---|---|---|---|---|---|---|---|---|
| Subject I | SVM (Poly) | Accuracy | 25.00 | 29.31 | 31.41 | 28.27 | 29.84 | 36.64 | 35.07 | 23.03 | 25.65 | 25.00 |
| | | Time | 241.5 | 216.3 | 212.2 | 214.1 | 212.5 | 263.4 | 271.4 | 229.3 | 231.2 | 231.4 |
| | SVM (RBF) | Accuracy | 28.64 | 31.93 | 32.46 | 26.17 | 30.36 | 29.31 | 32.46 | 36.64 | 26.17 | 26.17 |
| | | Time | 244.1 | 230.0 | 210.9 | 210.6 | 211.6 | 266.3 | 233.1 | 230.9 | 231.5 | 233.4 |
| | Random Forest | Accuracy | 30.20 | 31.41 | 34.03 | 29.31 | 39.26 | 36.12 | 27.74 | 31.93 | 26.70 | 27.74 |
| | | Time | 0.0 | 0.0 | 0.0 | 0.0 | 0.0 | 0.0 | 0.0 | 0.0 | 0.0 | 0.0 |
| | LDA | Accuracy | 26.56 | 33.50 | 30.89 | 26.70 | 33.50 | 28.79 | 27.74 | 31.41 | 26.70 | 29.84 |
| | | Time | 78.60 | 65.74 | 77.85 | 63.56 | 64.32 | 73.59 | 72.61 | 62.23 | 72.27 | 66.42 |
| | KNN | Accuracy | 28.64 | 27.22 | 23.03 | 20.41 | 28.79 | 30.36 | 34.03 | 31.41 | 20.41 | 23.56 |
| | | Time | 0.28 | 0.82 | 0.53 | 0.39 | 0.35 | 0.48 | 0.43 | 0.35 | 0.35 | 0.31 |
| | Dummy | Accuracy | 21.35 | 21.46 | 21.46 | 14.13 | 26.17 | 18.84 | 26.17 | 27.22 | 17.27 | 24.08 |
| | | Time | | | | | | | | | | |

Table 12S, 10-fold cross validation, Accuracies (%) and Training Times (Second) for each fold in Time-Frequency domain (Wigner), After PCA (150) scenario.

| | | | | | | | | | | | | |
|---|---|---|---|---|---|---|---|---|---|---|---|---|
| Subject B | SVM (Poly) | Accuracy | 32.80 | 28.57 | 22.22 | 26.45 | 35.97 | 26.98 | 28.57 | 26.45 | 26.45 | 29.10 |
| | | Time | 0.51 | 0.49 | 0.50 | 0.50 | 0.48 | 0.50 | 0.49 | 0.48 | 0.49 | 0.50 |
| | SVM (RBF) | Accuracy | 37.56 | 31.74 | 26.98 | 36.50 | 38.62 | 33.86 | 39.15 | 27.51 | 30.68 | 35.44 |
| | | Time | 0.62 | 0.65 | 0.71 | 0.66 | 0.63 | 0.63 | 0.67 | 0.74 | 0.63 | 0.66 |
| | Random Forest | Accuracy | 32.27 | 29.10 | 25.39 | 30.68 | 31.21 | 26.45 | 26.45 | 31.21 | 23.28 | 28.04 |
| | | Time | 0.0 | 0.0 | 0.0 | 0.0 | 0.0 | 0.0 | 0.0 | 0.0 | 0.0 | 0.0 |
| | LDA | Accuracy | 37.56 | 35.97 | 33.33 | 35.97 | 40.74 | 30.68 | 42.32 | 31.21 | 30.68 | 30.15 |
| | | Time | 0.05 | 0.05 | 0.03 | 0.02 | 0.03 | 0.04 | 0.04 | 0.04 | 0.04 | 0.03 |
| | KNN | Accuracy | 29.62 | 27.51 | 22.22 | 22.75 | 29.10 | 22.75 | 24.86 | 27.51 | 23.80 | 19.57 |
| | | Time | 0.0 | 0.0 | 0.0 | 0.0 | 0.0 | 0.0 | 0.0 | 0.0 | 0.0 | 0.0 |
| | Dummy | Accuracy | 22.75 | 19.57 | 21.16 | 23.28 | 26.45 | 18.51 | 18.51 | 21.69 | 22.22 | 24.33 |
| | | Time | | | | | | | | | | |
| Subject C | SVM (Poly) | Accuracy | 28.72 | 39.36 | 30.85 | 41.48 | 27.65 | 30.85 | 32.97 | 47.87 | 40.42 | 40.86 |
| | | Time | 0.12 | 0.12 | 0.12 | 0.12 | 0.14 | 0.12 | 0.12 | 0.12 | 0.12 | 0.12 |
| | SVM (RBF) | Accuracy | 39.36 | 47.87 | 42.55 | 34.04 | 39.36 | 39.36 | 40.42 | 40.42 | 42.55 | 39.78 |

| | | | | | | | | | | | | |
|---|---|---|---|---|---|---|---|---|---|---|---|---|
| | | Time | 0.17 | 0.17 | 0.18 | 0.20 | 0.20 | 0.21 | 0.20 | 0.20 | 0.20 | 0.20 |
| | Random Forest | Accuracy | 25.53 | 38.29 | 27.65 | 42.55 | 31.91 | 25.53 | 34.04 | 36.17 | 30.85 | 31.18 |
| | | Time | 0.0 | 0.0 | 0.0 | 0.0 | 0.0 | 0.0 | 0.0 | 0.0 | 0.0 | 0.0 |
| | LDA | Accuracy | 45.74 | 57.44 | 44.68 | 39.36 | 48.93 | 45.74 | 42.55 | 50.00 | 38.29 | 43.01 |
| | | Time | 0.03 | 0.03 | 0.03 | 0.04 | 0.03 | 0.03 | 0.03 | 0.03 | 0.03 | 0.03 |
| | KNN | Accuracy | 24.46 | 24.46 | 26.59 | 36.17 | 27.65 | 21.27 | 36.17 | 31.91 | 25.53 | 32.25 |
| | | Time | 0.0 | 0.0 | 0.0 | 0.0 | 0.0 | 0.0 | 0.0 | 0.0 | 0.0 | 0.0 |
| | Dummy | Accuracy | 25.53 | 20.21 | 23.40 | 24.46 | 22.34 | 14.89 | 20.21 | 22.34 | 19.14 | 24.73 |
| | | Time | | | | | | | | | | |
| Subject E | SVM (Poly) | Accuracy | 37.89 | 36.14 | 47.18 | 43.30 | 42.95 | 39.43 | 39.78 | 36.26 | 39.78 | 39.78 |
| | | Time | 1.12 | 1.14 | 1.23 | 1.17 | 1.29 | 1.40 | 1.49 | 1.43 | 1.41 | 1.40 |
| | SVM (RBF) | Accuracy | 34.73 | 42.10 | 50.00 | 38.38 | 42.25 | 48.59 | 39.08 | 40.14 | 45.42 | 39.78 |
| | | Time | 1.02 | 0.98 | 0.99 | 1.01 | 1.03 | 1.23 | 1.15 | 1.26 | 1.24 | 1.21 |
| | Random Forest | Accuracy | 36.84 | 34.73 | 40.49 | 34.85 | 35.91 | 40.49 | 38.02 | 32.39 | 36.61 | 32.39 |
| | | Time | 0.0 | 0.0 | 0.0 | 0.0 | 0.0 | 0.0 | 0.0 | 0.0 | 0.0 | 0.0 |
| | LDA | Accuracy | 38.94 | 39.29 | 47.18 | 40.14 | 47.88 | 45.77 | 44.71 | 44.71 | 42.60 | 40.14 |
| | | Time | 0.07 | 0.08 | 0.07 | 0.07 | 0.07 | 0.07 | 0.07 | 0.09 | 0.09 | 0.07 |
| | KNN | Accuracy | 28.07 | 30.52 | 36.61 | 33.80 | 30.98 | 33.09 | 33.80 | 35.56 | 33.09 | 35.91 |
| | | Time | 0.0 | 0.0 | 0.0 | 0.0 | 0.0 | 0.0 | 0.0 | 0.0 | 0.0 | 0.0 |
| | Dummy | Accuracy | 23.15 | 20.00 | 20.77 | 20.77 | 21.12 | 23.23 | 22.18 | 22.18 | 21.83 | 22.88 |
| | | Time | | | | | | | | | | |
| Subject F | SVM (Poly) | Accuracy | 20.83 | 28.12 | 22.91 | 34.37 | 29.16 | 21.87 | 26.04 | 24.21 | 18.94 | 30.52 |
| | | Time | 0.12 | 0.12 | 0.12 | 0.14 | 0.12 | 0.12 | 0.14 | 0.11 | 0.14 | 0.14 |
| | SVM (RBF) | Accuracy | 29.16 | 36.45 | 26.04 | 30.20 | 23.95 | 23.95 | 32.29 | 34.73 | 21.05 | 31.57 |
| | | Time | 0.20 | 0.23 | 0.20 | 0.24 | 0.20 | 0.25 | 0.24 | 0.23 | 0.21 | 0.23 |
| | Random Forest | Accuracy | 23.95 | 31.25 | 18.75 | 26.04 | 33.33 | 34.37 | 25.00 | 25.26 | 28.42 | 26.31 |
| | | Time | 0.0 | 0.0 | 0.0 | 0.0 | 0.0 | 0.0 | 0.0 | 0.0 | 0.0 | 0.0 |
| | LDA | Accuracy | 36.45 | 33.33 | 25.00 | 23.95 | 40.62 | 33.33 | 25.00 | 33.68 | 26.31 | 29.47 |
| | | Time | 0.03 | 0.03 | 0.02 | 0.03 | 0.01 | 0.01 | 0.03 | 0.03 | 0.03 | 0.03 |
| | KNN | Accuracy | 23.95 | 32.29 | 19.79 | 22.91 | 29.16 | 26.04 | 20.83 | 23.15 | 18.94 | 26.31 |

| | | | | | | | | | | | |
|---|---|---|---|---|---|---|---|---|---|---|---|
| | | Time | 0.0 | 0.0 | 0.0 | 0.0 | 0.0 | 0.0 | 0.0 | 0.0 | 0.0 | 0.0 |
| | Dummy | Accuracy | 19.79 | 26.04 | 17.70 | 21.87 | 26.04 | 26.04 | 17.70 | 20.00 | 21.05 | 21.05 |
| | | Time | | | | | | | | | | |
| Subject G | SVM (Poly) | Accuracy | 30.52 | 26.84 | 30.00 | 26.31 | 24.21 | 29.10 | 26.98 | 22.75 | 27.51 | 22.22 |
| | | Time | 0.53 | 0.53 | 0.53 | 0.53 | 0.51 | 0.64 | 0.65 | 0.65 | 0.68 | 0.67 |
| | SVM (RBF) | Accuracy | 35.26 | 31.57 | 27.89 | 33.68 | 23.68 | 24.33 | 33.33 | 30.68 | 30.15 | 29.62 |
| | | Time | 0.57 | 0.59 | 0.54 | 0.57 | 0.71 | 0.76 | 0.71 | 0.74 | 0.73 | 0.76 |
| | Random Forest | Accuracy | 27.36 | 22.10 | 27.89 | 29.45 | 26.31 | 23.28 | 25.39 | 32.80 | 29.62 | 27.51 |
| | | Time | 0.0 | 0.0 | 0.0 | 0.0 | 0.0 | 0.0 | 0.0 | 0.0 | 0.0 | 0.0 |
| | LDA | Accuracy | 38.42 | 25.26 | 30.00 | 33.68 | 30.00 | 30.68 | 35.97 | 26.98 | 30.68 | 34.92 |
| | | Time | 0.06 | 0.05 | 0.06 | 0.06 | 0.05 | 0.06 | 0.06 | 0.06 | 0.06 | 0.06 |
| | KNN | Accuracy | 24.21 | 27.89 | 28.42 | 30.52 | 21.05 | 21.16 | 23.80 | 28.04 | 25.92 | 20.63 |
| | | Time | 0.0 | 0.0 | 0.0 | 0.0 | 0.0 | 0.0 | 0.0 | 0.0 | 0.0 | 0.0 |
| | Dummy | Accuracy | 20.52 | 19.47 | 25.26 | 23.15 | 21.57 | 20.10 | 22.22 | 24.86 | 21.16 | 20.63 |
| | | Time | | | | | | | | | | |
| Subject H | SVM (Poly) | Accuracy | 22.82 | 16.30 | 21.73 | 27.17 | 10.86 | 20.65 | 21.73 | 18.47 | 21.73 | 26.37 |
| | | Time | 0.20 | 0.24 | 0.20 | 0.20 | 0.18 | 0.20 | 0.24 | 0.20 | 0.17 | 0.22 |
| | SVM (RBF) | Accuracy | 19.56 | 17.39 | 16.30 | 18.47 | 23.91 | 14.13 | 16.30 | 16.30 | 26.08 | 16.48 |
| | | Time | 0.17 | 0.18 | 0.15 | 0.20 | 0.14 | 0.17 | 0.17 | 0.14 | 0.18 | 0.15 |
| | Random Forest | Accuracy | 16.30 | 19.56 | 18.47 | 16.30 | 16.30 | 20.65 | 18.47 | 17.39 | 16.30 | 20.87 |
| | | Time | 0.0 | 0.0 | 0.0 | 0.0 | 0.0 | 0.0 | 0.0 | 0.0 | 0.0 | 0.0 |
| | LDA | Accuracy | 16.30 | 26.08 | 15.21 | 20.65 | 20.65 | 19.56 | 11.95 | 17.39 | 22.82 | 23.07 |
| | | Time | 0.02 | 0.02 | 0.03 | 0.03 | 0.01 | 0.01 | 0.03 | 0.03 | 0.03 | 0.03 |
| | KNN | Accuracy | 17.39 | 14.13 | 22.82 | 21.73 | 20.65 | 19.56 | 13.04 | 23.91 | 17.39 | 15.38 |
| | | Time | 0.0 | 0.0 | 0.0 | 0.0 | 0.0 | 0.0 | 0.0 | 0.0 | 0.0 | 0.0 |
| | Dummy | Accuracy | 27.17 | 16.30 | 15.21 | 22.82 | 10.86 | 18.47 | 17.39 | 21.73 | 22.82 | 26.37 |
| | | Time | | | | | | | | | | |
| Subject I | SVM (Poly) | Accuracy | 30.20 | 31.41 | 28.27 | 34.03 | 30.89 | 31.41 | 24.08 | 31.93 | 21.46 | 20.94 |
| | | Time | 0.86 | 0.76 | 0.68 | 0.68 | 0.66 | 0.67 | 0.75 | 0.57 | 0.72 | 0.62 |
| | SVM (RBF) | Accuracy | 32.81 | 31.41 | 35.60 | 34.03 | 32.46 | 30.36 | 35.60 | 30.36 | 31.41 | 25.13 |

| | | | | | | | | | | | | |
|---|---|---|---|---|---|---|---|---|---|---|---|---|
| | | Time | 0.48 | 0.54 | 0.56 | 0.48 | 0.51 | 0.50 | 0.54 | 0.53 | 0.52 | 0.56 |
| | Random Forest | Accuracy | 28.64 | 28.27 | 31.93 | 29.31 | 31.93 | 27.22 | 31.41 | 30.36 | 26.70 | 26.70 |
| | | Time | 0.0 | 0.0 | 0.0 | 0.0 | 0.0 | 0.0 | 0.0 | 0.0 | 0.0 | 0.0 |
| | LDA | Accuracy | 29.68 | 35.07 | 41.88 | 34.55 | 31.93 | 32.46 | 34.03 | 28.27 | 27.74 | 26.17 |
| | | Time | 0.06 | 0.04 | 0.05 | 0.06 | 0.05 | 0.04 | 0.04 | 0.06 | 0.04 | 0.06 |
| | KNN | Accuracy | 27.08 | 27.22 | 27.74 | 25.13 | 21.98 | 23.56 | 26.17 | 26.70 | 23.03 | 23.03 |
| | | Time | 0.0 | 0.0 | 0.0 | 0.0 | 0.0 | 0.0 | 0.0 | 0.0 | 0.0 | 0.0 |
| | Dummy | Accuracy | 21.35 | 21.46 | 21.46 | 14.13 | 26.17 | 18.84 | 26.17 | 27.22 | 17.27 | 24.08 |
| | | Time | | | | | | | | | | |
| Independent | SVM (Poly) | Accuracy | 29.75 | 25.80 | 29.66 | 25.19 | 29.16 | 26.34 | 28.01 | 27.84 | 29.77 | 24.14 |
| | | Time | 61.92 | 78.16 | 65.91 | 68.25 | 60.59 | 64.16 | 59.50 | 59.51 | 60.86 | 65.83 |
| | SVM (RBF) | Accuracy | 32.06 | 31.16 | 29.48 | 27.84 | 31.66 | 28.89 | 32.24 | 29.42 | 30.57 | 29.86 |
| | | Time | 27.13 | 26.08 | 31.96 | 30.67 | 30.43 | 26.91 | 26.19 | 25.89 | 29.26 | 25.61 |
| | Random Forest | Accuracy | 26.76 | 26.93 | 26.84 | 27.75 | 29.25 | 26.78 | 26.51 | 27.04 | 29.25 | 27.48 |
| | | Time | 0.06 | 0.15 | 0.01 | 0.01 | 0.01 | 0.01 | 0.0 | 0.01 | 0.01 | 0.0 |
| | LDA | Accuracy | 33.09 | 33.18 | 29.75 | 27.92 | 31.62 | 27.66 | 31.80 | 28.81 | 28.98 | 30.48 |
| | | Time | 0.35 | 0.32 | 0.31 | 0.34 | 0.31 | 0.35 | 0.30 | 0.32 | 0.32 | 0.31 |
| | KNN | Accuracy | 26.23 | 25.26 | 26.49 | 26.34 | 28.72 | 25.63 | 26.87 | 25.19 | 27.48 | 23.17 |
| | | Time | 0.0 | 0.0 | 0.01 | 0.0 | 0.0 | 0.01 | 0.0 | 0.0 | 0.0 | 0.0 |
| | Dummy | Accuracy | 21.03 | 22.53 | 20.51 | 22.02 | 23.08 | 21.76 | 20.96 | 21.05 | 22.81 | 22.55 |
| | | Time | | | | | | | | | | |

Table 13S, Comparing Time domain (before PCA) and Wigner (before PCA) accuracies. P_Values calculated with one-way ANOVA test. Accuracies are mean accuracies of 10-fold cross validation ± var

| | | Accuracy (time) | Accuracy (time-freq) | statistic | P-Value |
|---|---|---|---|---|---|
| Subject B | Poly | 41.39 ±6.85 | 33.32 ±9.41 | 35.33 | 0.000 *** |
| | RBF | 46.50 ±15.48 | 33.82 ±14.27 | 48.62 | 0.001 ** |
| | Random Forest | 37.98 ±11.36 | 32.79 ±13.26 | 9.82 | 0.005 ** |
| | LDA | 36.44 ±14.14 | 28.88 ±10.14 | 21.20 | 0.000 *** |
| | KNN | 35.97 ±6.49 | 22.95 ±10.64 | 88.96 | 0.000 *** |

| | | | | | | |
|---|---|---|---|---|---|---|
| Subject C | | Poly | 55.89 ±14.47 | 46.42 ±35.49 | 16.14 | 0.000 *** |
| | | RBF | 58.03 ±17.42 | 47.70 ±23.54 | 23.43 | 0.000 *** |
| | | Random Forest | 44.08 ±8.27 | 39.50 ±16.09 | 7.75 | 0.012 * |
| | | LDA | 37.38 ±20.18 | 40.68 ±32.41 | 1.86 | 0.189 |
| | | KNN | 45.76 ±39.01 | 25.98 ±28.25 | 52.34 | 0.009 ** |
| Subject E | | Poly | 57.62 ±5.12 | 43.91 ±12.23 | 97.50 | 0.000 *** |
| | | RBF | 60.69 ±3.66 | 45.14 ±8.92 | 172.9 | 0.000 *** |
| | | Random Forest | 44.01 ±2.86 | 39.60 ±28.47 | 5.59 | 0.029 * |
| | | LDA | 54.46 ±1.73 | 38.10 ±11.70 | 179.20 | 0.000 *** |
| | | KNN | 47.91 ±11.30 | 29.09 ±5.95 | 184.70 | 0.000 *** |
| Subject F | | Poly | 41.26 ±33.40 | 32.48 ±23.36 | 12.21 | 0.002 ** |
| | | RBF | 43.24 ±15.11 | 32.07 ±26.91 | 26.73 | 0.000 *** |
| | | Random Forest | 40.62 ±41.21 | 31.22 ±21.83 | 12.61 | 0.002 ** |
| | | LDA | 28.20 ±11.10 | 28.41 ±15.10 | 0.014 | 0.905 |
| | | KNN | 35.72 ±22.94 | 21.51 ±17.76 | 44.60 | 0.000 *** |
| Subject G | | Poly | 39.99 ±3.67 | 28.70 ±7.71 | 100.74 | 0.000 *** |
| | | RBF | 41.84 ±8.73 | 28.54 ±6.27 | 106.05 | 0.000 *** |
| | | Random Forest | 36.30 ±13.30 | 30.01 ±12.82 | 13.59 | 0.001 ** |
| | | LDA | 32.23 ±15.20 | 26.69 ±9.66 | 11.10 | 0.003 ** |
| | | KNN | 32.08 ±8.63 | 24.43 ±2.64 | 46.70 | 0.000 *** |
| Subject H | | Poly | 24.58 ±6.22 | 23.71 ±6.86 | 0.52 | 0.470 |
| | | RBF | 22.08 ±19.00 | 22.41 ±9.19 | 0.034 | 0.854 |
| | | Random Forest | 22.41 ±24.03 | 20.78 ±17.83 | 0.57 | 0.450 |
| | | LDA | 22.08 ±15.07 | 23.83 ±9.53 | 1.11 | 0.304 |
| | | KNN | 20.88 ±11.39 | 18.27 ±10.27 | 2.83 | 0.109 |
| Subject I | | Poly | 35.94 ±8.58 | 28.92 ±18.11 | 16.61 | 0.000 *** |
| | | RBF | 41.59 ±13.09 | 30.03 ±10.63 | 50.71 | 0.000 *** |
| | | Random Forest | 37.35 ±12.56 | 31.44 ±14.59 | 11.58 | 0.003 ** |
| | | LDA | 35.47 ±12.77 | 29.56 ±6.55 | 16.26 | 0.000 *** |
| | | KNN | 35.47 ±7.40 | 26.78 ±20.00 | 24.78 | 0.000 *** |

Table 14S, LDA mean accuracies for before PCA and after PCA scenario in time domain. P_Values calculated with one-way ANOVA test. Accuracies are mean accuracies of 10-fold cross validation ± var.

| | Before PCA | After PCA | Statistic | P-Value |
|---|---|---|---|---|
| Subject B | 36.44<br>±14.14 | 46.34<br>±10.76 | 35.37 | 0.000<br>*** |
| Subject C | 37.38<br>±20.18 | 60.58<br>±8.69 | 167.66 | 0.000<br>*** |
| Subject E | 54.46<br>±1.73 | 60.37<br>±6.80 | 36.80 | 0.000<br>*** |
| Subject F | 28.20<br>±11.10 | 42.79<br>±17.52 | 63.77 | 0.000<br>*** |
| Subject G | 33.13<br>±6.05 | 40.63<br>±5.40 | 44.12 | 0.000<br>*** |
| Subject H | 22.08<br>±15.07 | 25.46<br>±34.24 | 2.09 | 0.165 |
| Subject I | 35.47<br>±12.77 | 42.69<br>±18.07 | 15.22 | 0.001<br>** |

Table 15S, LDA mean accuracies for before PCA and after PCA scenario in time-frequency domain (Morlet). P_Values calculated with one-way ANOVA test. Accuracies are mean accuracies of 10-fold cross validation ± var.

| | Before PCA | After PCA | Statistic | P-Value |
|---|---|---|---|---|
| Subject B | 21.26<br>±18.70 | 27.08<br>±6.09 | 12.30 | 0.002<br>** |
| Subject C | 24.91<br>±16.44 | 37.58<br>±14.01 | 47.40 | 0.000<br>*** |
| Subject E | 26.28<br>±2.92 | 41.50<br>±6.61 | 218.50 | 0.000<br>*** |
| Subject F | 24.34<br>±20.32 | 27.59<br>±35.97 | 1.68 | 0.210 |
| Subject G | 21.68<br>±6.12 | 30.49<br>±7.67 | 53.19 | 0.000<br>*** |
| Subject H | 19.69<br>±13.66 | 21.97<br>±29.05 | 1.097 | 0.308 |
| Subject I | 22.60<br>±4.06 | 29.61<br>±6.93 | 40.17 | 0.000<br>*** |

Table 16S, LDA mean accuracies for before PCA and after PCA scenario in time-frequency domain (Wigner-Ville). P_Values calculated with one-way ANOVA test. Accuracies are mean accuracies of 10-fold cross validation ± var.

| | Before PCA | After PCA | Statistic | P-Value |
|---|---|---|---|---|
| Subject B | 28.88<br>±10.14 | 34.86<br>±17.27 | 11.72 | 0.003<br>** |
| Subject C | 40.68<br>±32.41 | 45.57<br>±27.98 | 3.56 | 0.07 |
| Subject E | 38.10<br>±11.70 | 43.13<br>±10.13 | 10.42 | 0.004<br>** |

| | | | | |
|---|---|---|---|---|
| Subject F | 28.41 ±15.10 | 30.71 ±28 | 1.09 | 0.309 |
| Subject G | 26.69 ±9.66 | 31.65 ±14.92 | 9.01 | 0.007 ** |
| Subject H | 23.78 ±10.06 | 19.36 ±15.96 | 6.74 | 0.018 * |
| Subject I | 29.56 ±6.55 | 32.17 ±18.89 | 2.41 | 0.137 |

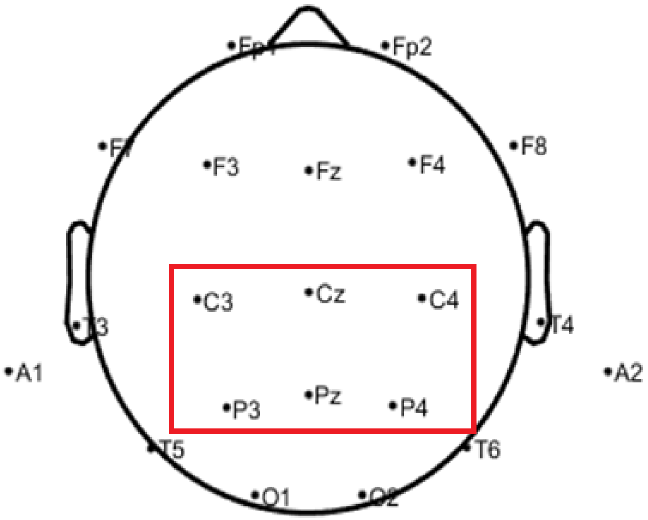

Figure-1S. Electrode positions. The common 6 EEG channels used in the present study.

3. Statistical Tests for Comparing Classification Algorithms | by Tiago Toledo Jr. | Towards Data Science. https://towardsdatascience.com/statistical-tests-for-comparing-classification-algorithms-ac1804e79bb7.

4. Home - mlxtend. http://rasbt.github.io/mlxtend/.
